# Human jejunal enteroids for studies of epithelial drug transport and metabolism

**DOI:** 10.1101/2025.03.17.643545

**Authors:** Merve Ceylan, Foteini Tzioufa, Maria Letizia Di Martino, Rebekkah Hammar, Ana C. C. Lopes, Jens Eriksson, Dinh Son Vo, Magnus Sundbom, Martin Skogar, Per M. Hellström, Dominic-Luc Webb, Maria Karlgren, Iain Gardner, Patrik Lundquist, Daisy Hjelmqvist, Mikael E. Sellin, Madlen Hubert, Per Artursson

## Abstract

Intestinal enteroids are stem cell-based “mini-guts” that mimic many aspects of the corresponding epithelial barrier *in vivo*. Here, we established and characterized differentiated apical-out (AO) and basal-out (BO) jejunal enteroids in suspension and followed their differentiation by quantitative global proteomics and different microscopic techniques. The barrier integrity and function and subcellular location of nutrient and clinically important drug transporters were investigated in the matured enteroids using live-cell microscopy. The presystemic metabolism of two drugs by CYP3A4 was determined and the results were used to predict the pharmacokinetics after oral administration by a PBPK population model. The differentiated AO enteroids displayed a protein profile that overlapped both qualitatively and quantitatively with that of freshly isolated jejunal enterocytes and tissue. They exhibited a morphology that recapitulates the mature villus enterocyte *in vivo*, formed an intact barrier with a well-developed glycocalyx and are impermeable to the hydrophilic low molecular weight compound lucifer yellow and transported a medium chain fatty acid derivative by FATP4 into lipid deposits. The clinically important ABC-transporters Pgp and BCRP were expressed at near *in vivo* levels, had the correct subcellular localization and effluxed their substrates. Terfenadine and midazolam were metabolized by CYP3A4 and the results were used to predict the clinical pharmacokinetics of the drugs after oral administration with good accuracy. We conclude that suspended 3D AO enteroids provide a physiologically relevant model for studies of intestinal function that offers convenient access to the apical surface and is easy to dispense in multi-wells formats for large scale experimentation.

## Introduction

The single layer of epithelial cells lining the inner lumen of the human intestine provides a barrier against ingested noxious compounds and microorganisms and at the same time assures absorption of nutrients and other essential food constituents such as vitamins and trace elements. Most nutrients are absorbed by mature enterocytes at the villus tips in the upper part of the small intestine, the jejunum. This is also the major absorption site for orally administered drugs and it is thus not surprising that the selective barrier function of the jejunal epithelium is of relevance also in drug discovery and development.^1^ *In vitro* models of the human jejunum are therefore of great interest, but no model has so far encompassed all the features of the absorptive villus enterocyte *in vivo*.

Absorptive enterocytes originate from stem cells in the crypts and differentiate during their journey along the crypt-villus axis to fully differentiated enterocytes at the tips of the villi, at which point they have developed the features required to perform their essential dual functions for a short period before they are being shed off and replaced by new cells. The lifespan of human enterocytes is not more than 3-5 days^2^ and they have therefore been notoriously difficult to maintain for functional studies in cell culture. Freshly isolated villus cells in suspension maintain *in vivo*-like features such as metabolic capacity for a few hours but their functions and viability decline after isolation. In absence of primary cell models, the Caco-2 cell line, established from a colorectal carcinoma, has been useful in mechanistic studies ranging from transport mechanisms of essential food constituents and drugs, to barrier dynamics, *e.g.,* after exposure to enteropathogens.^3,4^ Caco-2 cells have a phenotype of villus enterocytes, form a tight junction barrier and express functional nutrient transporters. Due to their cancerous origin, Caco-2 cells however also have certain abnormalities, including truncated glycosylation of surface-bound mucins which reduces their physical glycocalyx barrier to infectious microorganisms and nanoparticles as compared to *in vivo.*^5,6^ Caco-2 cells also have aberrant expression of key proteins, exemplified by a metabolic enzyme panel that is partly of fetal origin.^7^

Organoids of different origins represent a major advance throughout the entire field of biomedicine^8–10^ and organoids from adult human tissue stem cells are now an important tool for studies of normal and pathological conditions of the mucosal barrier. Intestinal epithelial organoids originating from adult stem cells, named enteroids when from the small intestine, preserve important functions of the intestinal epithelium^11^ and can be grown in different configurations. These include 2D monolayers^12^, 3D enteroids entrapped in an extracellular matrix as originally described^13,14^ and 3D enteroid suspensions, each with their advantages and disadvantages.^15^ A drawback with the original 3D basal-out (BO) configuration is that the apical surface is facing an inner confined lumen, that can only be accessed by perforation of the epithelial barrier *e.g.*, by microinjection. This can be addressed by dispersing the enteroids and reseeding them as 2D monolayers.^12^ This configuration has been used to study nutrient, drug transport and, metabolism function^16–19^, as well as microbial infection.^20^ Some of us recently developed protocols for expansion and differentiation of jejunal enteroids into 2D monolayers and observed that optimally differentiated monolayers provided improved protection against *Salmonella* infection.^20^ The 2D monolayers have barrier features of the human absorptive villus cells, with the expected morphology comprising a dense carpet of microvilli and well developed glycocalyx.^20^ Further, quantitative global proteomics indicated that many proteins of the glycocalyx, surface-bound mucins, and tight and adherent junctions were present in quantities comparable to, or approaching, those found in human jejunum *in vivo*. However, many membrane transporters and metabolizing enzymes had markedly lower expression levels than observed for native epithelial tissue.

As the intestinal epithelium senses curvature, e.g. crypt-like regions of enteroids preferentially localize in indented areas of a supporting scaffold, while differentiated villus cells cover the finger-like projections.^21^, we reasoned that a curvature mimicking the latter would promote the development of fully differentiated absorptive enterocytes. Morphogenically everted 3D apical-out (AO) enteroids as originally described by Co *et al*.^15^ have the convex curvature of villus tips, and we therefore set out to characterize and investigate these enteroids for functional studies of nutrient and in particular drug transport and metabolism, using primary jejunal enterocytes and tissue as references. We first established conditions for optimal growth and differentiation of enteroids derived from human intestinal jejunum, using quantitative global proteomics and compared the result with BO enteroids, jejunal enterocytes and tissues, partly isolated from the same individuals (Table S1). We then developed live-cell microscopy techniques to characterize the barrier function and important transport functions in the enteroid population. Finally, we investigated the metabolism of two orally administered drugs (terfenadine and midazolam), normalized the metabolism against enzyme expression and predicted the clinical pharmacokinetics of the two drugs using physiologically based pharmacokinetic population models. Our results show that optimally differentiated AO enteroids have a phenotype close to fully differentiated absorptive villus enterocytes, express key proteins at levels comparable to those in jejunal tissue and are fully functional with regard to uptake of fatty acid, efflux via P-glycoprotein and drug metabolism by cytochrome P450. The jejunal enteroids hence provide a physiologically relevant model for studies of intestinal function in health and disease as well as in studies of intestinal drug transport and metabolism.

## Material and Methods

### Human jejunum tissue

Samples from human jejunum were collected from obese subjects undergoing Roux-en-Y gastric bypass surgery at the Uppsala University Hospital, Uppsala, Sweden. Subjects followed a low-calorie diet for three weeks prior to surgery and did not suffer from inflammatory or infectious bowel diseases at the time of surgery. Subjects diagnosed with type I or II diabetes were excluded from the study. Samples were taken 60 cm from the ligament of Treitz. Group level demographic information on the donor tissue samples is summarized in Table S1. All donors gave written informed consent. Samples were pseudonymized before proceeding with further analysis to protect patient identities and personal details. The study was reviewed and approved by local governing body (Etikprövningsmyndigheten, Sweden, permit number 2010-157 with addendum 2010-157-1 and 2020-05754, and permit number 2023-01525-01). The tissue samples were either snap frozen in liquid nitrogen and stored at −150°C, pending global proteomics analysis, or used immediately for isolation of primary jejunal enterocytes or adult jejunal stem cell-containing crypts for enteroid culturing.

### Primary enterocyte isolation from jejunal tissue

Intestinal samples (2 - 4 cm long pieces of jejunum) were collected in 50 mL tubes with oxygenated, ice-cold, calcium- and magnesium-free Krebs-Henseleit buffer (Sigma-Aldrich, St. Louis, MO, USA). Samples were immediately transported to the laboratory. The jejunal samples were opened along the mesentery and the mucosa was isolated along the mucosa muscularis. The dissected mucosa was carefully washed three times in cold Krebs-Henseleit buffer and cut into small pieces (approximately 0.5 - 1 cm^2^). The jejunal primary enterocytes were dissociated from jejunal tissue samples by incubating 0.5 - 1 cm^2^ pieces of tissue in Krebs-Henseleit buffer supplemented with dithiothreitol (DTT, 1 mM, Sigma-Aldrich, St. Louis, MO, USA) and ethylenediaminetetraacetic acid (EDTA, 3 mM, Sigma Aldrich). Briefly, tissue samples amounting to approximately 1 g were first shaken on ice for 30 min on a nutating mixer (VWR, Stockholm, Sweden), and then shaken at 37°C for 60 min. Mucus, larger tissue pieces, cell aggregates, and large debris were removed from the supernatant by filtering through a cell strainer (70 µm pore size, mini cell strainer, Nordic Biosite AB, Täby, Sweden). To enrich the epithelial cells and remove blood cells, cell suspensions were centrifuged in Percoll® (Sigma-Aldrich) density gradients [20%, 40% Percoll in Krebs-Henseleit buffer with 2% bovine serum albumin (BSA, Sigma-Aldrich)] at 10,000 g for 10 min, followed by washing with Krebs-Henseleit buffer with 2% BSA. The enriched cells were washed twice with 10 mL Krebs-Henseleit and resuspended in ice cold Dulbecco′s Modified Eagle′s Medium (DMEM, Gibco, Thermo Fisher Scientific, Waltham, MA, USA). Viability was determined to be > 85% by acridine orange and propidium iodide (AOPI) method in a Nexcelom Cellometer Vision (Nexcelom Bioscience, Lawrence, MA, USA).

### Adult jejunal epithelial stem cell isolation

Human jejunal adult stem cell lines were established as described previously.^22,23^ Briefly, freshly isolated jejunal tissue was transferred to ice-cold phosphate buffered saline (PBS, Thermo Fisher Scientific) and placed on a Styrofoam cushion. PBS was used to remove particles, and the muscle layer was removed with a scissor. A ∼6 mm tissue piece was taken from the mucosa, rinsed with PBS, minced using surgical scissors, and passed through a 1 mL pipette tip. After centrifugation at 300 g for 5 min and washing with cold PBS, the minced tissue was incubated in Gentle Cell Dissociation Reagent (STEMCELL technologies, Vancouver, BC, Canada) under gentle shaking on ice for 30 min. After a second centrifugation (300 g, 5 min) and resuspension in cold DMEM F12 (Gibco, Thermo Fisher Scientific) with 0.25% BSA, epithelial crypts were extracted. After passing the suspension through a 70 μm cell strainer, the crypt concentration was quantified. To achieve a density of 250 - 750 crypts per dome, crypts were centrifuged at 300 g for 10 min, resuspended in DMEM F12 containing 75% Matrigel (Corning, Corning, NY, USA), and seeded in 50 μL domes in multiwell plates (Sarstedt AG & Co. KG, Nümbrecht, Germany). After 10 min of solidification at 37°C, growth medium (OGM, IntestiCult Human organoid growth medium, STEMCELL technologies) supplemented with 10 μM ROCK Inhibitor Y-27632 (Stemcell Technologies) and 100 U/mL penicillin-streptomycin (PenStrep, Gibco, Thermo Fisher Scientific) was added. After two days, ROCK Inhibitor Y-27632 was removed from cultures. From then on, the medium was replaced every three to four days. Following establishment, enteroids were expanded by continuous enteroid subculturing at day seven. At passage two, newly formed enteroids were frozen in DMEM F12 supplemented with 10% fetal bovine serum (FBS) and 10% dimethyl sulfoxide (DMSO) and cryopreserved in liquid nitrogen.

### Enteroid expansion and differentiation

Cryopreserved enteroids were thawed and resuspended in a cold mixture containing 75% Matrigel and 25% OGM. Three 15 - 20 μL domes of the suspended cells were added to each well in a 24-well plate. The domes were solidified for 10 min at 37°C, and then overlaid with 500 μL of OGM containing 100 U/mL PenStrep and 10 μM of ROCK inhibitor Y-27632 for two days. Afterwards OGM was replenished excluding the Y-27632 inhibitor but containing 100 U/mL PenStrep. The medium was replaced with fresh OGM medium every two to three days and cultures were maintained at 37°C in 5% CO_2_ until the enteroids were large enough for expansion. Thereafter, the enteroids were passaged every seven to ten days. Briefly, the Matrigel domes were dispersed through repeated gentle pipetting with 1 mL of Gentle Cell Dissociation Reagent per well. The enteroids were then centrifuged at 300 g for 5 min at 4°C. The pelleted enteroid fragments were washed with 1 mL per well of DMEM F12 supplemented with 0.25% BSA and then centrifuged a second time at 300 g for 5 min at 4°C, diluted between 1:3 and 1:8 times and then re-embedded in 15-20 μL Matrigel domes for further expansion as described above. Passages 4 to 20 were used for all experiments.

The enteroid differentiation was initiated at day 3 post splitting. The OGM was removed and 500 μL of cold Cell Recovery Solution (Corning, NY, USA) was added. The Matrigel domes were broken by gently pipetting up and down three times with a 1 mL pipette tip. Enteroids were transferred to a 15 mL Falcon tube and incubated on ice for 1 h on a nutating table (20 rpm, FisherBrand, Pittsburg PA, USA). Supernatants were carefully removed and the enteroid pellets were washed with 3 mL of ice-cold DMEM F12 + 0.25% BSA. After removing the supernatants, the pellets were re-suspended in organoid differentiation medium (ODM, STEMCELL technologies) without DAPT. The enteroid suspensions (500 μL per well) were transferred to pre-cooled 24-well ultra-low attachment plates (Sarstedt AG & Co. KG). Matrigel was added at 7.5% to the medium to maintain the natural basal-out configuration of the enteroids in the suspension cultures. To reverse the configuration of the enteroids in suspension cultures to apical-out, the Matrigel was omitted as previously described.^15^ The same procedure was used to remove Matrigel from the basal-out cultures immediately before the experiments.

All suspension cultures were maintained at 37°C in 5% CO_2_, and the ODM medium was replaced every second day. To simplify the description of the different growth conditions leading to different types of enteroids, the nomenclature has been simplified as in Table 1, where the first letter indicates enteroids grown in Matrigel (M), basal-out (B) or apical-out (A) configuration, the second letter indicates the growth medium (G) or differentiation medium (D), and the number four or six indicates the number of days the enteroids were cultured before use.

**Table 1.**
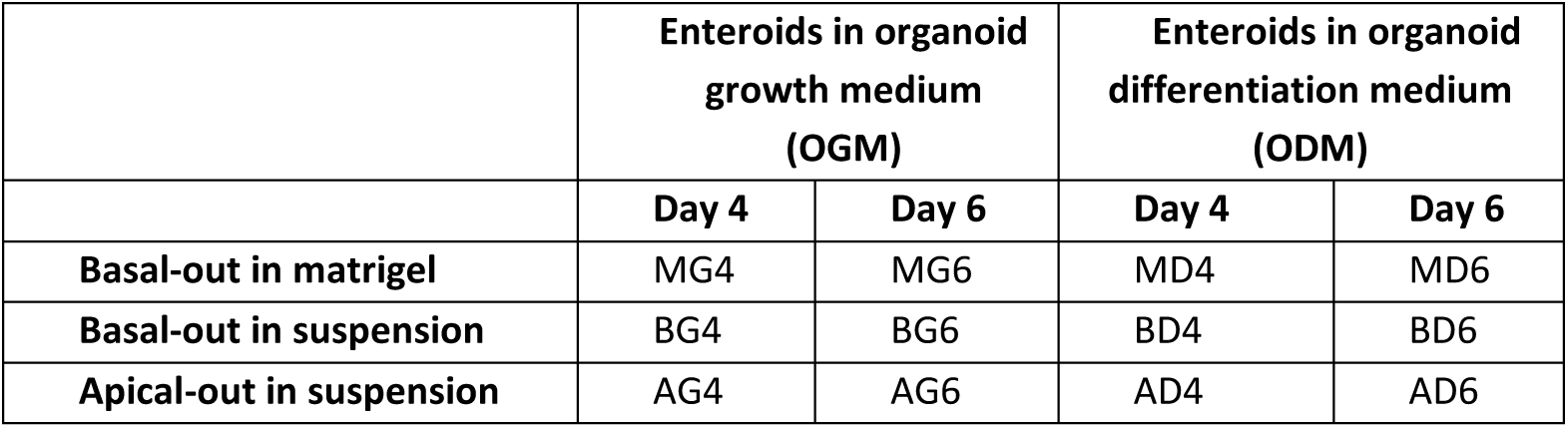
Nomenclature of enteroids in different culture conditions.

|  | Enteroids in organoid growth medium (OGM) |  | Enteroids in organoid differentiation medium (ODM) |  |
| --- | --- | --- | --- | --- |
|  | Day 4 | Day 6 | Day 4 | Day 6 |
| **Basal-out in matrigel** | MG4 | MG6 | MD4 | MD6 |
| **Basal-out in suspension** | BG4 | BG6 | BD4 | BD6 |
| **Apical-out in suspension** | AG4 | AG6 | AD4 | AD6 |

### Global proteomics

Jejunal tissues (n = 5, from different donors), enterocytes (n = 4, from different donors) and all enteroid formats listed in Table 1 were lysed in lysis buffer (100 mM Tris/HCl, Sigma-Aldrich) containing 50 mM DTT and 2% (w/v) sodium dodecyl sulfate (SDS, Sigma-Aldrich) with a pH of 7.8. The samples were then heated (95°C, 5 min) and sonicated (20 pulses of 1s, 20 % amplitude with Vibra-Cell ultrasonic processor (Sonics & Materials Inc., Connecticut, US). The multienzyme digestion filter aided sample preparation (FASP) method was performed using trypsin (Nordic Biolabs AB, Täby, Sweden) and Lys-C peptidase (Thermo Fisher Scientific) to digest proteins to peptides.^24^ An amount of 10 μg of peptide mixture was dried using SpeedVac (Thermo Fisher Scientific) and stored at -15°C. Before the proteomics run, 10 µg of the resulting peptide mixture was dissolved in 10 μL mobile phase A [0.1% formic acid (VWR) in Milli-Q water (Milli-Q lab water purification system, Merck, Darmstadt, Germany)]. Proteomic analysis was performed on an Orbitrap Q Exactive HF mass spectrometer (MS, Thermo Fisher Scientific) which was coupled to a nano–liquid chromatography (nLC) and MS was set to a Top-N method (Data Dependent Acquisition). An EASY-spray C18-column (50 cm, 75 µm inner diameter, Thermo Fisher Scientific) was used to separate peptides in acetonitrile/water gradient (0.1% formic acid) at 300 nL/min flow rate. Proteins were identified using MaxQuant software^25^ and total protein approach (TPA) was used as the quantification method.^26^

### Brightfield microscopy

Enteroid development was monitored by brightfield microscopy, both in Matrigel domes and in suspension cultures. Prior to microscopy, the suspension cultures were strained in 40 µm pore size cell strainers (Funakoshi, Tokyo, Japan), washed three times with ice-cold DMEM F12 to remove cellular debris and excess Matrigel and resuspended in DMEM F12 in an ultra-low attachment plate. Brightfield imaging was performed in 4x, 10x and 20x magnifications using a Nikon Eclipse TS100 Inverted Routine Microscope (Nikon Corporation, Tokyo, Japan). Enteroid size (cross sectional area, Feret’s diameter) and shape descriptors (shape factor) for both apical-out (AD6; n = 127) and basal-out (BD6; n = 94) were approximated using images in 4x magnification. Individual enteroids were segmented and masked using Fiji ImageJ (version 1.54j).^27^ Shape descriptors were acquired by measuring the masks of the segmented enteroids.

### Scanning electron microscopy

Enteroids were fixed at 4°C overnight with 2.5% glutaraldehyde (TAAB Laboratories, Aldermaston, England) in 0.1 M PHEM-buffer [60 mM piperazine-N,N’-bis(2-ethanesulfonic acid) (Sigma Aldrich), 25 mM HEPES (4-(2-hydroxyethyl)-1-piperazineethanesulfonic acid (Sigma-Aldrich), 10 mM egtazic acid (Merck), and 4 mM MgCl_2_ x 6H20 (Sigma-Aldrich), pH 6.9]. Samples were allowed to adhere to poly-D-lysine coated cover slips (Ted Pella, Redding, CA, USA) overnight. Cover slips were washed and dehydrated with a graded series of ethanol using the Biowave pro+ microwave (Pelco, Ted Pella) and subsequently critically point dried (Leica EM300). The cover slips were then mounted with carbon tape on an aluminum stub and coated with 5 nm of platinum (Quorum Q150T ES, Quorum Technologies, Lewes, UK). The sample morphology was analyzed by field-emission scanning electron microscopy (FESEM; Carl Zeiss Merlin) using the Inlens secondary electron detector at an accelerating voltage of 5 kV and probe current of 120 pA.

### Transmission electron microscopy

Enteroids were fixed with 2.5% glutaraldehyde (TAAB Laboratories) in 0.1 M sodium cacodylate (Sigma-Aldrich) or PHEM-buffer. Samples were washed in buffer and post stained in 0.8% potassium ferricyanide (Sigma-Aldrich), 1% OsO4 (TAAB Laboratories) in buffer, further stained with 1% tannic acid (EMS, Hatfield, PA) in Milli-Q water and 1% uranyl acetate (TAAB Laboratories) and further dehydrated using a graded series of ethanol. Samples were infiltrated using a graded series of Spurr’s resin (TAAB Laboratories) and finally polymerized overnight in an oven at 65°C. All sample preparation except fixation was done using the Biowave Pro+ processing microwave (Pelco). Ultrathin sections of 70 nm were picked up on formvar coated grids and analyzed using a Talos L120C (FEI, Eindhoven, The Netherlands) operating at 120 kV. Micrographs were acquired with a Ceta 16M CCD camera (FEI, Eindhoven, The Netherlands) using Velox software (version 2.14.1.40, Thermo Fischer, Eindhoven, The Netherlands).

### Immunofluorescence staining

The enteroids were first fixed for 30 min at room temperature with a paraformaldehyde solution at pH 7.4, consisting of 2% paraformaldehyde (PFA, Thermo Fisher Scientific), 60 mM dibasic sodium phosphate (Sigma-Aldrich), and 14 mM monobasic sodium phosphate (Sigma-Aldrich). Following fixation, enteroids were washed three times with Dulbecco’s Phosphate-Buffered Saline (DPBS, Gibco, Thermo Fisher Scientific) for 10 minutes each. Enteroids were then permeabilized with 0.1% saponin (Thermo Fisher Scientific) and blocked with 2% donkey serum (Merck) in PBS, each for 30 minutes at room temperature.

Primary antibodies used included anti-zonula occludens-1 (ZO-1, 1:100, mouse monoclonal, Thermo Fisher Scientific), anti-CYP3A4 (1:200, mouse monoclonal, Thermo Fisher Scientific), anti-ABCG2/BCRP (1:100, rabbit monoclonal, Abcam), anti-ABCB1/Pgp (1:100, mouse monoclonal, Thermo Fisher Scientific), and anti-SLC27A4/FATP4 (1:100, rabbit monoclonal, Thermo Fisher Scientific). These antibodies were prepared in a blocking/permeabilization buffer containing 3% donkey serum, 1% saponin, and 1% Triton X-100 (Sigma-Aldrich) at the specified concentrations. Enteroids were resuspended in 150 µL of the primary antibody staining solution and incubated overnight at 4°C in anti-adherence solution (STEMCELL technologies, Vancouver, BC, Canada) pre-treated microcentrifuge tubes (Eppendorf, Hamburg, Germany). Following incubation, the enteroids were washed three times with PBS for 10 minutes each.

To detect the proteins of interest, fluorescence-labelled secondary antibodies were used: Alexa Fluor 488 (1:500, anti-mouse, Abcam), Alexa Fluor 488 (1:500, anti-rabbit, Thermo Fisher Scientific), and Alexa Fluor 568 (4 µg/mL, anti-rabbit, Thermo Fisher Scientific). Phalloidin conjugated with Alexa Fluor 660 (1:400, Thermo Fisher Scientific) was also added to stain actin. Enteroids were resuspended in 150 µL of secondary antibody and phalloidin solution and incubated overnight at 4°C. After three washes with PBS, each for 10 minutes, the enteroids were resuspended in 10 µL PBS and transferred to a microscopy glass slide (KEBO-Lab, Stockholm, Sweden). Excess PBS was removed, and VECTASHIELD® Vibrance™ Antifade Mounting Medium with DAPI (BioNordika, Solna, Sweden) was added to the sample. A coverslip, with vacuum grease applied at the corners as spacers, was placed on top of the slide. The enteroids were imaged using a Zeiss LSM800 confocal microscope equipped with a 20x objective and a 40x immersion oil objective.

### Live-cell imaging of enteroids in suspension

Live-cell imaging was performed on a custom-built microscope based on an Eclipse Ti2 body (Nikon), using a 10X/0.45 Plan Apo Lambda air objective (Nikon), and a back-lit sCMOS camera with a physical pixel size of 11 μm (Prime 95B, Photometrics), and a final image pixel size of 1.12 µm.^23^ The imaging chamber was maintained at 37°C, 5% CO_2_ in a moisturized atmosphere. Bright-field images were acquired by differential interference contrast (DIC), and fluorescence imaging by the excitation light engine Spectra-X (Lumencor) and emission collection through quadruple band pass filters (89402 & 89403; Chroma) and a spinning disk module (X-light V2, Crest optics).

### Lucifer yellow barrier integrity assay in live enteroids

Enteroids were washed three times with DMEM F12 and resuspended in DMEM F12 containing 100 μM of Lucifer yellow (LY) dilithium salt (Ex/Em 428/536 nm) fluorescent dye (VWR). The enteroids were incubated at a 37°C for 30 minutes with gentle shaking at 100 rpm (dye loading). The apical-out enteroid suspension was kept in the dye solution throughout the experiment, while the basal-out enteroid suspension was washed three times with DMEM F12 before analysis. Samples were observed 30, 60 and 90 minutes after addition of the dye at 37°C and 5% CO_2_ using live-cell time-lapse microscopy. In the control experiments, the enteroids were permeabilized by addition of EDTA in a final concentration of 2 mM at 30 minutes post dye loading. Quantification of LY fluorescence was performed using Fiji ImageJ (version 1.54j). In apical-out enteroids, quantification was performed for all enteroids imaged. On the other hand, for basal-out, only terminally differentiated enteroids were pre-selected for analysis, namely enteroids which exhibited crypt-like structures (budding), a defined lumen and a thick epithelial layer. The pre-selection for basal-out enteroids was made strictly on the brightfield channel. Integrity was determined by measuring the mean gray value of enteroids within a specific region of interest (ROI) compared to the mean gray value of the surrounding background (same sized ROI). Data were analyzed with one-way ANOVA Kruskal-Wallis test and multiple comparison analysis. An average of 208 ± 66 observations were made per timepoint.

### Fatty acid uptake assay using live-cell microscopy

Enteroids were washed trice with warm DMEM F12, resuspended in DMEM F12 containing 2.5 μM C1-BODIPY-C12 (Ex/Em 500/510 nm, Thermo Fisher Scientific) with 5 μM BSA and incubated for 10 min at 37°C.^15^ Then, the enteroids were washed and resuspended in DMEM F12 with HCS LipidTOX™ Deep Red Neutral Lipid Stain (1:5,000, Ex/Em 637/655 nm, Thermo Fisher Scientific).^28^ Live-cell imaging was performed at 60x magnification using the custom-built microscope described above (37°C, 5% CO_2_). Enteroids were imaged at 60x to visualize accumulation of C1-BODIPY-C12 within lipid droplets.

### Analysis of P-glycoprotein function using rhodamine 123 in live enteroids

Enteroids were strained and incubated in DMEM F12 with 10 μM of the P-glycoprotein (Pgp) fluorescent substrate rhodamine 123 (Rho123, Ex/Em 508/528 nm, Sigma-Aldrich) in a shaking water bath at 100 rpm for 1 hour in 37°C in dark conditions. In control experiments, the enteroids were pre-treated with 2 μM of Pgp inhibitor elacridar (ELC, Sigma-Aldrich) for 1 hour at 37 °C, and subsequently treated with 10 μM of Rho123 in presence of 2 μM ELC in DMEM F12 for 1 hour at 37°C. After incubation, the enteroids were washed three times at 37°C, resuspended in warm DMEM F12 and subsequently live imaged at 4x and 60x magnifications using the custom-built microscope described above. Dye localization in each organoid was determined in 60x magnification images, using the Line Profile function of Fiji ImageJ: a line was drawn on the brightfield channel across each enteroid’s diameter and the mean gray value in the fluorescent channel was measured along the length of that line. On a population scale, fluorescence was determined by measuring mean grey values in 4x magnification images, using an in-house developed algorithm for Fiji ImageJ. Similarly, to the barrier integrity assay, the whole AO population was imaged, whereas in basal-out enteroids a preselection was made at the brightfield channel. Only basal-out enteroids which exhibited a differentiated phenotype with budding were analysed. Moreover, the only basal-out enteroids selected for analysis had the focal plane strictly on their fully defined lumen. For apical-out enteroids, which have the Pgp transporters facing outwards, the whole organoid fluorescence was compared to that of the background fluorescence. For basal-out enteroids, which in contrast to apical out enteroids have a clearly distinguishable lumen, the luminal fluorescence of each enteroid was compared to the fluorescence of the surrounding epithelium. Data analysis was performed using GraphPad Prism (version 10). For apical-out enteroids the mean gray value of the Rho123 exposed enteroids (n = 102) was compared to the control group; Rho123 exposed enteroids pre-incubated with the specific Pgp inhibitor ELC (n = 143). Significant differences between the two groups were investigated using Mann-Whitney unpaired t-test. For basal-out enteroids, Wilcoxon’s paired t-test was used for both Rho123 treated (n = 154) and Rho123 + ELC treated (n = 102) populations, since each enteroid’s luminal gray value was compared to its epithelial gray value.

### Measurement of BCRP efflux transporter activity using pheophorbide A

BCRP activity was measured using the BCRP substrate pheophorbide A (PhA) as previously described.^29^ In brief, apical-out enteroids (∼ 3 × 10^5^ cells) were incubated with 1 μM PhA, with or without 2 μM of the BCRP inhibitor KO143. After 18 h incubation, enteroids were washed five times with PBS containing Ca^2+/^Mg^2+.^ Thereafter, relative fluorescence was measured using a Tecan plate reader (Tecan, Grödig, Austria) using excitation and emission wavelengths of 395 nm and 670 nm, respectively. Results were compared to negative (MDCK cells without BCRP) and positive (MDCK cells overexpressing BCRP) controls as described in Wegler *et al*.^29^

### CYP3A4 metabolic activity

Enteroids were resuspended at a density corresponding to 3 × 10^6^ enterocytes per well (NucleoCounter, chemometec, Alleröd, Denmark) and incubated with 1 μM of either of the CYP3A4 substrates terfenadine or midazolam (Sigma-Aldrich) for 2 h. As negative control, cells were incubated for 5 min at 95°C and cooled on ice before the experiment. Cell suspensions and compound solutions were equilibrated to 37°C prior to experiment start. Samples were collected at 0, 30, 60 and 120 min and lysed with 120 µL 100% acetonitrile containing 5 nM warfarin. Samples were centrifuged at 4°C at 3500 rpm for 20 min. A sample volume of 100 µL was transferred to a new plate and 100 µL of 60% acetonitrile containing 3 nM warfarin was added. The protein content of the pellets was quantified using the tryptophan method.^30^ Compound concentrations were determined by UPLC-MS/MS using an Acquity UPLC coupled to a XEVO TQ triple-quadrupole mass spectrometer with positive electrospray ionization (Waters Corp., Milford, MA). For chromatographic separation, a C18 BEH 1.7 µm column (Waters Corp.) was used, with a general gradient of 1% to 90% of mobile phase B over a total running time of 5 min. Mobile phase A consisted of 5% acetonitrile and 0.1% formic acid in purified water, and mobile phase B of 0.1% formic acid in 100% acetonitrile. The flow rate was set to 0.5 mL/min and 5 µL of the sample was injected. Multiple reaction monitoring methods were optimized for each compound, metabolite and internal standard.

### Physiologically-based pharmacokinetic simulation

Simcyp simulator V23 r2 (Certara Predictive Technologies, Sheffield, United Kingdom) was used for the physiologically-based pharmacokinetic (PBPK) simulation. For simulations using AO enteroid data the intestinal metabolism in the baseline model was replaced with the *in vitro* intrinsic clearance measured in apical out enteroids. The intrinsic clearance (CL_int_) (µL/min/million cells) was converted to a rate of metabolism per mg of microsomal protein by correcting for the measured protein abundance per million cells and with an assumption that the endoplasmic reticulum accounts for 40% of total cellular protein. The *in vitro* CL_int_ was corrected for binding^31^ assuming that the binding in intestinal cells and liver cells is the same. The final CL_int_ used in the model was 177.6 µL/min/mg microsomal protein in the intestine (coefficient of variation = 11%). In the simulation where no intestinal metabolism was considered the CL_int_ in the intestine was set to 0.

For terfenadine a PBPK model was constructed and the input parameters in the model are listed in Table S2. There is no intravenous study published that describes terfenadine metabolism and oral data is scarce with many studies failing to detect parent terfenadine in many dosed subject and only reporting levels of the acidic metabolite fexofenadine.^32–36^ Using more sensitive assay methods has allowed the oral PK of terfenadine to be described but the exposure is generally low (Cmax < 5 ng/ml).^37–40^ Due to these limitations, a fit-for-purpose model was developed. For terfenadine absorption, the advanced dissolution, absorption and metabolism (ADAM) model^41^ was used with an assumption that the drug remains in solution at all times. Permeability was predicted using the mechanistic permeability model.^42^ A minimal PBPK model was employed to describe the distribution of terfenadine with parameters for the peripheral compartment adjusted to recover the concentration time profiles as reported earlier.^38^ Elimination was assigned to be predominantly by CYP3A4 (fraction metabolized: ∼0.98). This value gave terfenadine concentrations in the presence of ketoconazole in line with those reported earlier.^34^ The binding in the intestine [(unbound fraction (fu_gut_)] was set to a value of 0.15 in the final model yielding an intestinal availability (F_g_) of ∼ 0.35 in the baseline model. This value of F_g_ was in line with the value estimated for F_g_ based on the assumption that grapefruit juice inhibits CYP3A in the intestine with minimal effects of hepatic CYP3A4. This value for terfenadine F_g_ is also similar to the value estimated by Gertz *et al* (40%).^43^ The same fu_gut_ model was used in the baseline model and in the model parameterized with metabolism data from the apical enteroids.

For simulation of midazolam concentration – time profiles the midazolam file within the simulator Simcyp simulator V23 r2 (Certara Predictive Technologies) was used without changes for the baseline midazolam simulation. A single oral dose of 2 mg was simulated (10 trials of 15 subjects, 18-55 years, 19% female). The clinical data was extracted from existing literature.^44–47^

### Statistical analysis

For global proteomics, Perseus (version 1.6.2.3)^48^ was used for data analysis (PCA, hierarchical clustering, volcano plot, scatter plot) and enrichment analysis of DEPs (differentially expressed proteins) was performed using various (DAVID^49^, Proteomap, GOrilla) web-based programs. Statistical analysis and figures were made using GraphPad Prism (version 10). Four biological replicates were used for global proteomics and three biological replicates were used in drug transport and metabolism studies.

## Results

### Establishment of optimally differentiated enteroids from human jejunal tissue

For many applications, easy access to the apical surface of the intestinal epithelium is an advantage or even a prerequisite. We therefore established conditions for complete morphogenic eversion of jejunal 3D enteroids from the native basal-out (BO) to apical-out (AO) orientation and followed their differentiation by global proteomics into matured villus-like enteroids (Fig. 1). We compared the differentiation over time (for 4-6 days; below indicated as “4” or “6” in the sample abbreviations) with Matrigel-imbedded and suspended enteroids in the native BO orientation (Fig. 1A). For comparative purposes, we used the same lineage of primary jejunal epithelial cells as in our previous studies in the 2D format.^20^ We also included healthy jejunal villus tissue and freshly isolated jejunal villus enterocytes from this tissue as references.

**Figure 1.**
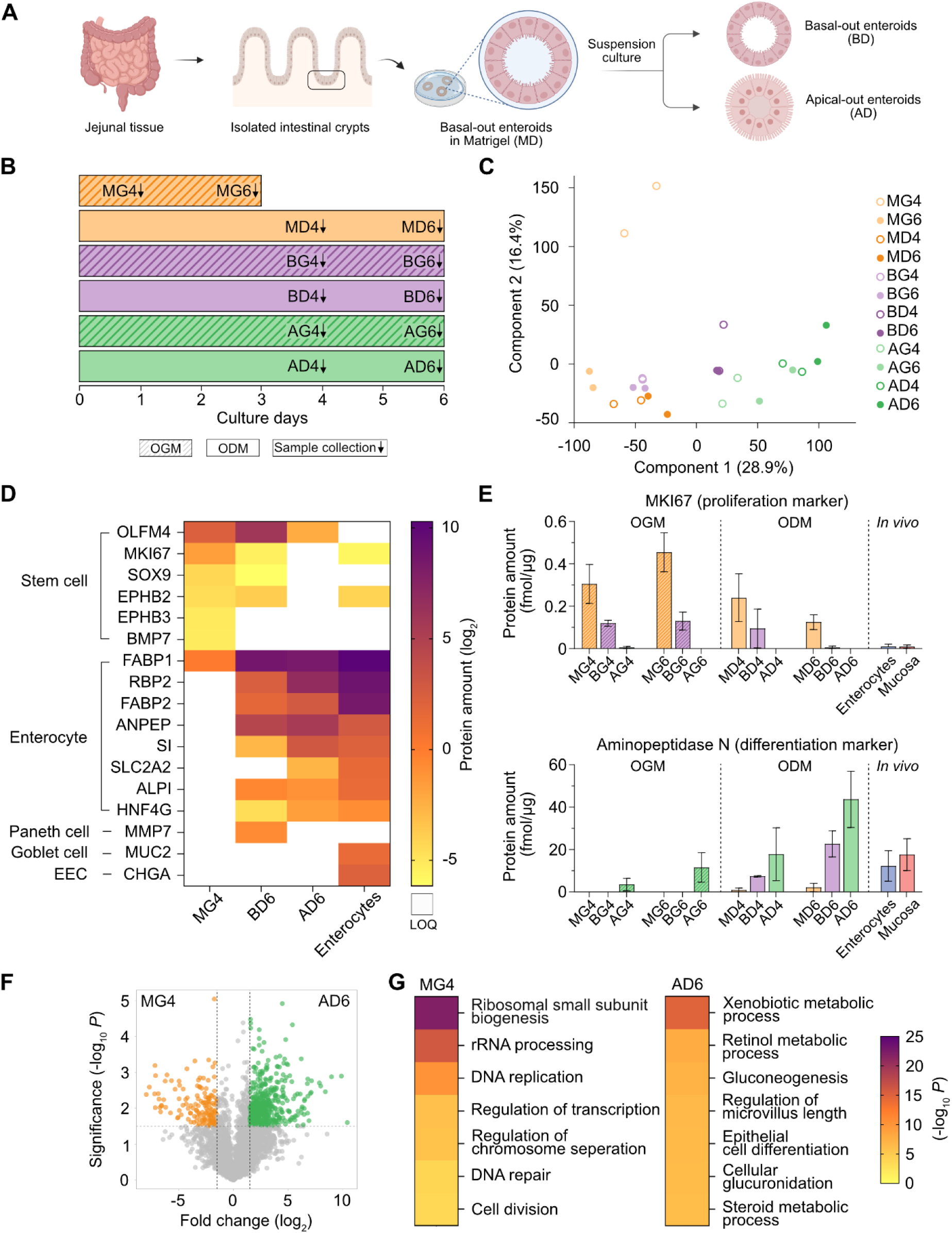
Global protein profiling of human adult stem cell-derived enteroids in different culture formats. **(A)** Schematic of enteroid generation from human tissue samples, followed by culture in Matrigel and suspension formats. **(B)** Schematic of the differentiation protocol. 3D enteroids were initially expanded in Matrigel with OGM for 3 days (not shown on the timeline for simplicity). Enteroids were then cultured in various formats: 3D Matrigel enteroids were either maintained in OGM (MG4, MG6) or differentiated in ODM for 4–6 days (MD4, MD6). Basal-out enteroids in suspension were cultured for 4–6 days in either OGM (BG4, BG6) or ODM (BD4, BD6) with 7.5% Matrigel. Apical-out enteroids in suspension were also cultured in OGM (AG4, AG6) or ODM (AD4, AD6) for 4 – 6 days. **(C)** Principal component analysis (PCA) of protein expression profiles across all enteroid samples, highlighting variations between different culture formats. **(D)** Proteome profiles of 3D basal-out and apical-out enteroids after 6 days in suspension culture. Matrigel-grown enteroids in OGM (MG4) served as a stemlike control, while primary jejunal enterocytes represented differentiated villus enterocytes. BD6 enteroids in ODM retained a low expression of stem cell markers (OLFM4, MKI67, SOX9, EPHB2) and modest expression of enterocyte markers (FABP1, RBP2, ANPEP, GDGAT1). In contrast, AD6 enteroids exhibited a profile and concentration of enterocyte markers (notably FABP1, RBP2, ANPEP, SI, SLC2A2, DGAT1) approaching that of freshly isolated jejunal enterocytes, with minimal or undetectable stem cell marker expression. Both BD6 and AD6 exhibited low or no detectable expression of markers for Paneth cells, goblet cells, and enteroendocrine cells (EECs). LOQ indicates expression levels below the limit of quantification. **(E)** Quantification of KI-67 (proliferation marker, top) and aminopeptidase N (differentiation marker, bottom) across 3D enteroid formats (M: Matrigel, B: basal-out, A: apical-out) cultured in OGM or ODM for 4–6 days, compared to primary enterocyte and jejunal mucosa *in vivo* levels. Data are presented as mean ± SD. **(F)** Volcano plot of differentially expressed proteins between Matrigel-grown enteroids in OGM (day 4, MG4, orange) and apical-out enteroids in ODM (day 6, AD6, green), with corresponding enrichment analysis. Orange and green dots represent proteins significantly upregulated or downregulated (FDR ≤ 0.01, fold change ≥ 1.5) in MG4 (152 proteins) compared to AD6 (501 proteins). **(G)** Heatmap of GO enrichment of biological processes in MG4 and AD6.

Human enteroids were expanded in Matrigel domes with OGM medium^20^ (Fig. 1B) and subsequently maintained in either OGM (below indicated as “G” for growth in the sample abbreviations) or ODM (below indicated as “D” for differentiation in the sample abbreviations), which lacks stemness-promoting factors. For suspension cultures, enteroids were removed from Matrigel and kept suspended for proliferation in OGM and differentiation in ODM. BO enteroids were supported by a low amount of Matrigel in the medium to maintain BO orientation. In contrast, polarity reversal to AO orientation was achieved by suspending the organoids in media without additional Matrigel (Fig 1A-B).^15^ Principal component analysis of proteomes revealed clear differences between the growth conditions, where a time-dependent separation of Matrigel-grown enteroids (“MG” and “MD”), from basal-out (“BG” and “BD”) to apical-out enteroid (“AG” and “AD”) suspensions was evident (Fig. 1C). As expected, MG4 enteroids retained proliferative capacity (MKI67) and expressed markers of an immature phenotype (*e.g.*, OLFM4, SOX9, BMP7) while the expression of most enterocyte markers was below the limit of detection (Fig. 1D). The expression pattern of AO enteroids at day 6 of differentiation (AD6) was close to that of freshly isolated jejunal villus enterocytes in that most proliferation and stem cell markers were below their detection levels, while most markers for absorptive cells, such as brush border enzymes were increased to jejunal tissue levels (Fig. 1D, Fig. S1). The expression of markers for goblet, enteroendocrine and Paneth cells were either below or close to the limit of detection, underscoring the dominance of the absorptive cell linage. BO enteroids at day 6 of differentiation (BD6) in suspension represented an intermediate state compared to MG4 and AD6, with significant, but lower expression of several stem cell markers, and increased expression of absorptive cell markers (Fig. 1D, Fig. S1).

The time dependent gradual maturation towards a more differentiated phenotype is exemplified for all growth conditions and compared to freshly isolated enterocytes and jejunal villus tissue, using the proliferation marker MKI-67 and the brush border enzyme Aminopeptidase N as differentiation marker (Fig. 1E). Similar trends were observed for other markers, such as CD44 (proliferation marker) and SI (absorptive cell marker, Fig. S1). The substantial clustering distance observed in the PCA between enteroids in the growth phase (MG4) and mature differentiated enteroids (AD6) was further corroborated by differential expression analysis (Fig. 1F). Of the 3478 quantified proteins, in total 653 were differentially expressed between AD6 and MG4. Enrichment analysis revealed that biological processes enriched in MG4 were associated with proliferation, including RNA and DNA replication, regulation of such processes, and cell division (Fig. 1G). In contrast, pathways enriched in AD6 were related to xenobiotic metabolism and epithelial cell differentiation.

Overall, human jejunal 3D enteroids representing varying degrees of differentiation towards the mature villus phenotype were identified depending on the applied growth condition. AD6 enteroids exhibited the most differentiated state, closely resembling freshly isolated jejunal enterocytes and jejunal tissue. The near-complete absence of stem cell and proliferation markers, as well as markers for other cell types, suggested that AD6 enteroids consisted predominantly of mature absorptive enterocytes. Although BD6 enteroids retained some proliferative capacity, their expression of enterocyte differentiation markers approached that of AD6. We therefore focused our subsequent studies on AD6 and BD6 enteroids in suspension, hereafter referred to simply as AO and BO, respectively, in the remaining text.

### Apical-out and basal-out enteroids provide an intact epithelial barrier

The human intestinal epithelium serves as a physical barrier between the gut lumen and underlying tissue, maintained by tight junctions. The 3D AO and BO enteroids consisted of a continuous monolayer of enterocytes (Fig. 2). AO enteroids, which had ceased to proliferate (Fig. 1E), exhibited a predominantly spherical morphology with a median diameter of 138.4 ± 46.7 μm and a cross-sectional shape factor of 0.86 ± 0.08 (i.e., circularity factor) where 1.0 represents a perfect circle (Fig. 2A, 2C, 2D; Table S3). The resolution in the bright-field microscope sometimes allowed determination of the height (approximately 30 µm) and width (approximately 11 µm) of the AO enterocytes (Table S3), which is consistent with measurements reported for human jejunal enterocytes.^50^ This corresponds to approximately 174 enterocytes per AO enteroid. Scanning electron microscopy (SEM) revealed polygonal cell surfaces densely covered with microvilli, characteristic of a mature villus enterocyte phenotype. Occasionally, AO enteroids displayed ongoing cell extrusion (Fig. S2A). In intestinal tissue, most extruded cells are healthy, although a subset is caspase-3-positive.^51^ Apoptosis in differentiated enteroids was confirmed, as evidenced by the early (CASP8; higher expression than in tissue) and late (CASP3 and 7; comparable or lower (MG4) expression than in tissue, respectively) apoptosis markers (Fig. S2B). The expression of apoptosis markers in the differentiated enteroids align with the observation that the anti-apoptotic protein BCL2L1 decreased to tissue levels during differentiation (Fig. S2B).

**Figure 2.**
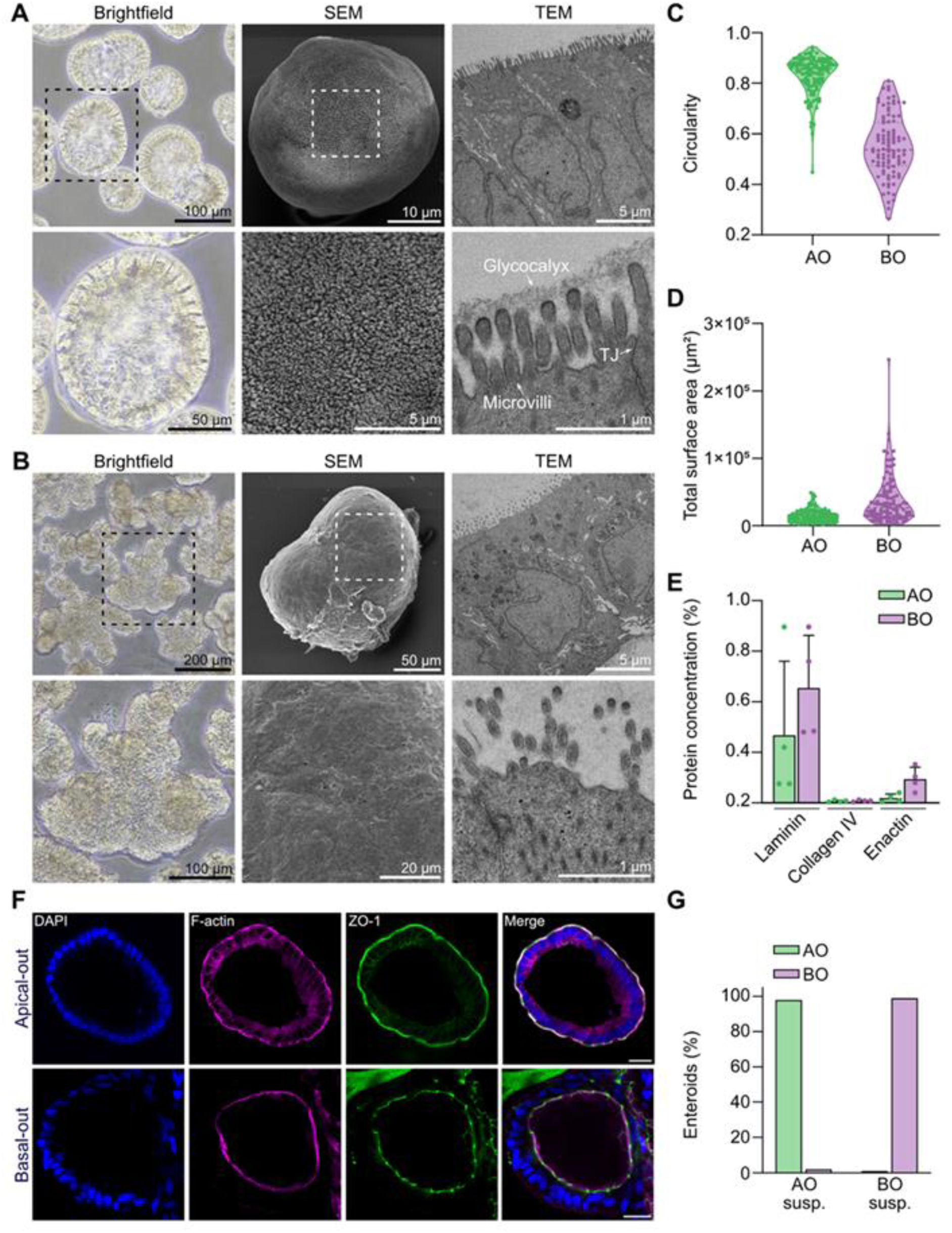
Characterization of morphology, size and shape distribution of differentiated 3D enteroids in suspension. **(A)** Representative micrographs of AD6 enteroids: bright field microscopy shows live enteroids in suspension (left), scanning electron microscopy (SEM) reveals a dense microvillar surface (middle), and transmission electron microscopy (TEM) shows the organized microvilli-rich brush border, tight junctions (TJ) and glycocalyx (right). High-magnification SEM-images (bottom row) correspond to the areas outlined by dotted squares in the upper row. **(B)** Representative micrographs of BD6 enteroids: live bright field microscopy illustrates typical basal-out morphology (left), SEM shows the smooth basal surface (middle), and TEM reveals the microvilli-rich brush border on the lumen-facing apical surface, confirming basal-out polarity (right). High-magnification views (bottom row) correspond to the areas outlined in the upper row. **(C)** Circularity analysis of AO (n = 118) and BO enteroids (n = 94) in suspension from bright field micrographs. (D). The total cross sectional surface area at 6 days of suspension culture for AO (n = 122) and BO (n = 94) enteroids. **(E)** Concentrations of Matrigel proteins present in AO and BO enteroids in suspension cultures after washing. Data from four donors are shown. Data are presented as means + SD. **(F)** Representative confocal images following staining of zonula occludens-1 (ZO-1, green), phalloidin (F-actin; magenta) and nuclei (DAPI; blue), highlighting the continuous monolayer barrier in AO and BO enteroids. Scale bars, 20 µm. **(G)** Quantification of apical-out and basal-out polarity in enteroid suspensions by confocal microscopy (n = 100 per population). Data is shown as mean.

Transmission electron microscopy (TEM) revealed that AO enteroids exhibit a columnar morphology closely resembling villus enterocytes *in vivo*. These cells displayed basally located nuclei and well-developed tight and adherens junctions (Fig. 2A). Proteins contributing to the normal appearance of microvilli^52^ were detected at concentrations comparable to *in vivo* levels and included: F-actin bundling/microvilli elongating proteins (*e.g.*, villin, espin, EPS8), membrane crosslinking proteins (*e.g.*, ezrin, PDCK1, myo1) and intramicrovillar anchoring proteins (*e.g.*, ANKS4B, CHHR2, CDHR5) (Table S4). The microvilli-anchored glycocalyx barrier was clearly visualized in the AO-oriented enteroids by TEM (Fig. 2A). Proteomic analysis further confirmed the presence of surface-bound mucins, including those integral to the glycocalyx structure (*e.g.*, MUC1, MUC13, MUC17). Additionally, several fucosyltransferases (*e.g.*, FUT2, FUT3, FUT8) were identified, that mediate the fucosylation of surface mucins and other glycoproteins, thereby fortifying the cell surface by creating complex branched carbohydrate structures (Table S4).

The BO enteroids, which retained some proliferative capacity (Fig. 1E), expanded with time in culture and formed crypt-like protuberances as previously described.^53,54^ This resulted in larger and more heterogenous enteroids with a median diameter of 260.5 ± 135.3 μm, a shape factor of 0.54 ± 0.12 (Fig. 2B-D and Table S3) and a clearly distinguishable lumen. Unlike the morphologically homogenous AO enteroids, the BO population included a subpopulation of smaller enteroids with a stem-like phenotype. These smaller structures displayed a more spherical shape and a thinner, less columnar cell layer, similar to what has been observed in fetal crypt-derived and induced pluripotent stem cells-derived BO enteroid suspension cultures (Fig. S3).^55^ SEM revealed that BO enteroids had a smooth basolateral surface without distinct cell borders, likely due to residual basement membrane components from Matrigel (Fig. 2B). Notably, only small amounts of Matrigel remained after washing prior to imaging and subsequent experiments (Fig. 2E and Fig. S4). TEM showed that BO-oriented enterocytes had a similar appearance to those in AO enteroids but were slightly less columnar. The lumen contained deposits of extracellular vesicles and proteins, and the glycocalyx appeared more diffuse. Despite these differences, proteins involved in microvillus structure and the glycocalyx barrier were present in amounts comparable to those found in AO enteroids (Table S4). Fluorescence staining revealed that over 98% of AO and BO enteroids exhibited correctly oriented, continuous actin belts and ZO-1-stained tight junctions, facing the surrounding medium in AO enteroids and the inner lumen in BO enteroids, respectively, which agrees with previous observations^15^ (Fig. 2F-G).

At the proteome level, key components of adherens junctions (*e.g.*, CDH1, CTNND1, vinculin) and tight junctions (*e.g.*, claudins, occluding, tricellulin) were quantified, with levels often comparable to those found in jejunal tissue (Fig. 3A, Table S5). Claudins (CLDs), which use their extracellular hydrophobic domains to mediate adhesion between adjacent cells, are categorized by their roles in gate (*e.g.*, CLD2, CLD15) and fence functions (*e.g.*, CLD3, CLD5, CLD7).^56^ Both classes of claudins were present in AO and BO enteroids, indicating the formation of fully functional junctional complexes (Fig. 3A). Interestingly, CLD2, which selectively permits the passage of small cations such as sodium ions, exhibited higher expression in BO enteroids. This elevated expression may be linked to its role in maintaining sodium ion homeostasis within the closed luminal space characteristic of this configuration (Table S5).^57^ The key proteins of desmosomes, which provide a secondary support system to the tight and adherens junctions in reinforcing epithelial integrity, were generally expressed at levels within 2-3-fold of those found in jejunal tissue, as exemplified by the extracellular adhesion proteins (*e.g.*, Dsg2 and Dsc2) and intracellular anchoring proteins (*e.g.*, desmoplakin and plakophilin-2, Table S5). In summary, both AO and BO enteroids formed continuous monolayer barriers, with well-developed microvilli, glycocalyx and junctional complexes, recapitulating key structural and protein expression features of the human intestinal epithelium.

**Figure 3.**
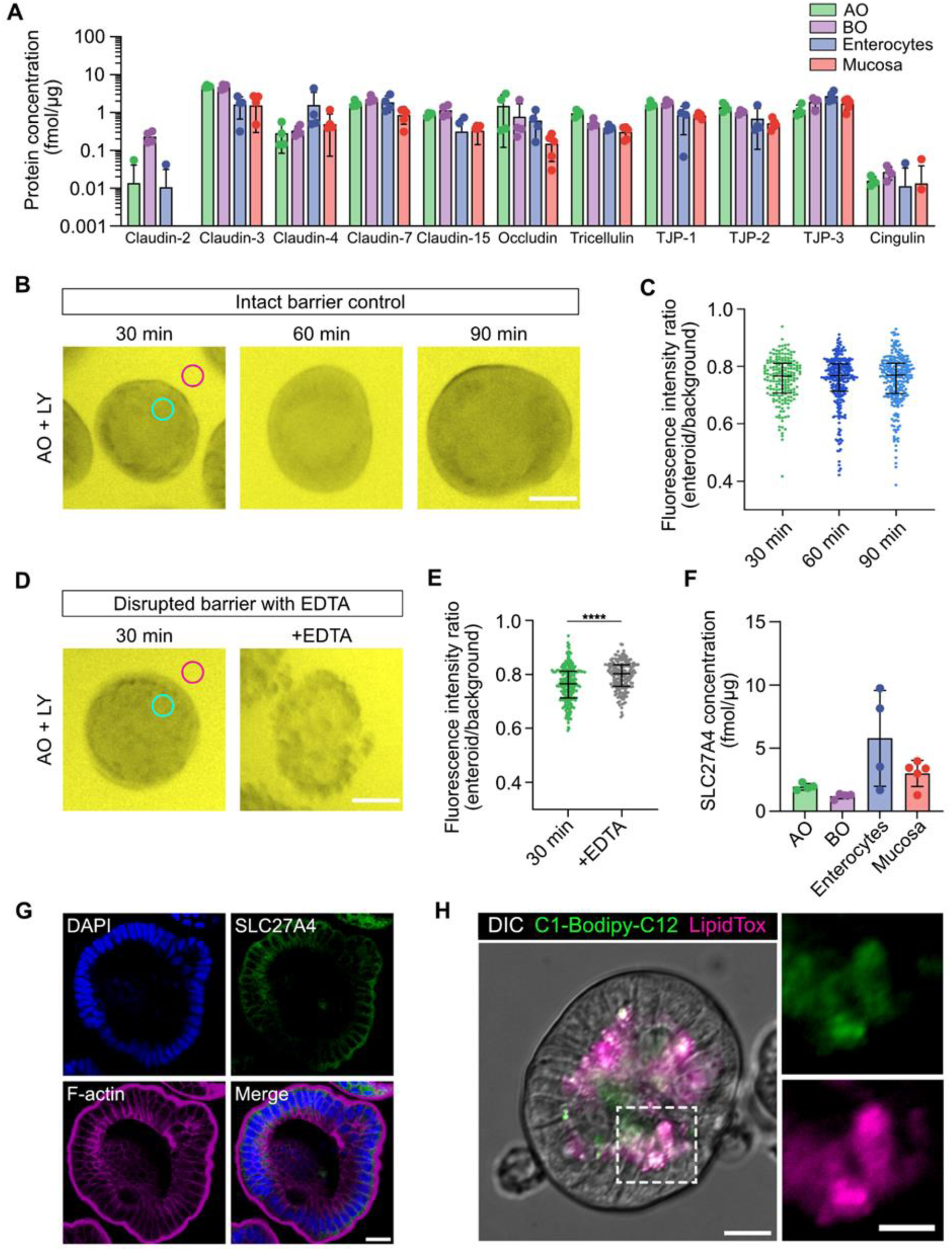
Epithelial integrity and fatty acid transport in 3D enteroids in suspension. **(A)** Expression of most important tight junction proteins in AO and BO as compared to primary enterocyte and jejunal mucosa *in vivo* levels. Data from four donors are shown. Data are presented as mean ± SD. **(B)** Representative time-lapse series showing suspended AO enteroids that were incubated in medium containing Lucifer yellow (LY, 100 µM) at 37°C to asses intact epithelial barrier over a time period of 120 min. Enteroids were imaged live by confocal microscopy and LY fluorescence intensity was determined within a region of interest within the enteroid (green ROI) and background (magenta ROI). Scale bars, 50 µm. **(C)** Quantification of LY fluorescence intensity within enteroids (green ROI) was normalized to background (magenta ROI). For the experiment, an average of 208 ± 66 observations per time point were made; Kruskal-Wallis ANOVA test showed no statistically significant difference between timepoints. Data is shown as median ± IQR. **(D)** Incubation with Ca^2+^ chelator EDTA disrupts paracellular barrier integrity in apical-out differentiated enteroids. The LY permeability assay was used to evaluate apical-out enteroids treated with EDTA for 30 min. Enteroids were imaged as described in (B) using live-cell confocal microscopy. Scale bars, 50 µm. **(E)** Quantification of LY fluorescence intensity within untreated enteroids (incubation with LY for 30 min) and enteroids treated with EDTA (additional incubation for 30 min with EDTA, green ROI) was normalized to background (magenta ROI). For the experiment, an average of 209 observations were made; ****p < 0.0001 (Mann-Whitney u test). Data is shown as median ± IQR. **(F)** Expression levels of SLC27A4 protein as determined by global proteomics in in AO and BO enteroids compared to primary enterocyte and jejunal mucosa *in vivo* levels. Data from four donors are shown. Data are presented as mean ± SD. **(G)** Representative confocal images following staining with DAPI (nuclei; blue), SLC27A4 (solute carrier family 27 member 4, green) and phalloidin (F-actin; magenta), highlighting expression pattern of the fatty acid transport protein in AD6 enteroids. Scale bars, 20 µm. **(H)** AD6 enteroids were incubated with fluorescent fatty acid analog C1-BODIPY-C12 for 10 min and LipidTOX-DR was used to visualize lipid droplets. Enteroids were imaged live using confocal microscopy. Scale bars, 20 μm; inset scale bars, 10 µm. Global protein profiling of human adult stem cell-derived enteroids in various culture formats.

Although proteome and imaging data indicated an intact and continuous epithelial barrier, enteroid populations exhibited heterogeneity in size, and for BO enteroids, also in shape and maturation (Fig. 2B). To assess whether this heterogeneity affects barrier integrity, we developed a semi-automated live-cell enteroid imaging method. This new technique enables the simultaneous investigation of the permeability of hundreds of enteroids using Lucifer Yellow (LY), a well-established low molecular weight hydrophilic marker for assessing epithelial barrier function.^58^ The LY assay measures barrier integrity by comparing mean gray value intensity differences between enteroids and the background. On average, the AO enteroid population excluded LY, demonstrating a retained barrier function over a time period of 90 min, making them suitable for functional experiments within this time interval (Fig. 3B-C). However, variability in intensity ratios around the mean suggested a degree of heterogeneity in barrier integrity across the population. Exposure to 2 mM EDTA which chelates calcium and thereby compromises the tight junction barrier, increased the luminal concentration of LY (Fig. 3D-E).

In contrast to AO enteroids, BO enteroids exhibited high luminal fluorescence (Fig. S5). This is attributed to Lucifer Yellow (LY) being a substrate of BCRP, an apically located ABC transporter (also called ABCG2) that effluxes metabolites and xenobiotics into the intestinal lumen *in vivo*.^59^ Consequently, LY is actively transported by BCRP into the closed BO lumen. To evaluate the barrier integrity of BO enteroids, we preloaded them with LY, removed excess dye through washing, and monitored changes in intensity over time. Similar to AO enteroids, the mean intensity ratio of BO enteroids remained constant for over time, indicating that the paracellular barrier remained intact across the majority of the population. Notably, a significant decline in intensity ratio was observed following 30 minutes of exposure to 2 mM EDTA (Fig. S5), further supporting that BO enteroids maintained an intact paracellular barrier under normal conditions.

### Apical-out and basal-out enteroids express an in vivo-like panel of jejunal membrane transporters and metabolizing enzymes

As any cells, the intestinal epithelium needs to take up nutrients to proliferate, differentiate and exert their functions. Further, the enterocytes at the villus tips are responsible for the selective absorption of critical nutrients from food, such as nutrients, vitamins and trace elements. These functions are handled by membrane-spanning transport proteins of which the solute carrier transporters (SLCs) is the largest with more than 455 members arranged into 66 families.^60^ We found 117-132 SLC-transporters expressed across enteroids, primary enterocytes and jejunal tissue samples (Table S6). For instance, transporters for important amino acids were found (*e.g.*, SLC1A5/ASCT2, SLC1A1/EEAT1, SLC7A5/LAT1, SLC36A1), as were transporters for glucose and other monosaccharides (*e.g.*, SLC5A1/SGLT1, SLC2A1/GLUT1 and SLC2A5/GLUT2). The expression levels of these transporters were generally in good agreement across sample types, and variations within 2 - 5-fold were often observed, although there were exceptions. The largest differences were observed for some nutrient and metabolite transporters such as the glucose uptake transporter GLUT1 and MCT4, which transports excess lactate and other monocarboxylates from the cell interior to the extracellular space, often as a result of increased glucose consumption in cell culture (Table S6). Both of these transporters were expressed at about 10-fold higher levels in the enteroids as compared to primary enterocytes and jejunal tissue. Nutrient transporters that had a lower expression in the enteroids included LAT2 (SLC7A8) and BAT1 (SLC6A19) and several orphan transporters (Table S6).

Here, we confirmed functional fatty acid transport in AO enteroids using our live-cell enteroid imaging set up (Fig. 3G-H).^61^ The fluorescent C12 analogue was rapidly taken up by AO enteroids and colocalized in intracellular lipid droplets stained with LipidTOX™ Deep Red, indicating functional apical transport in the AO enteroids, while only passive uptake and no co-localization could be observed in the BO enteroids.Both AO and BO enteroids expressed four members of the FATP proteins family (SLC27A1-4), while the expression of fatty acid translocase was low or below detection level (Fig. S6). The highest protein expression was observed for the medium and long chain acid (C12) transporter FATP4, both in the enteroids and tissues, making FATP4 the prime candidate for the C1-BODIPY-C12 transport (Fig. 3F).

We next focused on clinically relevant transporters that influence the pharmacokinetics and hence the effects of different drugs (Table S7).^62^ In general, the expression of these transporters and enzymes was lower in the MD6 than in the BO and AO enteroids (Fig. 4A and Fig. S7). Another trend was that transporters located in the membrane facing the nutritious cell culture medium tended to have a higher expression than those facing the closed inner luminal space. This is exemplified by the apically located di- and tripeptide transporter, PEPT1 (SLC15A) and for BO enteroids the two ABC transporters MRP2 (ABCC2) and Pgp (ABCB1) (Fig. 4A and Fig. S7). The corresponding examples for basolaterally located transporters in BO enteroids are MRP1 (ABCC1), MRP3 (ABCC3) and OST alpha/beta (SLC51a/b) (Fig. 4B-C). In general, the ABC-transporters had a similar or somewhat higher expression in the enteroids than in the tissues (Fig. 4B). The exception was the peroxisomal long chain fatty acid transporter ABCD1, which had a 10 - 20-fold lower enteroid expression (Fig. 4B and Table S8).

**Figure 4.**
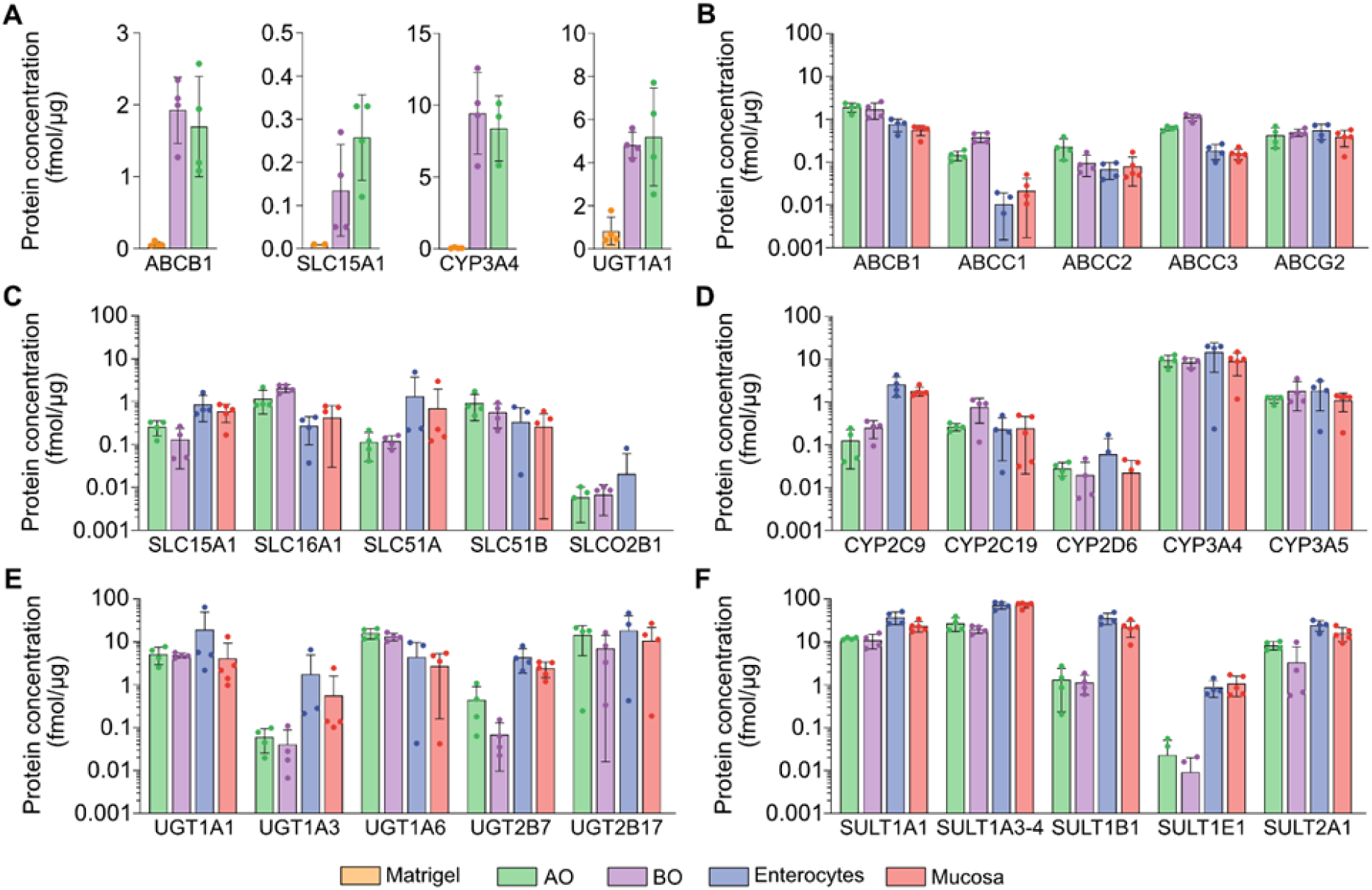
Protein expression levels of drug transporting and drug metabolizing enzymes in 3D enteroids in suspension. **(A)** Expression of transporters and enzymes in Matrigel-embedded enteroids (MD6) vs. apical-out (AO) and basal-out (BO) enteroids. **(B-F)** Expression of key transporters such as (B) ATP-binding cassette (ABC) transporters and (C) solute carrier (SLC) transporters, as well as key enzymes for (D) cytochrome P450 (CYP) enzymes, (E) UDP-glucuronosyltransferases (UGT), (F) sulfotransferases (SULT), and compared to primary enterocyte and jejunal mucosa *in vivo* levels. Data from four donors are shown. Data are presented as mean ± SD.

### The clinically important transporters Pgp and BCRP are functional in both apical-out and basal-out enteroids

When new drug candidates are filed for evaluation and approval by regulatory agencies, studies of drug transport and drug interactions with nine clinically relevant transporters (BCRP, MATE1, MATE2K, OAT1, OAT3, OATP1B1, OATP1B3, OCT2, Pgp) are recommended.^62^ Among these, Pgp and BCRP are particularly relevant in the small intestine. Therefore, we investigated their function in the enteroids. The expression of the intestinal efflux transporter Pgp was approximately two-fold higher compared to freshly isolated enterocytes and jejunal tissue (Fig. 4B and Fig. S8A). Immunofluorescence staining confirmed the correct localization of Pgp at the apical side, with staining observed along the cell borders (Fig. S8B). To evaluate Pgp functionality, we developed a live-cell microscopy assay using Rho123, a fluorescent substrate for the apically located Pgp efflux transporter (Fig. 5).^63^ In AO enteroids, the Pgp transporter efficiently pumped the fluorescent dye into the surrounding medium, leaving negligible intracellular amounts (Fig. 5A and C). In contrast, when the Pgp-specific inhibitor elacridar (ELC) was added, significant retention of Rho123 within the epithelium was observed. Conversely, for BO enteroids, Rho123 accumulated within the inner confined lumen, consistent with the apical (luminal) localization of Pgp (Fig. 5D-E). When ELC was added, efflux into the inner lumen was partially inhibited and Rho123 remained inside the enterocytes without being effluxed into the inner luminal space (Fig. 5F-G). Overall, the average fluorescence intensity in BO enteroids was fivefold higher than in the AO population (Fig. 5C, E and G). As observed also in the live-cell integrity assay with LY (Fig. 3B-E), the extent of Pgp transport varied within the organoid population (Fig. 5C, E and G).

**Figure 5.**
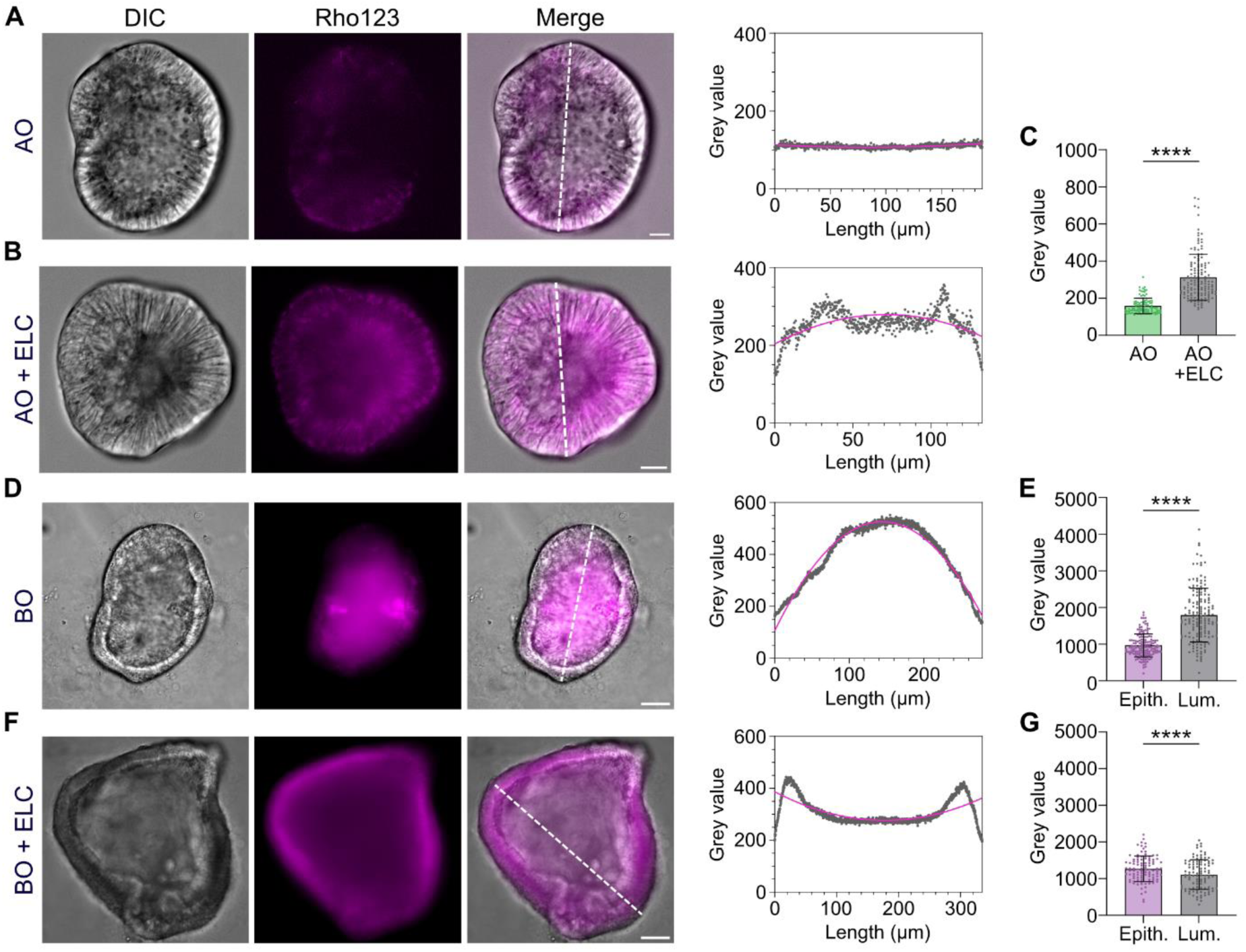
Real-time analysis of P-glycoprotein (Pgp)-mediated transport of Rhodamine123 (Rho123) in apical-out (AO) and basal-out (BO) enteroids. **(A)** Representative confocal images of AO enteroids following incubation with Rho123 (10 μM) at 37°C for 1 hour, followed by washing and live-cell imaging. The Rho123 fluorescence profile (right panel) corresponds to the white line in the merged image. Scale bars, 20 µm. **(B)** In control experiments, AO enteroids were pre-treated with the Pgp inhibitor elacridar (ELC, 2 μM) for 1 hour at 37 °C, then incubated with Rho123 (10 μM) in the presence of ELC (2 μM) for an additional hour. Representative live-cell confocal image shown, with the Rho123 fluorescence profile (right panel) corresponding to the white line in the merged image. Scale bars, 20 µm. **(C)** Quantification of Rho123 fluorescence intensity (gray values) between AO enteroids treated only with Rho123 (n = 284)) and control AO enteroids, treated with ELC prior to Rho123 addition (n = 227); ****p < 0.0001 (Mann-Whitney t-test). Data presented as mean ± SD. **(D)** Representative confocal images of BO enteroids incubated with Rho123 (10 μM) at 37°C for 1 hour, washed, and live imaged. The fluorescence profile (right panel) corresponds to the white line in the merged image. Scale bars, 20 µm. **(E)** Quantification of Rho123 fluorescence intensity (gray values) within BO enteroid epithelium and lumen (n = 154); ****p < 0.0001 (Wilcoxon paired t-test). Data shown as mean ± SD. **(F)** BO enteroids pre-treated with ELC (2 μM) for 1 hour, followed by Rho123 incubation (10 μM) with ELC for another hour at 37 °C. Representative confocal image and fluorescence profile (right panel) corresponding to the white line in merge. Scale bars, 20 µm. **(G)** Quantification of Rho123 fluorescence intensity in ELC treated BO enteroid epithelium and lumen (n = 102); ****p < 0.0001 (Wilcoxon paired t-test). Data shown as mean ± SD.

The expression of the other clinically important intestinal ABC-transporter BCRP was comparable to that of freshly isolated enterocytes and jejunal tissue (Fig. 6A). Immunostaining confirmed the correct localization of BCRP at the apical side, where it overlapped with the actin belt of the microvilli and terminal web (Fig. 6B). Although most enterocytes exhibited distinct BCRP staining, the intensity varied across the population (Fig. 6B). While the live-cell microscopy assay developed for Pgp provided a detailed fingerprint of functional heterogeneity within the organoid population, obtaining mean values is often sufficient for transporter assays. To evaluate BCRP functionality, we loaded AO enteroids with the fluorescent BCRP substrate pheophorbide A^64^ in the presence of the BCRP inhibitor KO143^65^. Efflux was measured under both conditions, and the ratio of the two efflux values was used as an indicator of efflux efficiency (Fig. 6C). The efflux ratio observed in the AO enteroids was higher than that previously reported in MDCK cells overexpressing BCRP.^29^ This finding indicates that BCRP effectively expels its substrate from the AO enteroid model.

**Figure 6.**
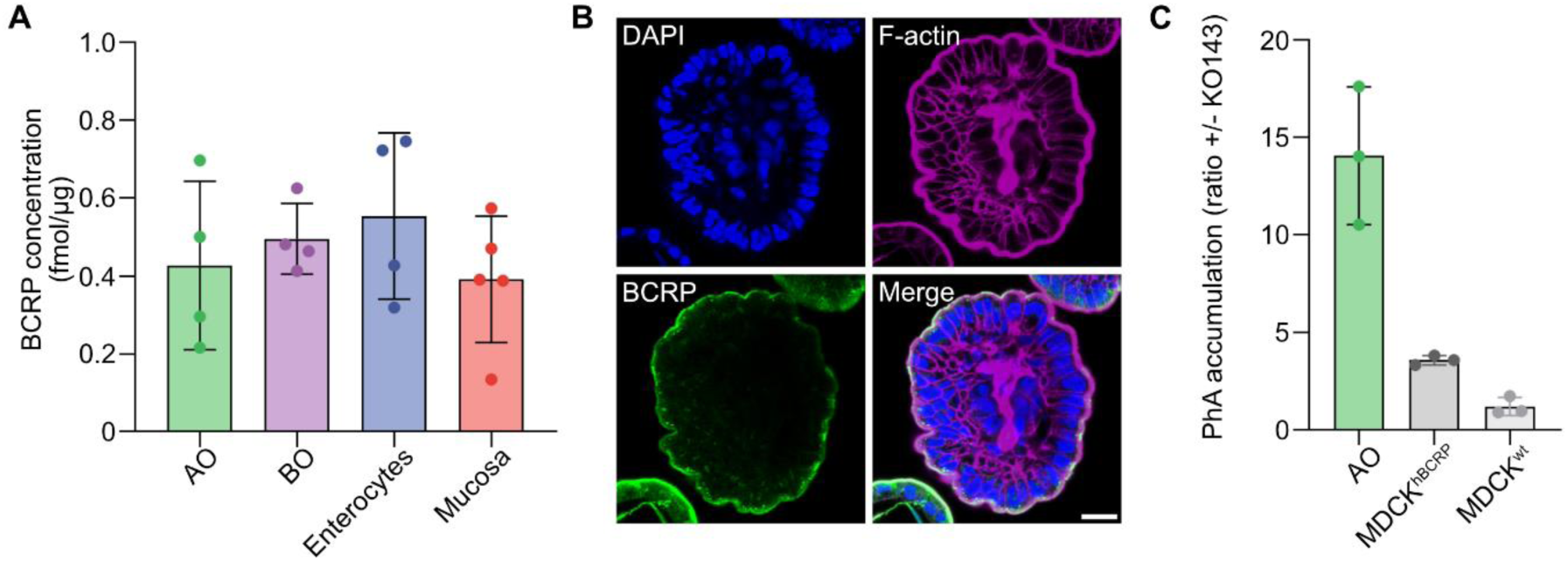
BCRP expression, localization and activity in apical-out enteroids. **(A)** Expression levels of BCRP protein as determined by global proteomics in AO and BO enteroids compared to primary enterocyte and jejunal mucosa *in vivo* levels. Data from four donors are shown. Data are presented as mean ± SD. **(B)** Representative confocal images following staining with DAPI (nuclei; blue), BCRP (green) and phalloidin (F-actin; magenta), highlighting expression pattern of the BCRP transporter in AD6 enteroids. Scale bars, 20 µm. **(C)** BCRP efflux in AD calculated by PhA accumulation; transporter substrate: PhA, transporter inhibitor: KO143. Data are presented as mean ± SD.

### Apical-out and basal-out enteroids express key metabolizing enzymes at in vivo levels

Widespread models of the intestinal epithelial barrier, such as Caco-2 monolayers, express an aberrant panel of metabolic enzymes due to their cancerous origin.^7^ So far, a small intestine barrier model with metabolic enzymes that accurately reflect those of mature villus enterocytes has not been fully characterized. We found that the differentiated AO and BO enteroids showed an almost complete overlap of quantifiable phase I and phase II drug metabolizing enzymes with primary enterocytes and jejunal tissue samples. For example, all 17 quantifiable cytochrome P450 (CYP) enzymes, which mediate oxidative reactions, were present in all samples (Table S9). Among these, CYP3A4, the clinically most important CYP enzyme responsible for metabolizing approximately 50% of all drugs^66^ was expressed at levels within two-fold of those in primary enterocytes and tissues (Fig. 4D and Table S9). Other clinically relevant polymorphic isoenzymes showed variable expression levels compared to *in vivo* conditions. For instance, CYP2C19 protein levels were ∼2-6 times higher, CYP2D6 ∼1-5 times lower, and CYP2C9 was 5 - 20 times lower in enteroids than in primary enterocyte or jejunal tissue samples (Fig. 4D and Table S9).

In addition to CYP enzymes, other phase I enzymes including carboxylesterases (CESs), which catalyze the hydrolysis of various substrates^67^, showed comparable expression levels between enteroids and enterocytes (Table S10). CES1, known for hydrolyzing substrates with larger acyl groups, and CES2, which targets smaller acyl groups such as CPT-11 and betamethasone valerate, were both detected in the enteroid samples. Alcohol dehydrogenases (ADHs) and aldehyde dehydrogenases (ALDHs), which play key roles in ethanol metabolism^68^, were expressed in AO and BO enteroids (Table S10). Within the ADH1 family, ADH1B and ADH1C were quantified, though their expression levels were approximately 4-fold lower than those in primary enterocytes and jejunal mucosa. Additionally, flavin-containing monooxygenases (FMOs) were assessed, revealing FMO5 as the most abundant isoform across all samples (Table S10).^69,70^ The high expression of FMO5 underscores its significant role in metabolic processes within both the enteroids and the small intestine.

We next evaluated phase II enzymes, which are essential for conjugation reactions that enhance metabolite and drug solubility and elimination. As for the the phase I enzymes, most phase II enzymes were quantifiable both in enteroids, primary enterocytes and jejunal tissue samples. For instance, UDP-glucuronosyltransferases (UGTs), which catalyze the addition of glucuronic acid to compounds like acetaminophen, morphine, and estrogens, showed comparable expression levels in enteroids and enterocytes.^71^ Among 12 UGTs analyzed, UGT1A1, UGT1A6, and UGT1A10 were expressed at higher levels in enteroids than *in vivo*, whereas others, such as UGT1A3, UGT2B7, and UGT2B17, showed lower expression (Fig. 4E and Table S11). Similarly, sulfotransferases (SULTs), which transfer sulfate groups to endogenous compounds and drugs, making them more water-soluble, showed slightly reduced expression levels (∼2-5-fold lower) compared to enterocytes and jejunal tissue (*e.g.*, SULT1A1, SULT1A3, and SULT2A1, Fig. 4F).^72^ However, some enzymes, such as SULT1C2, were expressed at higher levels in enteroids, while others, such as SULT1B1 and SULT1E1, were expressed at much lower levels (Fig. 4F and Table S12).

Another important enzyme family, the glutathione S-transferases (GSTs), plays a central role in detoxification by conjugating glutathione to xenobiotics and reactive oxygen species.^73^ Eleven GST enzymes were quantified, with GSTA1, GSTM1, and GSTT1 showing similar expression levels across enteroids, enterocytes, and jejunal tissue (Table S13). GSTP1, however, exhibited significantly higher expression in enteroids (∼2-3-fold higher) compared to *in vivo* levels. Finally, N-acetyltransferases^74^ (NAT1 and NAT2), which catalyze the acetylation of drugs and carcinogens, were expressed at levels comparable to those *in vivo* (Table S13). Together, these results further highlight the metabolic fidelity of AO and BO enteroids and underscore their utility as models for studying intestinal metabolism.

### Apical-out enteroids exhibit functional CYP3A4 metabolism and enable PBPK modeling

The enzyme expression studies demonstrated that optimally differentiated suspension enteroids exhibit an enzyme expression profile closely resembling that of freshly isolated jejunal enterocytes and jejunal tissues. Based on this, we investigated the function of CYP3A4, the most clinically relevant drug-metabolizing enzyme, using two orally administered drugs whose pharmacokinetics are strongly influenced by intestinal CYP3A4 metabolism: terfenadine and midazolam (Fig. 7 and Fig. S9, respectively). The majority of the enterocytes in the AO enteroids had a strong intracellular CYP3A4 staining, mainly localized along the cell borders (Fig. 7A). The expression level in both AO and BO enteroids was approximately half that of freshly isolated enterocytes but comparable to jejunal mucosa (Fig. 7B). Terfenadine, which is rapidly metabolized by CYP3A4 to fexofenadine and, in parallel, azacyclonol^75^, was efficiently cleared from the medium when added to the enteroids, while fexofenadine simultaneously appeared in the medium (Fig. 7C). The intrinsic clearance of AO enteroids was 2.6 x µL/min/1×10^6^ cells, approximately 6-fold lower than that of freshly isolated jejunal enterocytes (Fig. 7D). However, after adjusting for differences in CYP3A4 protein expression (0.28 µL/min/fmol/µg CYP3A4 for AO vs. 0.75 µL/min/fmol/µg for enterocytes), the difference in intrinsic clearance was reduced to approximately 3-fold (Fig. 7E). PBPK modeling of terfenadine, where we incorporated AO enteroid metabolism rates corrected for CYP3A4 expression, was performed to evaluate the effectiveness of the model in assessing the contribution of intestinal metabolism (F_g_). The simulation showed the *in vivo* results following a 60 mg dose of terfenadine to 140 virtual male subjects aged 20 – 50 (Fig. 7E). The simulation based on the AO enteroid data gave a slight overprediction of the plasma concentration as compared to the clinical data, but the results remained well within the confidence limits of the virtual population (Fig. 7F). Similar findings were obtained for the more slowly metabolized drug midazolam (Fig. S9). These results indicate that AO enteroids are metabolically active and can be used to predict the impact of intestinal metabolism on systemic drug exposure.

**Figure 7.**
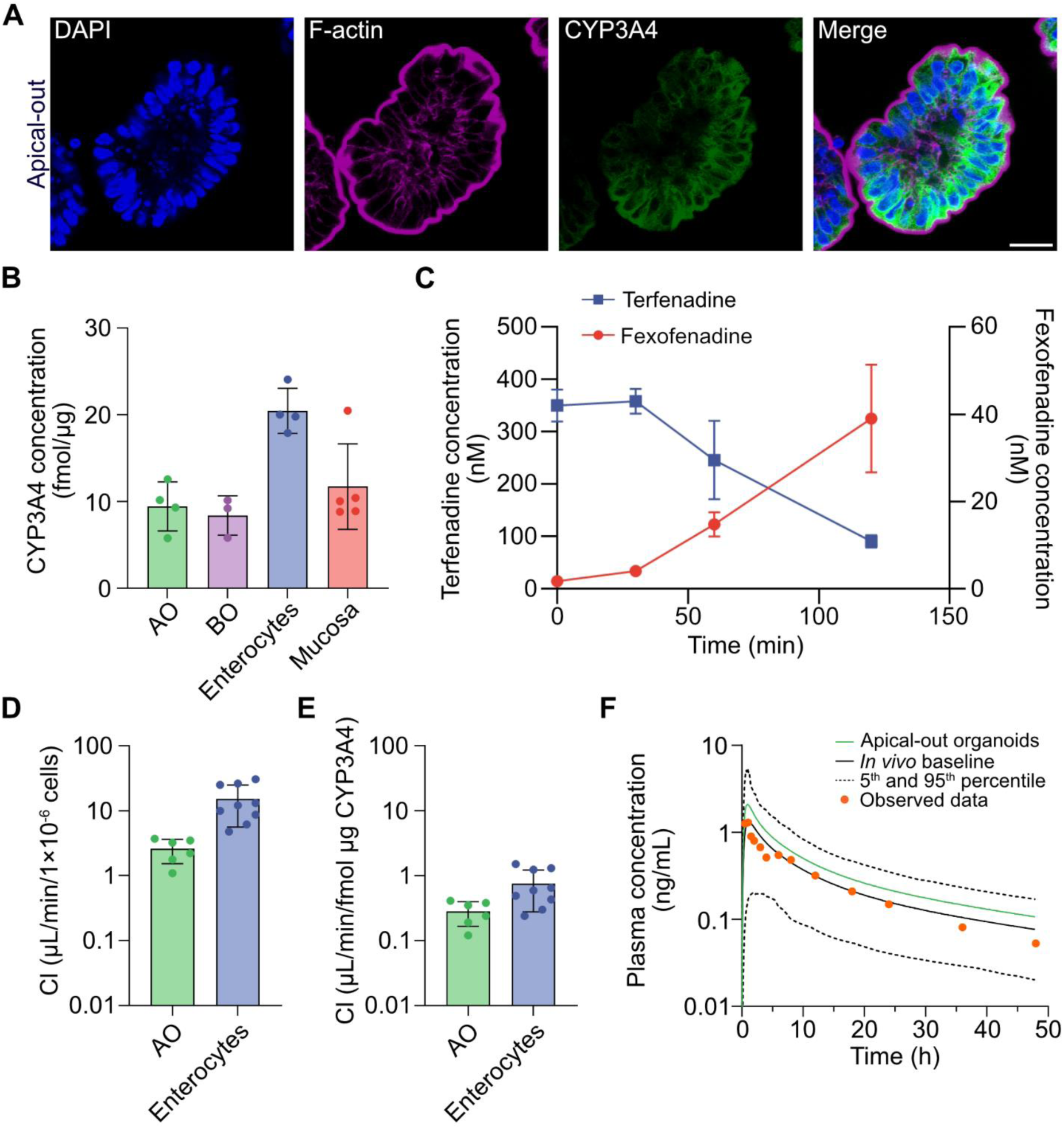
CYP3A4 metabolic activity in apical-out enteroids and PKPD simulation. **(A)** Representative confocal images following staining with DAPI (nuclei; blue), CYP3A4 (green) and phalloidin (F-actin; magenta), highlighting expression pattern of the CYP3A4 metabolic enzyme in AO enteroids. Scale bars, 20 µm. **(B)** Expression levels of CYP3A4 protein as determined by global proteomics in AO and BO enteroids compared to primary enterocyte and jejunal mucosa *in vivo* levels. Data from four donors are shown. Data are presented as mean ± SD. **(C)** CYP3A4 metabolic activity in AO over a time period of 120 min. The concentration of the enzyme substrate terfenadine was monitored over time as well as the formation of its metabolite fexofenadine. Data is shown as mean ± SD. **(D)** Intrinsic clearance of terfenadine in apical-out enteroids and primary enterocytes. Data is shown as mean ± SD. **(E)** Corrected intrinsic clearance of terfenadine considering the differences in CYP3A4 protein expression in apical-out enteroids and primary enterocytes. Data is shown as mean ± SD. **(F)** Simulated pharmacokinetics of a 60 mg dose of terfenadine in 140 virtual males (20–50 years, black line) with its 5th and 95th percentiles shown as dashed lines. The green line represents the PBPK model mean prediction for apical-out enteroids (log concentration). The orange dots depict mean observed data^37^.

## Discussion

In this study, we established conditions for optimally differentiated intestinal enterocytes from adult jejunal stem cells in 3D enteroids in suspension, both in their native basal out orientation and after morphogenic eversion to the more applicable apical-out orientation.^15^ We used quantitative global proteomics and different microscopic techniques to follow the maturation of the enterocytes into differentiated enteroids, that had a phenotype resembling absorptive cells at the jejunal villus tips. We then confirmed an intact barrier function by observing that the protein expression and subcellular location of barrier forming structures such as microvilli bound glycocalyx and tight junctions, were in qualitative and quantitative agreement with those of freshly isolated jejunal enterocytes and tissue. Further, we investigated the barrier function using a newly developed live-cell microscopy technique and the small molecular probe LY, which provided the opportunity to observe barrier imperfections and enteroid heterogeneity. We also discovered that the expression of membrane transporters and metabolizing enzymes were in good agreement with those observed *in vivo* and investigated the function of the clinically most important drug transporting and drug metabolizing proteins in the human intestine. Finally, we incorporated the enteroid-mediated intestinal metabolism of two drugs into a physiologically based pharmacokinetic population model and predicted the clinical pharmacokinetics of both drugs with good precision. We conclude that the differentiated enteroids presented in this study provides a versatile and, in our hands, reproducible “mini-gut” with broad applicability not only in mechanistic studies of the normal and diseased intestine, but also in studies of transport and metabolism of food constituents, drugs and other xenobiotics.

The exposure of the intestinal epithelial barrier to ingested material in the intestinal lumen takes place at the villus tips in the upper small intestine, the jejunum. The by far predominant cell type at the villus tips is the absorptive enterocyte, which accounts for about 95% of the cells in the upper small intestine, while the mucus-producing goblet cells make up most of the remaining 5% together with a small subpopulation of enteroendocrine cells.^76^ The focus of this study was to develop organoid cultures of differentiated absorptive cells equipped with both physical and metabolic barriers, providing a selective barrier that facilitates the absorption of essential nutrients via solute transporters while preventing exposure to toxicants, *e.g.*, via ABC transporters. Thus, in contrast to many other enteroid protocols, where efforts are made to assure that all epithelial cell types along the crypt-villus axis are represented, our goal was to provide a model that is representative of the villus tip enterocytes. The dominance of differentiated absorptive cells in our enteroids with a correspondingly low frequency of goblet cells precludes the formation of a continuous extracellular mucus barrier based on secreted soluble mucins. Detailed protocols that enhance the number of secretory cells such as goblet cells to about 5% in human enteroids have recently been provided^77^, but require increased stemcellness, which reduces the frequency of the differentiated absorptive cells with dense microvilli and an *in vivo*-like glycocalyx barrier (Table S4) that characterize the villus tips.

We thus endeavoured to differentiate our enteroids towards the absorptive cell lineage. We achieved this by extending the culture time in differentiation medium to such an extent that the expression of stem cell and proliferation markers such as SOX9 and MKI67, respectively, was minimized (BO enteroids) or below the limit of quantification (AO enteroids), and the expression of mature absorptive cell markers such as the brush border enzymes, *e.g.*, ANPEP and SI approached *in vivo* levels (Fig. 1). This was not without risk, as long-term culture of 3D enteroids leads to terminal differentiation and increased cell death. However, the levels of the late apoptotic markers were comparable to those found in our jejunal tissues, indicating that the extent of apoptosis was within the physiological range. Consistent with previous findings, the reduced but significant expression of stem cell and proliferation markers in the differentiated BO enteroids resulted in the formation of crypt-like structures that protruded into the surrounding medium and increased in size over time, whereas the non-proliferating differentiated AO enteroids remained largely circular and did not increase in size. Apart from the differentiation-promoting curvature, the limited size of the confined luminal space of AO enteroids constitutes a mechanical obstacle to the development of intraluminal crypts.

Despite these differences, the protein expression levels of differentiation markers, microvilli and tight junction barrier-related proteins were often remarkably similar between the differentiated AO and BO organoids. Moreover, these proteins were generally expressed at the same levels as in the freshly isolated villus enterocytes and human jejunum, indicating a high similarity to the corresponding barriers *in vivo*. This finding was supported by the immunohistochemical, TEM and SEM data which shows on continuous tight monolayers of correctly oriented and sized, fully differentiated enterocytes, with polygonal cell surfaces that have well developed microvilli with an attached thick glycocalyx barrier and all junctional complexes in place (Fig. 2).

Thus, the *in vivo* like expression of proteins of the tight and adherence junctions suggested a fully functional tight junction barrier and the apoptotic cells observed in the SEM and TEM had retained tight junction connections.^78^ Previous investigations on enteroid barrier integrity have often used fluorescently labelled dextran of a mean molecular weight of 4 kD as a probe for epithelial barrier integrity in enteroids.^79,80^ This is an insensitive probe that will fail to discover small imperfections in barrier integrity. In clinical studies of disease associated changes in intestinal permeability, much smaller hydrophilic probes are used, such as the lactulose-mannitol ratio (MW 342 and 182 respectively), where lactulose is impermeable, and mannitol is slowly absorbed from the healthy intestine.^81^ Similarly, orally administered drugs has an average size of between 200-500 D.^82^ To obtain a more nuanced analysis of the enteroid barrier, we therefore selected lucifer yellow (LY) as a fluorescent small molecular probe (MW 457 D) and measured the exclusion of this probe in hundreds of AO and BO enteroids. While live-cell microscopy showed that both AO and BO enteroids retained their tight junction barrier for at least 90 min, it also revealed that the barrier function was more heterogenous than expected from the proteomes and fixed microscope specimens. This is in agreement with two previous studies on AO enteroid permeability to LY. In the first study, where the integrity was compromised by exposure to lipopolysaccharide and oxygen depletion, a variable uptake of LY into the enteroid lumen was observed.^83^ In the second study, the variability was reduced by preselection of BO enteroids of comparable size and shape that were investigated one by one.^84^ In our study, we analyzed all AO enteroids and measured fluorescence using mean gray values, which may have contributed to variability in the population.. Nonetheless, in our hands, the individual AO enteroids excluded LY throughout the experimental period. In the case of the reverted experiments using BO enteroids, exclusion of stem-like BO enteroids without a luminal focal plane was required to obtain a population that was leak-proof to LY.

The high and *in vivo* like protein expression of nutrient and drug transporting proteins in the AO and BO enteroids prompt us to investigate their function. Previous investigation have used the fluorescent fatty acid analogue BODIPY-C12 to illustrate AO functional nutrient transport.^15^ Since the expression levels of the FATPs in the enteroids were in agreement with those *in vivo*, we set out to confirm these studies. However, our live-microscopy studies of fatty acid transport in the AO organoids turned out to be more complicated than expected, since the C12 analogue was taken up by not only FATP4 but also by passive distribution into the cell membranes. Thus, the cell lining of both AO and BO enteroids rapidly became fluorescent, and no polarity in cellular uptake could be seen. However, uptake and disposition of the C12 analogue into intracellular lipid droplets was polar and only occurred in AO enteroids, confirming FATP-mediated active transport of C12.

In drug discovery, it is mandatory to investigate the absorption mechanisms of drug candidates. For this purpose *in vitro* models of the intestinal epithelium, such as Caco-2 monolayers are routinely used, despite their well-known limitations.^7^ Of particular interest are, two clinically important drug transporters that limit drug and uptake after oral drug administration: the ABC-transporters P-glycoprotein (Pgp) and breast cancer resistance protein (BCRP).^62^ We first investigated Pgp functionality in the enteroid population using live-cell microscopy and the fluorescent substrate Rhodamine 123 (Rho123).^63^ Overall, Pgp was fully functional and effluxed Rho123 in both AO and BO enteroids. As in the integrity studies with LY, the efflux activity varied in the enteroid population. Nevertheless, the resolution was high and the effect of the specific Pgp inhibitor elacridar on the Pgp transport was highly significant in the population. Further, the transport and effect of the inhibitor could be clearly visualized in the individual enteroids. Once automated, the Pgp assay presented here will provide a new tool for FDA required investigations of drug-drug interaction studies with Pgp. In its present semi-automated format the assay is time consuming so for the studies of BCRP functionality, we instead performed the investigation of AO enteroids in the plate reader, using pheophorbide A as a specific BCRP substrate.^64^ The BCRP function was confirmed and the efflux was in fact even higher than that observed in MDCK cells overexpressing BCRP.^29^ We conclude that both of the two clinically important intestinal ABC-transporters are fully functional in the jejunal enteroids.

The bioavailability and hence effect of an orally administered drug is dependent on how much of the drug is eliminated during the “first pass” through the intestinal epithelium (F_g_) and liver (F_h_). While F_h_ can be predicted from *in vitro* models based on human hepatocytes with preserved metabolic capacity, the corresponding *in vitro* models of the human intestine have not expressed the right panel of drug metabolizing enzymes. Recent studies in intestinal enteroids have provided promising but variable results in this regard, mainly in 2D monolayers but also in other formats. In one case, induction via the vitamin D receptor was used to enhance metabolism by the major drug metabolizing enzyme CYP3A4^85^, supporting our observation of weak baseline metabolism in the 2D format, while others obtained *in vivo*-like CYP3A4 metabolism without induction, e.g.^19^ Our extensive mapping of the metabolic enzymes in the 3D enteroids showed *in vivo* like levels of all investigated drug metabolizing classes, suggesting that they are metabolically competent and can be used for predictions of F_g_. We therefore investigated the metabolic clearance of two drugs that are mainly metabolized by CYP3A4, and after normalization of CYP3A4 protein expression, predicted the human bioavailability in a virtual patient population with good accuracy. These results provide a first indication of the usefulness of the human jejunal enteroids for predictions of presystemic metabolism in humans.

Transporters and metabolic enzymes often have the same substrates or work in series. For example, a drug or toxicant can be taken up by an SLC transporter, metabolized by multiple enzymes inside the enterocyte and then the metabolite can be effluxed by an ABC transporter. The functional transport and metabolic processes, such as those reported in this study therefore opens up for a multitude of studies of transporter-enzyme interplay that have hitherto not been possible. Another possibility is that slowly formed metabolites, that are not detectable in short term studies with primary enterocytes can be added to the differentiated organoid cultures and allowed to accumulate and be concentrated in the closed luminal space or extracellular medium over prolonged time periods. Finally, many of the metabolic enzymes have detoxifying functions, and their expression at *in vivo* like levels opens up for more predictive toxicity studies.

In conclusion, our enteroids of differentiated jejunal absorptive epithelial cells has a proteome that closely resembles the fully differentiated absorptive cells found at the villus tips in the human jejunum, provide a complete epithelial cell barrier and is fully functional with regard to the here investigated transporter and enzyme activities. The AO and BO jejunal enteroids hence provide emerging physiologically relevant models for studies of intestinal epithelial function in health and disease, as well as in studies of intestinal drug transport and metabolism interplay.

## Author contributions

Conceptualization: MES, MH, PA; Methodology: MC, TF, MLDM, RH, ACCL, JE, DSV, IG, PL, DH, MES, MH, PA; Investigation: MC, TF, MLDM, RH, ACCL, JE, DSV, IG, PL, DH; Formal analysis: MC, TF, MLDM, RH, ACCL, JE, DSV, IG, PL, DH; Resources: MSu, Msk, PMH, DLW, MES, PA; Visualization: MC, FT, MH; Supervision: MK, PL, MES, MH, PA; Project administration: MSu, Msk, PMH, DLW, MES, MH, PA; Funding acquisition: MES, MH, PA; Writing - Original Draft: MC, FT, DSV, MH, PA; Writing - Review & Editing: All authors.

## Acknowledgement

The Republic of Turkey, Ministry of National Education is acknowledged for providing financial support under the YLSY doctoral scholarship program (MC). This research was funded by the Swedish Research Council grant no. 2020-05186 and 2017-01951 (PA), Swedish Foundation for Strategic Research grant no. FFL18-0165 (MES), VINNOVA (2019-00048) for the Swedish Drug Delivery Center (SweDeliver) and by the European Union’s Horizon 2020 research and innovation program under the Marie Skłodowska-Curie grant agreement no. 956851 (RH). This work was supported by the project RISK-HUNT3R: RISK assessment of chemicals integrating HUman centric Next generation Testing strategies promoting the 3Rs. RISK-HUNT3R has received funding from the European Union’s Horizon 2020 research and innovation programme under grant agreement No 964537 and is part of the ASPIS cluster (PA and PL). This work reflects only the authors’ views, and the European Commission is not responsible for any use that may be made of the information it contains. Additionally, the research was supported by the European Union (GA 101057491, PA and MH). Views and opinions expressed are however those of the author(s) only and do not necessarily reflect those of the European Union or the European Health and Digital Executive Agency (HaDEA). Neither the European Union nor the granting authority can be held responsible for them. The authors acknowledge the facilities and technical assistance of the Umeå Centre for Electron Microscopy (UCEM) at the Chemical Biological Centre (KBC), Umeå University, a part of the National Microscopy Infrastructure NMI (VR-RFI 2019-00217).

## Supplementary Information

**Table S1.** Group level demographic information on the donor tissue samples used in this study. The age groups included young adult (< 31), middle-age adult (31-45) and old-aged adults (46-75).

| **Age group** | **No** | **% Female** |
| --- | --- | --- |
| Young adult | 2 | 100 |
| Middle -age adult | 9 | 90 |
| Old-age adult | 4 | 75 |
| Total | 15 |  |
| Average age | 40 |  |

**Table S2.** Input parameters for the terfenadine PBPK model.

| Parameter | Value | Source |
| --- | --- | --- |
| MW | 471.7 | Pubchem |
| Log P | 5.01 | ter Laak et al 1994<sup>1</sup> |
| pKa | 8.6 (base) |  |
| BP | 0.78 | Predicted in the simcyp simulator |
| $f_{u_{\text{plasma}}}$ | 0.03 | Chen et al 2007<sup>2</sup> |
| Absorption | ADAM model<br>(solution) | Jamei et al 2009<sup>3</sup> |
| $F_{u_{\text{gut}}}$ | 0.15 | estimated |
| Elimination – HLM (ml/min/mg<br>microsomal protein) | 13 342 | CYP3A4 |
| Elimination – HLM (ml/min/mg<br>microsomal protein) | 272 | Non-CYP3A4 metabolism |

**Figure S1.**
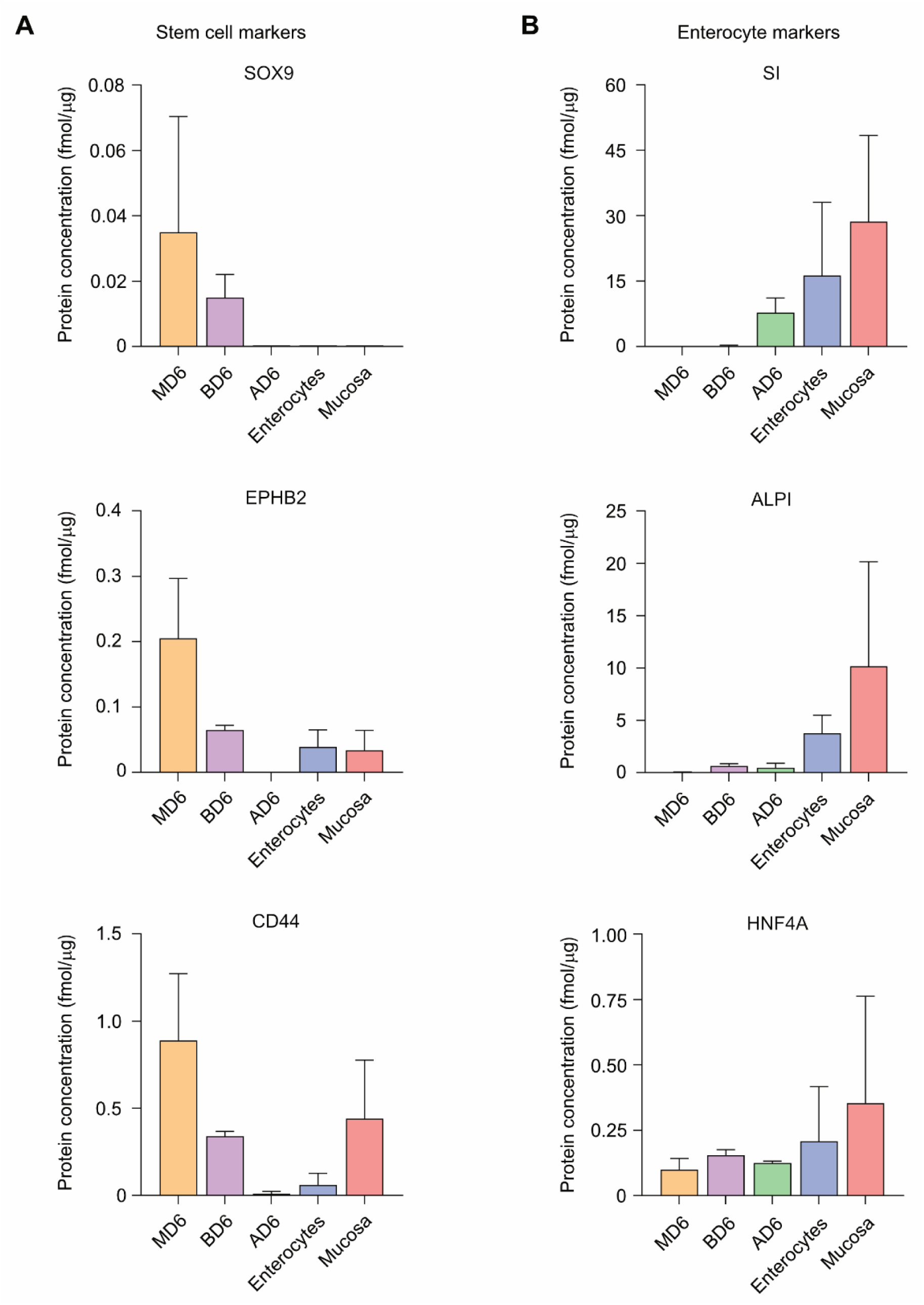
Global protein profiling of human enteroids in various culture formats. **(A)** Stem cell and **(B)** enterocyte marker protein expression in enteroids, enterocytes and jejunal mucosa. Quantification of SOX9, EPHB2, CD44 (stem cell markers, left) and SI, ALPI, HNF4A (enterocyte markers, right) across 3D enteroid formats (M: Matrigel, B: basal-out, A: apical-out) cultured in ODM for 6 days, compared to primary enterocyte and jejunal mucosa *in vivo* levels. Data are presented as mean ± SD.

**Table S3.**
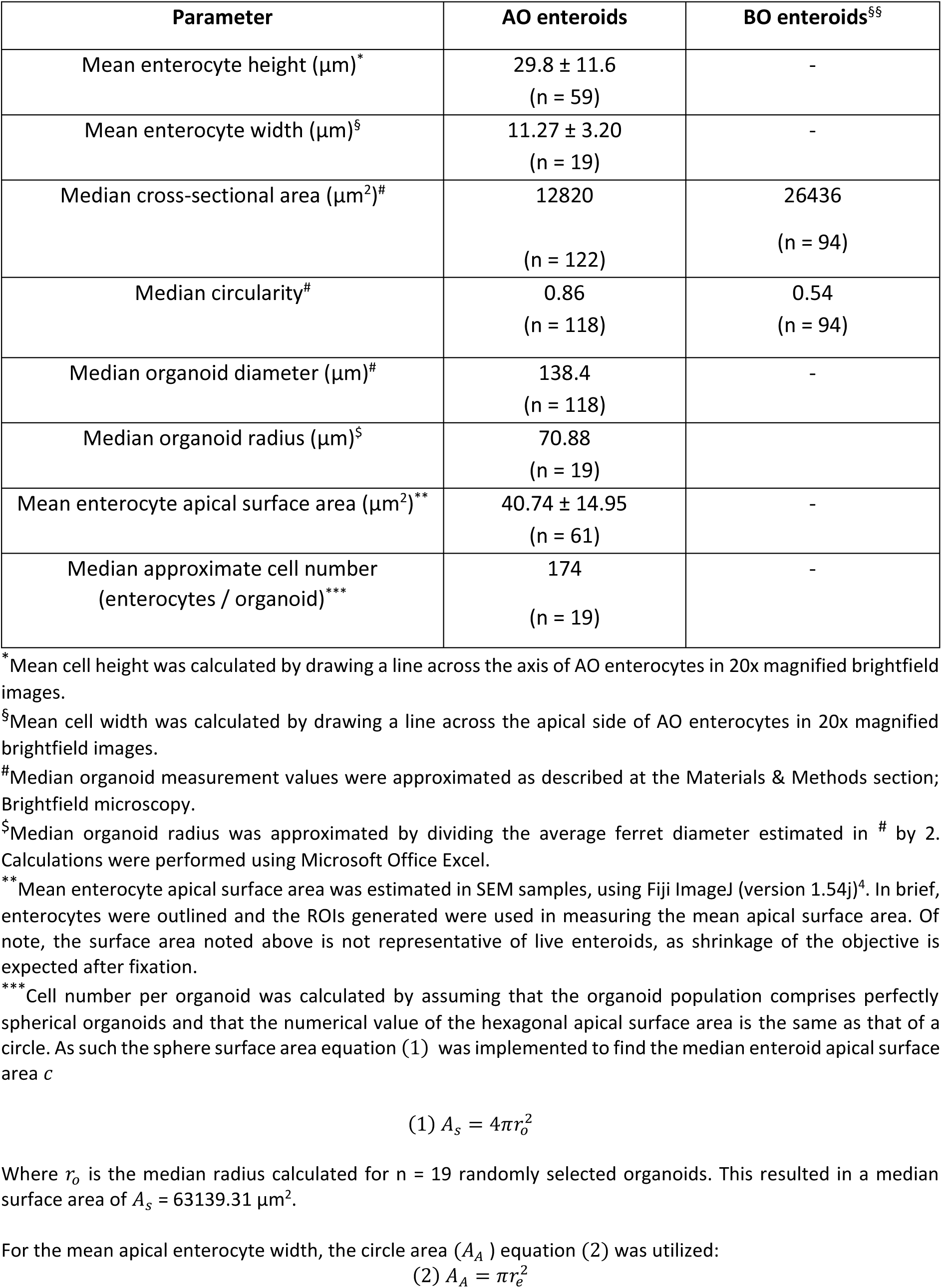
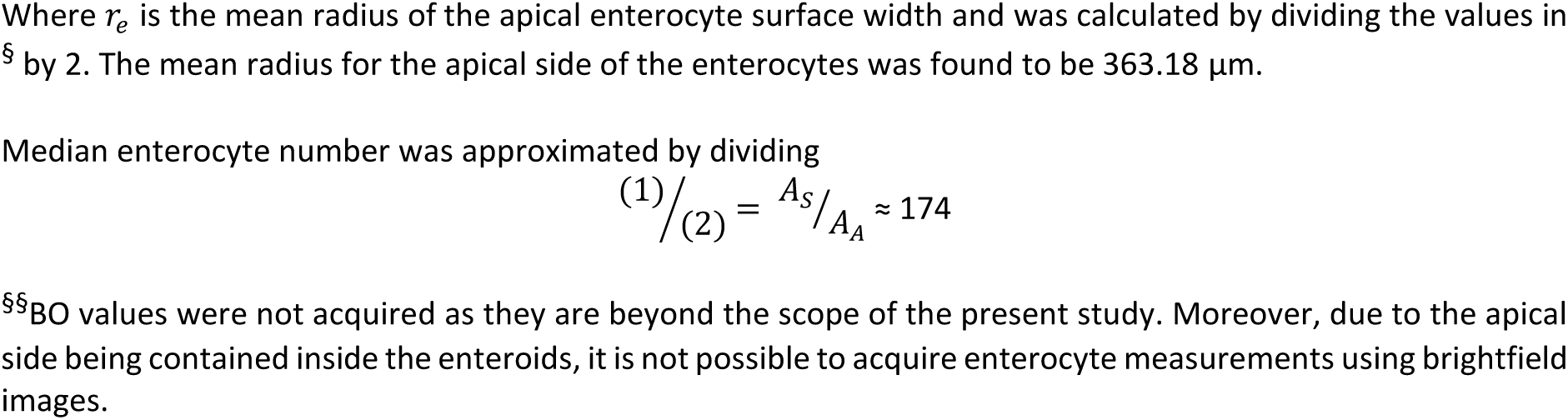
Summary of size and shape parameters for AO and BO enteroids. Analysis is based on measurements conducted on brightfield and SEM images. Data are presented as mean ± SD or median and n indicates number of enteroids analyzed.

| Parameter | AO enteroids | BO enteroids<sup>§§</sup> |
| --- | --- | --- |
| Mean enterocyte height ($\mu\text{m}$)<sup>*</sup> | 29.8 ± 11.6<br>(n = 59) | - |
| Mean enterocyte width ($\mu\text{m}$)<sup>§</sup> | 11.27 ± 3.20<br>(n = 19) | - |
| Median cross-sectional area ($\mu\text{m}^2$)<sup>#</sup> | 12820<br>(n = 122) | 26436<br>(n = 94) |
| Median circularity<sup>#</sup> | 0.86<br>(n = 118) | 0.54<br>(n = 94) |
| Median organoid diameter ($\mu\text{m}$)<sup>#</sup> | 138.4<br>(n = 118) | - |
| Median organoid radius ($\mu\text{m}$)<sup>§</sup> | 70.88<br>(n = 19) | |
| Mean enterocyte apical surface area ($\mu\text{m}^2$)<sup>**</sup> | 40.74 ± 14.95<br>(n = 61) | - |
| Median approximate cell number<br>(enterocytes / organoid)<sup>***</sup> | 174<br>(n = 19) | - |

\* Mean cell height was calculated by drawing a line across the axis of AO enterocytes in 20x magnified brightfield images.
§ Mean cell width was calculated by drawing a line across the apical side of AO enterocytes in 20x magnified brightfield images.

**Figure S2.**
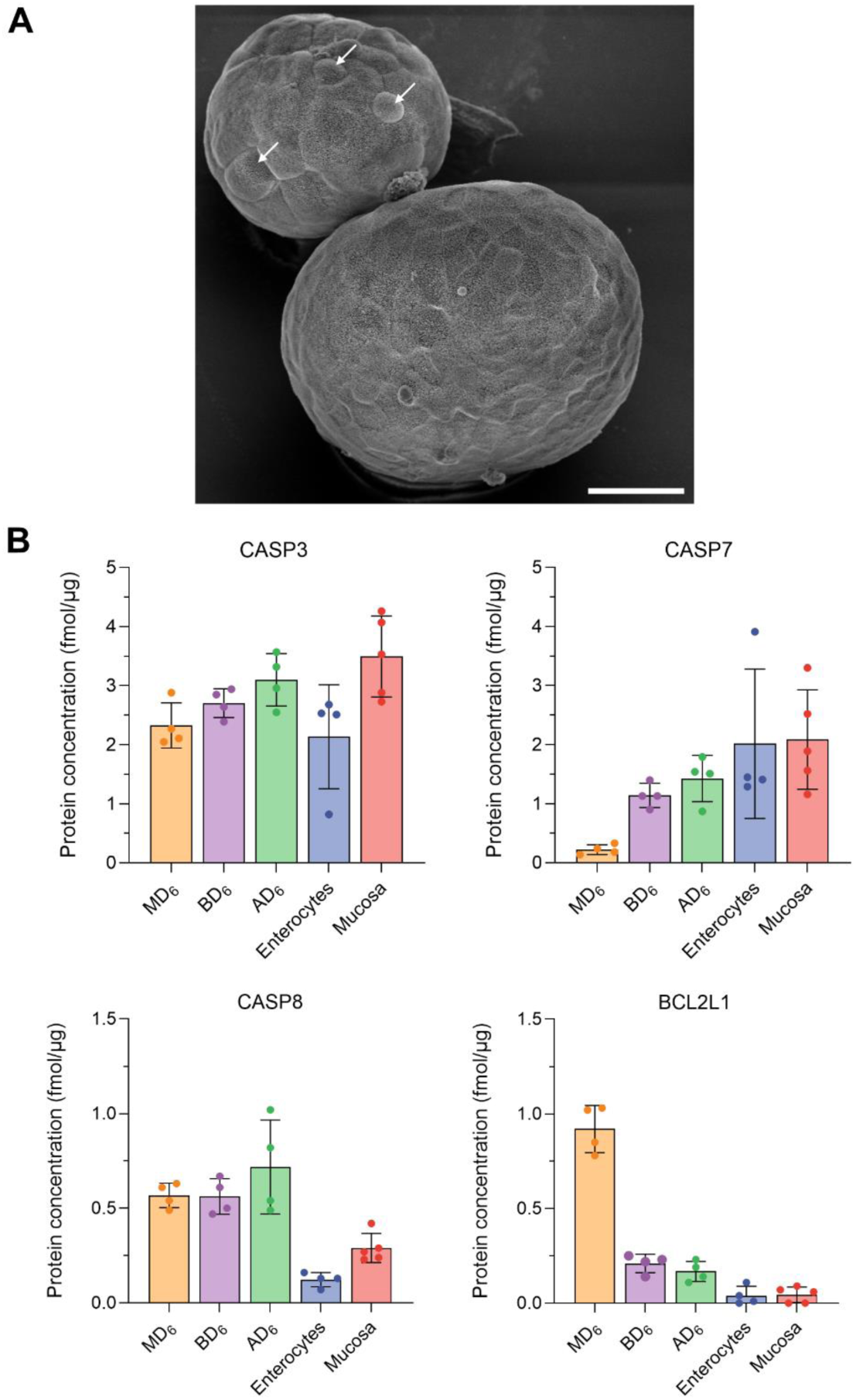
Morphology of cell extrusion from apical-out enteroids and expression of apoptosis-related proteins. **(A)** SEM image of apical-out (AO) enteroids where single enterocytes are beginning to buldge from enterocyte monolayer (as highlighted by white arrows). Scale bars, 20 µm. **(B)** Concentrations of apoptosis related proteins (CASP8, CASP3, CASP7, BCL2L1) present in different 3D enteroid formats cultured in ODM for 6 days (M: Matrigel, B: basal-out, A: apical-out), compared to primary enterocyte and jejunal mucosa *in vivo* levels. Data from four or five donors are shown. Data are presented as mean ± SD.

**Table S4.** Abundance of proteins related to microvillus structure in human enteroids, primary enterocytes, and jejunal mucosa. Protein levels are given in fmol/µg protein. Data are presented as mean (min – max) and N indicates the number of donors for which the protein was quantified.

|  | MD6 | BD6 | AD6 | Primary enterocytes | Jejunal mucosa |
| --- | --- | --- | --- | --- | --- |
|  | Mean<br>(min - max)<br>N | Mean<br>(min - max)<br>N | Mean<br>(min - max)<br>N | Mean<br>(min - max)<br>N | Mean<br>(min - max)<br>N |
| VIL1 | 31.76<br>(29.88 - 35.76)<br>4 | 42.22<br>(35.13 - 45.12)<br>4 | 46.28<br>(44.98 - 60.71)<br>4 | 37.06<br>(31.63 - 42.48)<br>4 | 54.73<br>(47.43 - 60.42)<br>5 |
| ESPN | 0.19<br>(0.1 - 0.29)<br>4 | 0.21<br>(0.09 - 0.23)<br>4 | 0.28<br>(0.24 - 0.33)<br>4 | 0.38<br>(0.24 - 0.51)<br>4 | 0.44<br>(0.34 - 0.48)<br>5 |
| EPS8 | 0.66<br>(0.62 - 0.81)<br>4 | 1.3<br>(1.14 - 1.58)<br>4 | 1.07<br>(1.05 - 1.2)<br>4 | 0.62<br>(0.6 - 0.64)<br>4 | 0.87<br>(0.76 - 0.93)<br>5 |
| EZR | 24.78<br>(21.4 - 27.03)<br>4 | 33.9<br>(26.51 - 40.4)<br>4 | 44.03<br>(34.76 - 44.33)<br>4 | 22.26<br>(17.37 - 27.14)<br>4 | 32.23<br>(32.14 - 42.01)<br>5 |
| PDZK1 | 0.09<br>(0.02 - 0.12)<br>4 | 1.43<br>(0.48 - 4.12)<br>4 | 5.44<br>(1.78 - 6.89)<br>4 | 3.03<br>(2.86 - 3.19)<br>4 | 4.11<br>(3.86 - 4.82)<br>5 |
| MYO1A | 0.26<br>(0.19 - 0.29)<br>4 | 2.4<br>(2.02 - 2.65)<br>4 | 3.37<br>(2.51 - 5.69)<br>4 | 8.98<br>(8.38 - 9.58)<br>4 | 14.48<br>(11.13 - 16.57)<br>5 |
| ANKS4B | 0.09<br>(0.01 - 0.1)<br>4 | 0.45<br>(0.36 - 0.61)<br>4 | 0.42<br>(0.4 - 0.85)<br>4 | 1.31<br>(1.27 - 1.34)<br>4 | 0.93<br>(0.81 - 1.03)<br>5 |
| CDHR2 | 0.02<br>(0.01 - 0.02)<br>4 | 0.45<br>(0.2 - 0.74)<br>4 | 0.7<br>(0.56 - 0.97)<br>4 | 0.11<br>(0.09 - 0.12)<br>4 | 0.97<br>(0.88 - 1.26)<br>5 |
| CDHR5 | 0.01<br>(0.01 - 0.02)<br>4 | 0.92<br>(0.53 - 2.14)<br>4 | 3.01<br>(2.11 - 3.43)<br>4 | 1.07<br>(1.03 - 1.11)<br>4 | 1.21<br>(1.08 - 1.62)<br>5 |
| MUC1 | 0.79<br>(0.35 - 1.6)<br>4 | 1.54<br>(0.5 - 3.46)<br>4 | 0.82<br>(0.47 - 1.35)<br>4 | ND | ND |
| MUC13 | 0.4<br>(0.31 - 0.49)<br>4 | 5.18<br>(3.47 - 17.49)<br>4 | 7.31<br>(5.97 - 10.63)<br>4 | 4.8<br>(4.55 - 5.04)<br>4 | 6.58<br>(5.49 - 7.86)<br>5 |
| MUC17 | 0.02<br>(0.01 - 0.03)<br>4 | 0.07<br>(0.06 - 0.08)<br>4 | 0.06<br>(0.03 - 0.08)<br>4 | 0.02<br>(0.01 - 0.02)<br>4 | 0.01<br>(0.01)<br>2 |
| FUT2 | 0.36<br>(0.02 - 0.69)<br>3 | 1.92<br>(1.65 - 2.19)<br>2 | 1.16<br>(1.01 - 1.31)<br>2 | 0.74<br>(0.74)<br>1 | 0.42<br>(0.42)<br>1 |
| FUT3 | 1.47<br>(0.32 - 2.79)<br>4 | 2.66<br>(0.54 - 5.38)<br>4 | 1.49<br>(0.22 - 2.58)<br>4 | 1.93<br>(1.6 - 2.25)<br>4 | 0.84<br>(0.15 - 1.76)<br>5 |
| FUT8 | 0.39<br>(0.28 - 0.44)<br>4 | 0.27<br>(0.16 - 0.34)<br>4 | 0.19<br>(0.1 - 0.3)<br>4 | 0.11<br>(0.08 - 0.13)<br>4 | 0.1<br>(0.09 - 0.11)<br>5 |

**Figure S3.**
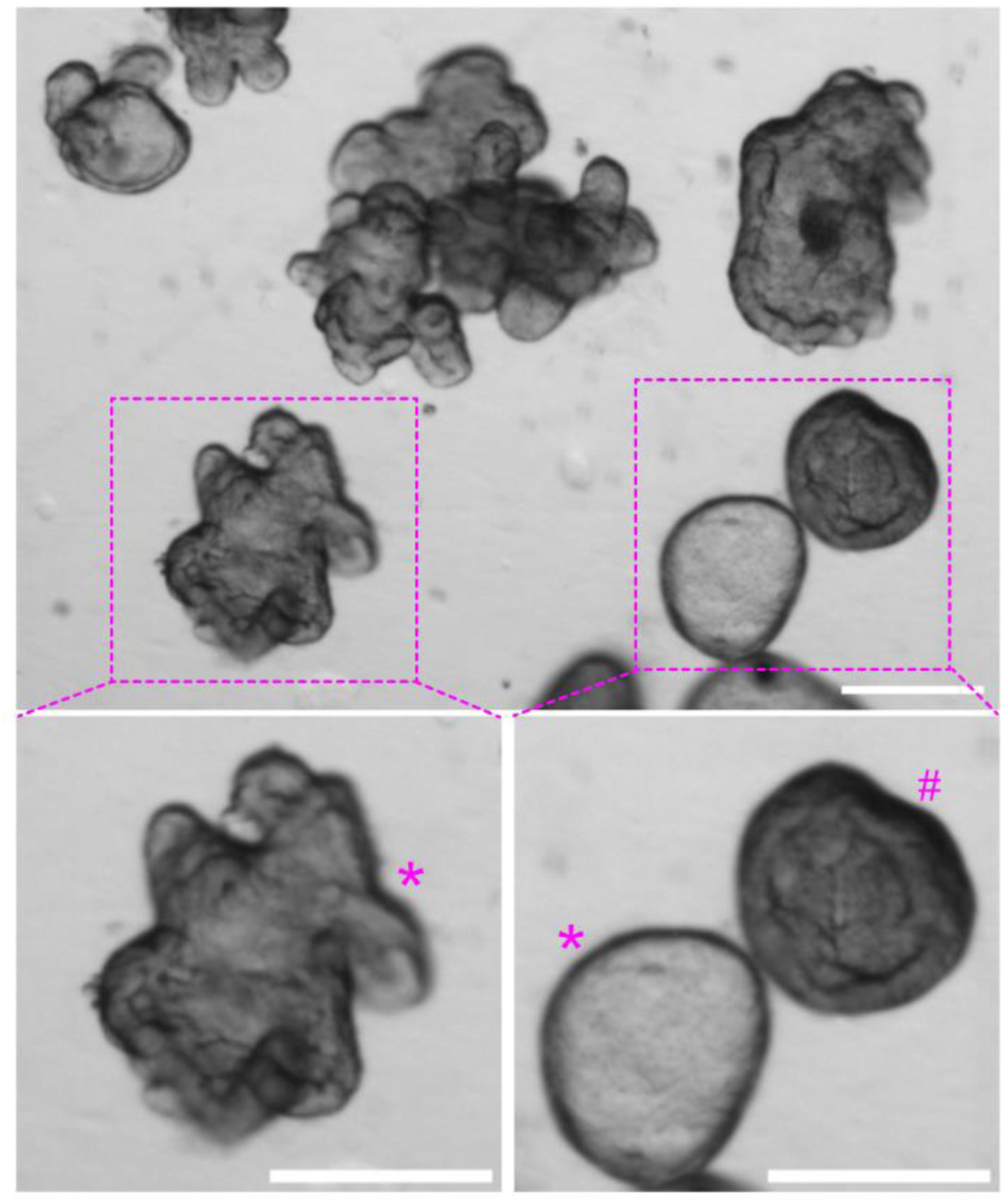
Morphology of differentiated 3D basal-out enteroids in suspension. Representative DIC micrographs of live BO enteroids at day 6 of differentiation in suspension. The image illustrates fully differentiated enteroids (marked by *) and smaller enteroids with a stem-like phenotype (marked by #). Scale bars, 200 µm.

**Figure S4.**
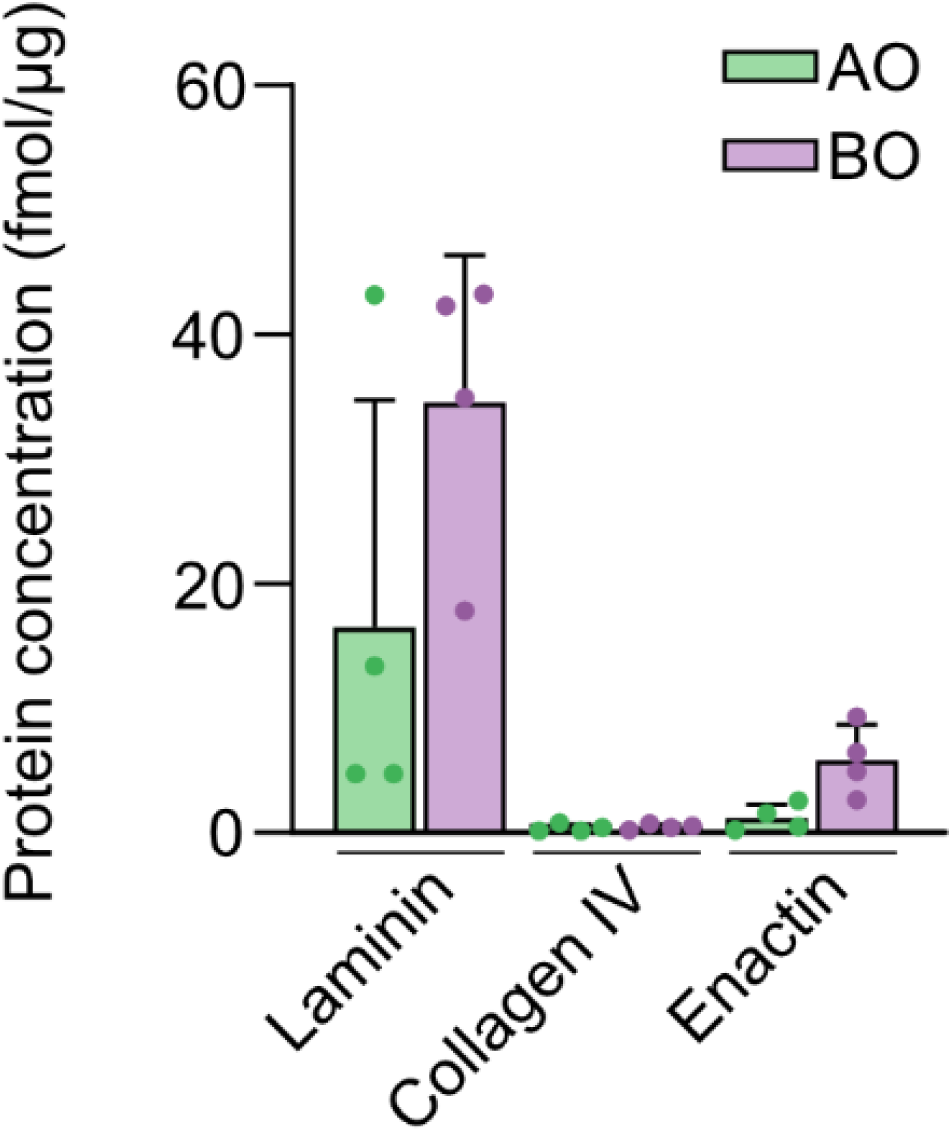
Concentrations of Matrigel proteins in AO (AD6) and BO (BD6) organoids in suspension culture. Data from four donors are shown. Data are presented as mean + SD.

**Table S5.** Abundance of tight junction, adherens junction and desmosomal proteins in human 3D enteroids, primary enterocytes, and jejunal mucosa. Protein levels are given in fmol/µg protein. Data are presented as mean (min – max) and N indicates the number of donors for which the protein was quantified. ND refers to not detected.

|  | MD6 | BD6 | AD6 | Primary enterocytes | Jejunal mucosa |
| --- | --- | --- | --- | --- | --- |
|  | Mean<br>(min - max)<br>N | Mean<br>(min - max)<br>N | Mean<br>(min - max)<br>N | Mean<br>(min - max)<br>N | Mean<br>(min - max)<br>N |
| CLD2 | 1.73<br>(1.21 - 2.43)<br>4 | 0.23<br>(0.16 - 0.32)<br>4 | 0.05<br>(0.05)<br>1 | 0.04<br>(0.04)<br>1 | ND |
| CLD15 | 0.22<br>(0.11 - 0.41)<br>4 | 1.13<br>(0.79 - 1.56)<br>4 | 0.9<br>(0.74 - 0.98)<br>4 | 0.63<br>(0.52 - 0.73)<br>4 | 0.44<br>(0.38 - 0.44)<br>5 |
| CLD3 | 2.4<br>(1.94 - 2.8)<br>4 | 4.75<br>(3.98 - 5.41)<br>4 | 4.79<br>(4.53 - 5.32)<br>4 | 1.79<br>(1.76 - 1.81)<br>4 | 2.61<br>(1.88 - 2.68)<br>5 |
| CLD7 | 1.23<br>(1.15 - 1.4)<br>4 | 2.08<br>(1.94 - 2.9)<br>4 | 1.58<br>(1.47 - 2.15)<br>4 | 1.23<br>(1.08 - 1.37)<br>4 | 1.15<br>(0.99 - 1.19)<br>5 |
| OCLN | 0.35<br>(0.12 - 2.52)<br>4 | 0.39<br>(0.23 - 2.14)<br>4 | 1.26<br>(0.34 - 3.12)<br>4 | 0.43<br>(0.16 - 0.69)<br>4 | 0.17<br>(0.03 - 0.21)<br>5 |
| MARVELD2 | 0.025<br>(0.02 - 0.03)<br>2 | 0.02<br>(0.01 - 0.02)<br>4 | 0.03<br>(0.02 - 0.04)<br>4 | ND | 0.035<br>(0.01 - 0.06)<br>2 |
| CDH1 | 2.8<br>(2.16 - 3.1)<br>4 | 4.11<br>(3.73 - 4.26)<br>4 | 4.17<br>(3.71 - 4.5)<br>4 | 2.91<br>(2.81 - 3)<br>4 | 2.88<br>(2.84 - 3.57)<br>5 |
| CTNND1 | 4.44<br>(4.18 - 4.91)<br>4 | 5.43<br>(4.88 - 6)<br>4 | 5.49<br>(4.21 - 6.25)<br>4 | 5.54<br>(5.41 - 5.67)<br>4 | 5.24<br>(4.63 - 5.6)<br>5 |
| VCL | 9.46<br>(8.59 - 12.1)<br>4 | 8.39<br>(7.57 - 8.97)<br>4 | 9.53<br>(7.9 - 10.85)<br>4 | 4.47<br>(4.14 - 4.79)<br>4 | 8.29<br>(7.89 - 13.7)<br>5 |
| DSG2 | 5.83<br>(5.14 - 6.45)<br>4 | 4.67<br>(3.88 - 6.03)<br>4 | 1.68<br>(0.78 - 4.19)<br>4 | 1.82<br>(1.79 - 1.85)<br>4 | 2.23<br>(2.12 - 2.38)<br>5 |
| DSC2 | 0.43<br>(0.39 - 0.45)<br>4 | 0.87<br>(0.56 - 1.15)<br>4 | 0.32<br>(0.14 - 0.77)<br>4 | 0.44<br>(0.38 - 0.5)<br>4 | 0.69<br>(0.55 - 0.69)<br>5 |
| DSP | 8.18<br>(6.56 - 8.77)<br>4 | 6.75<br>(6.43 - 8.68)<br>4 | 6.05<br>(3.99 - 8.05)<br>4 | 5.56<br>(5.25 - 5.87)<br>4 | 5.12<br>(4.15 - 5.52)<br>5 |
| PKP2 | 4.17<br>(3.37 - 5.96)<br>4 | 4.57<br>(4.13 - 4.95)<br>4 | 2.76<br>(1.59 - 3.82)<br>4 | 1.04<br>(1.02 - 1.05)<br>4 | 2.38<br>(1.94 - 2.46)<br>5 |

**Figure S5.**
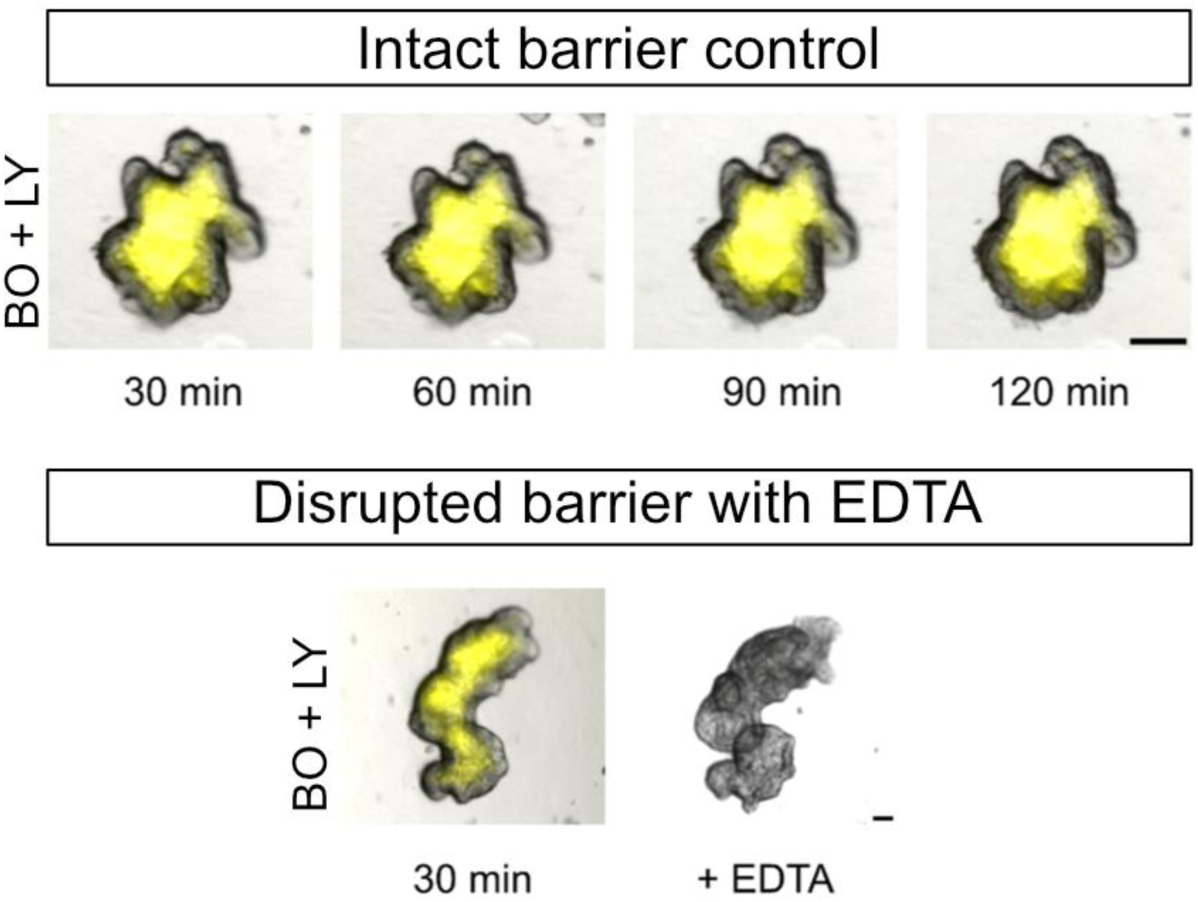
Barrier integrity assay in BO organoids in suspension. Representative images of suspended BO enteroids (n = 235) that were incubated in medium containing Lucifer yellow (LY, 100 µM) at 37°C to asses intact epithelial barrier over a time period of 120 min (upper panel). Enteroids treated with EDTA (n > 200) were used as control population (lower panel). Enteroids were imaged live by using confocal microscopy. Scale bars, 50 µm.

**Table S6.**
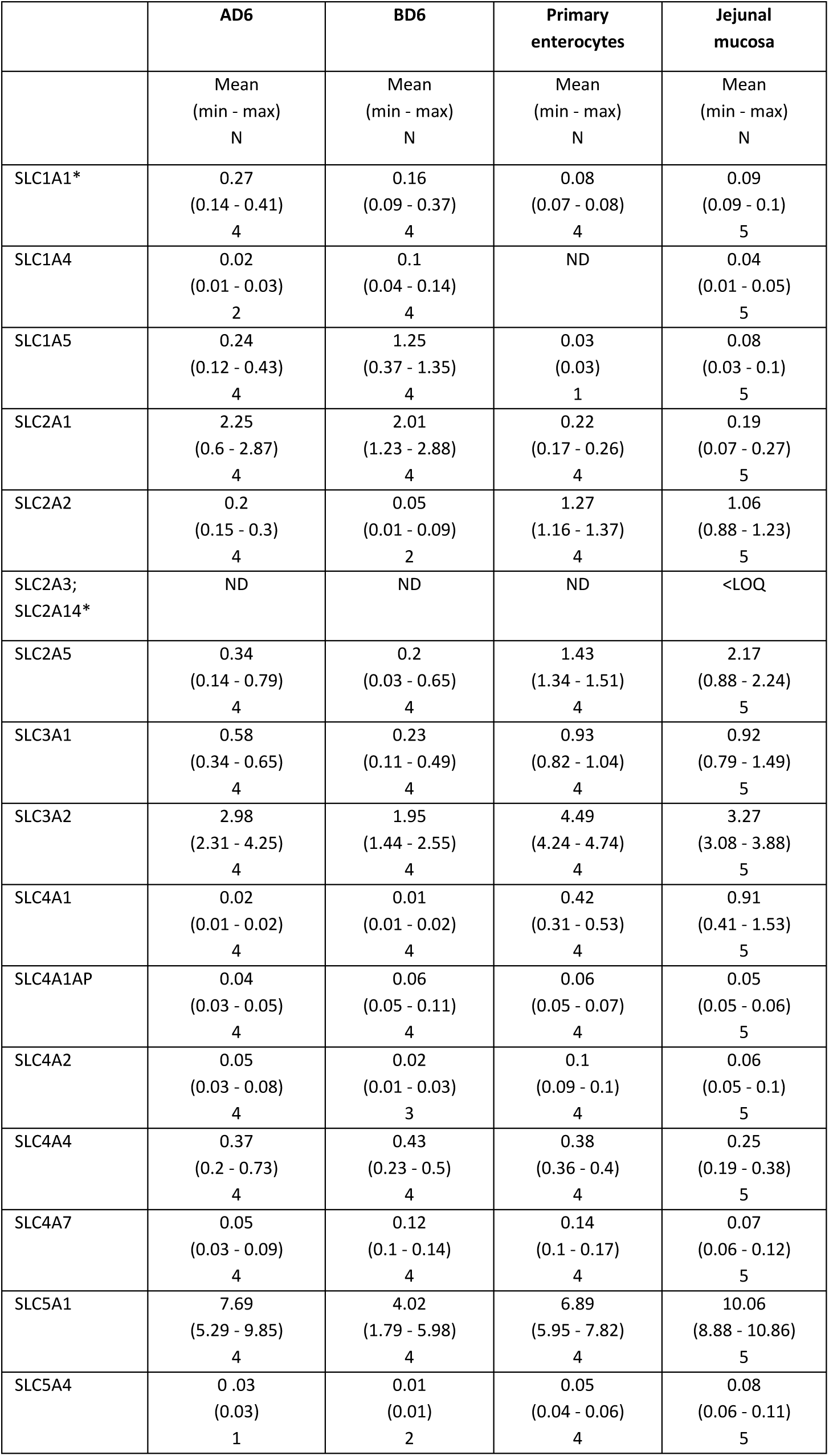
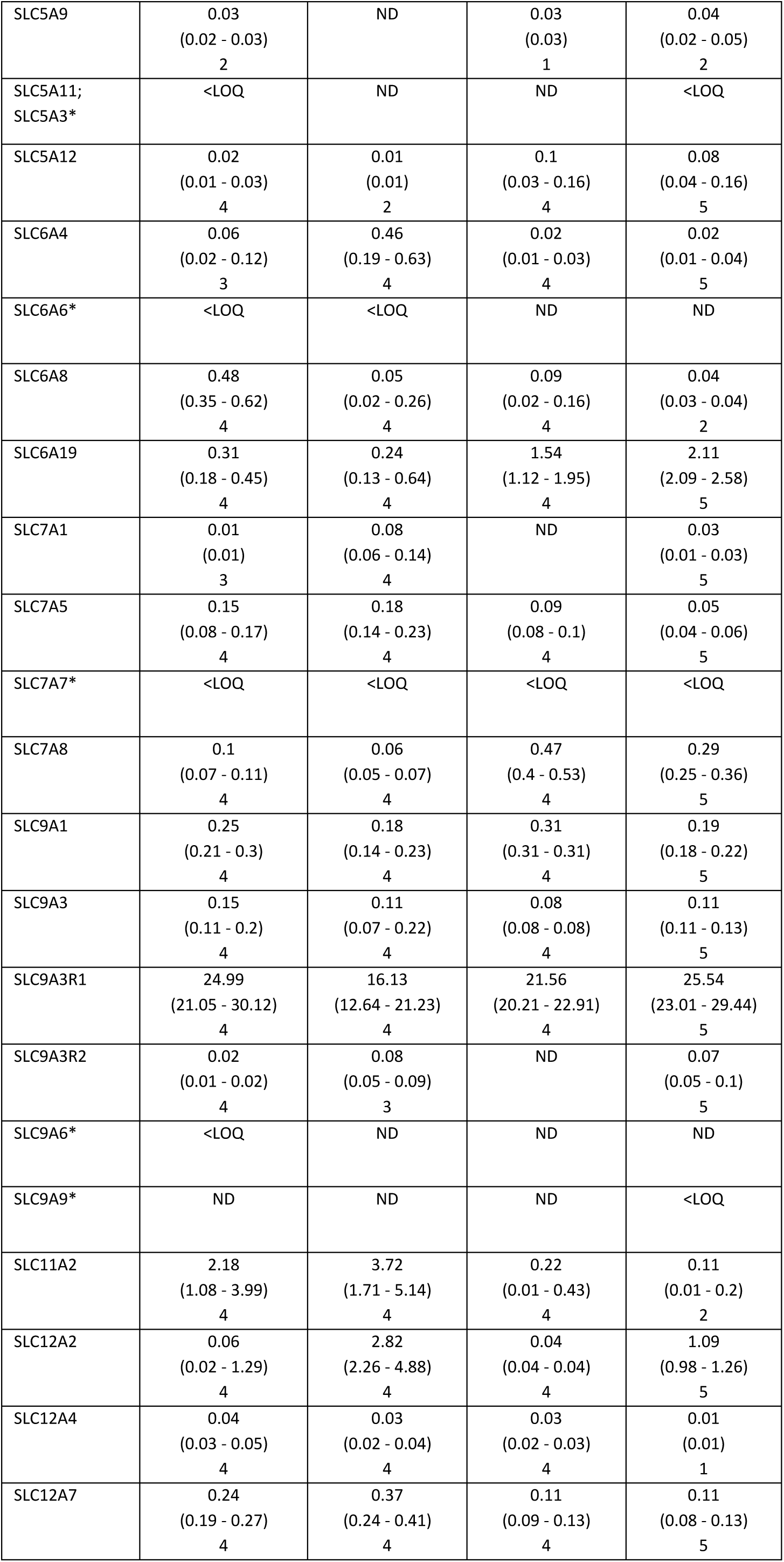
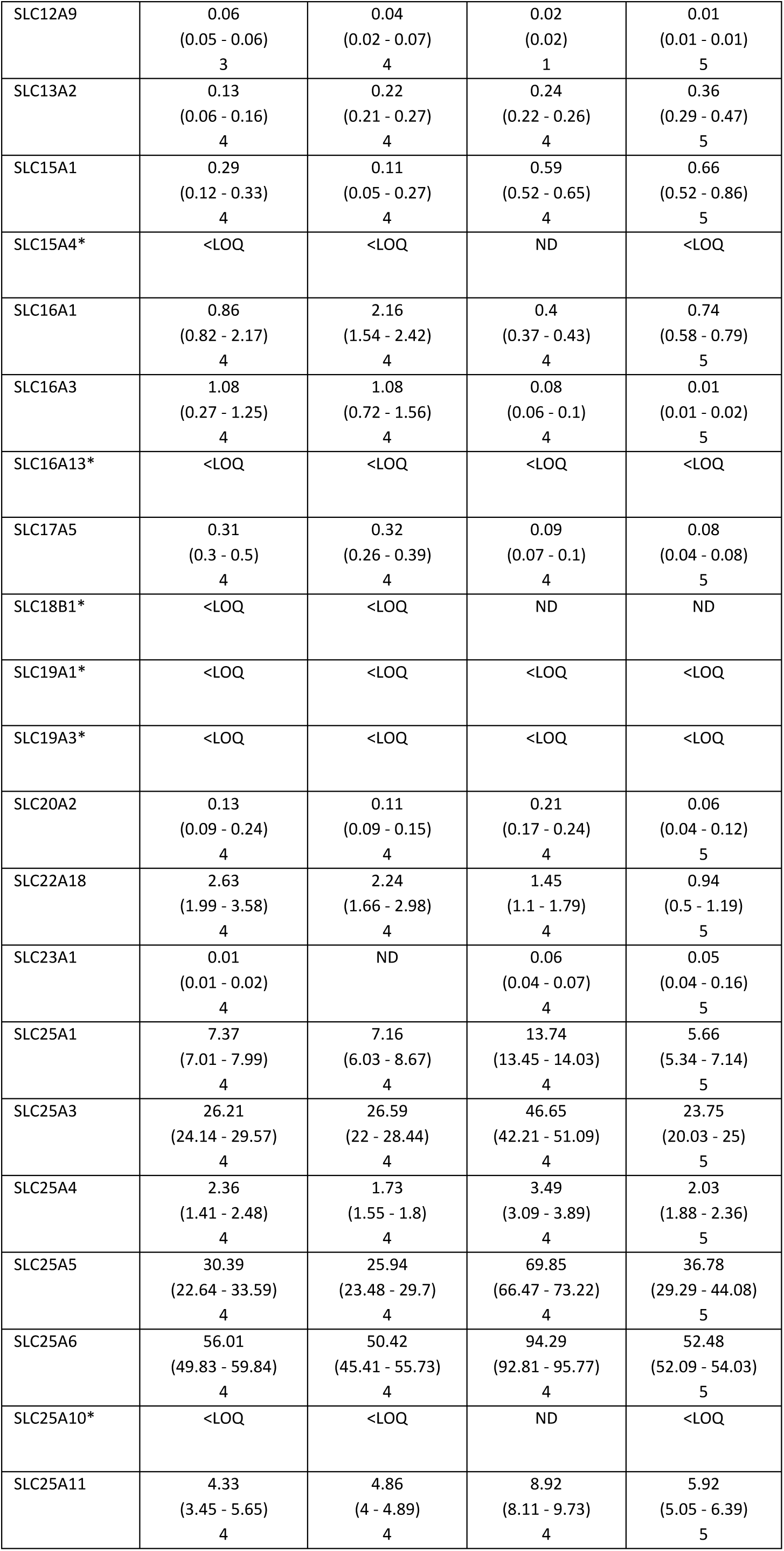
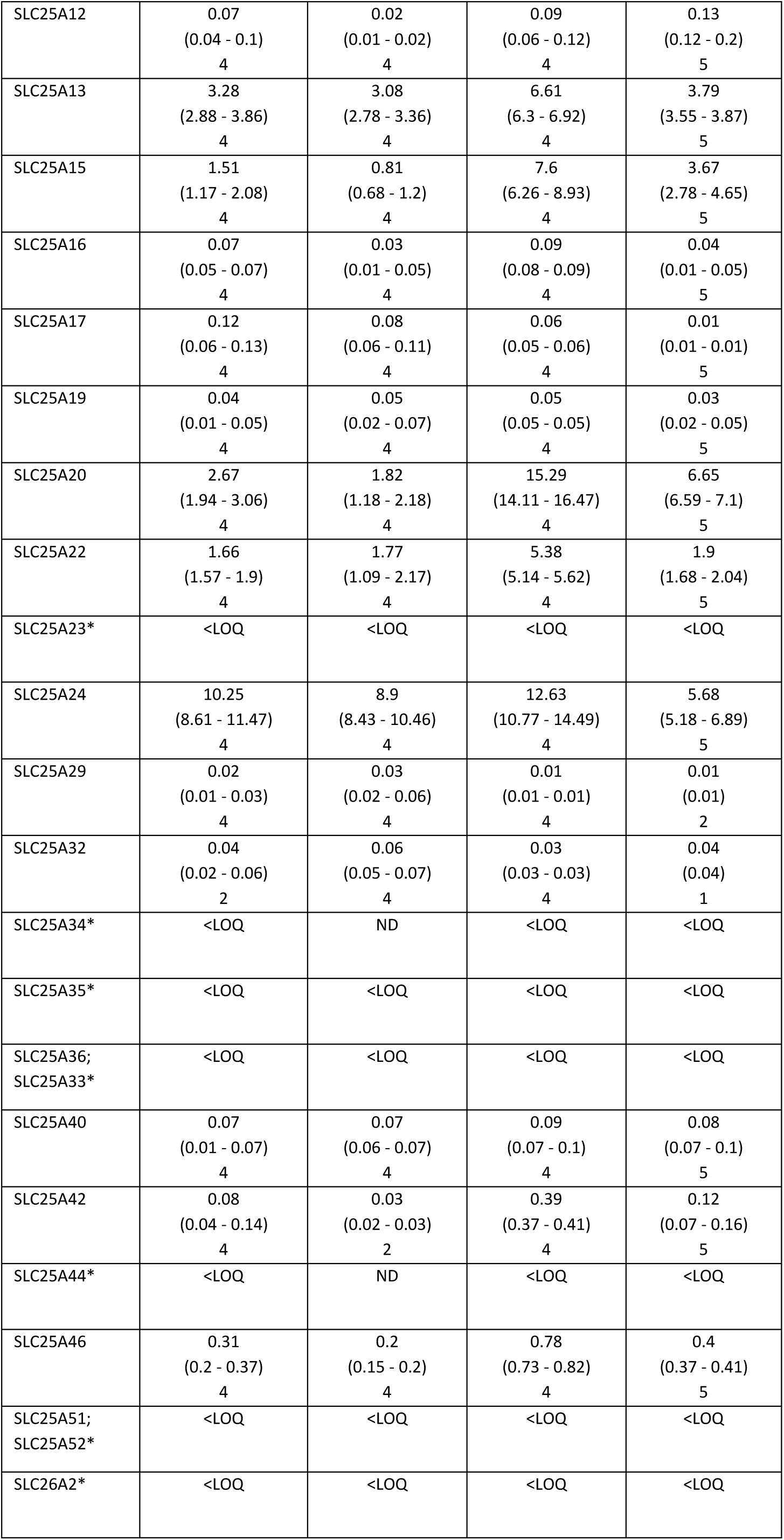
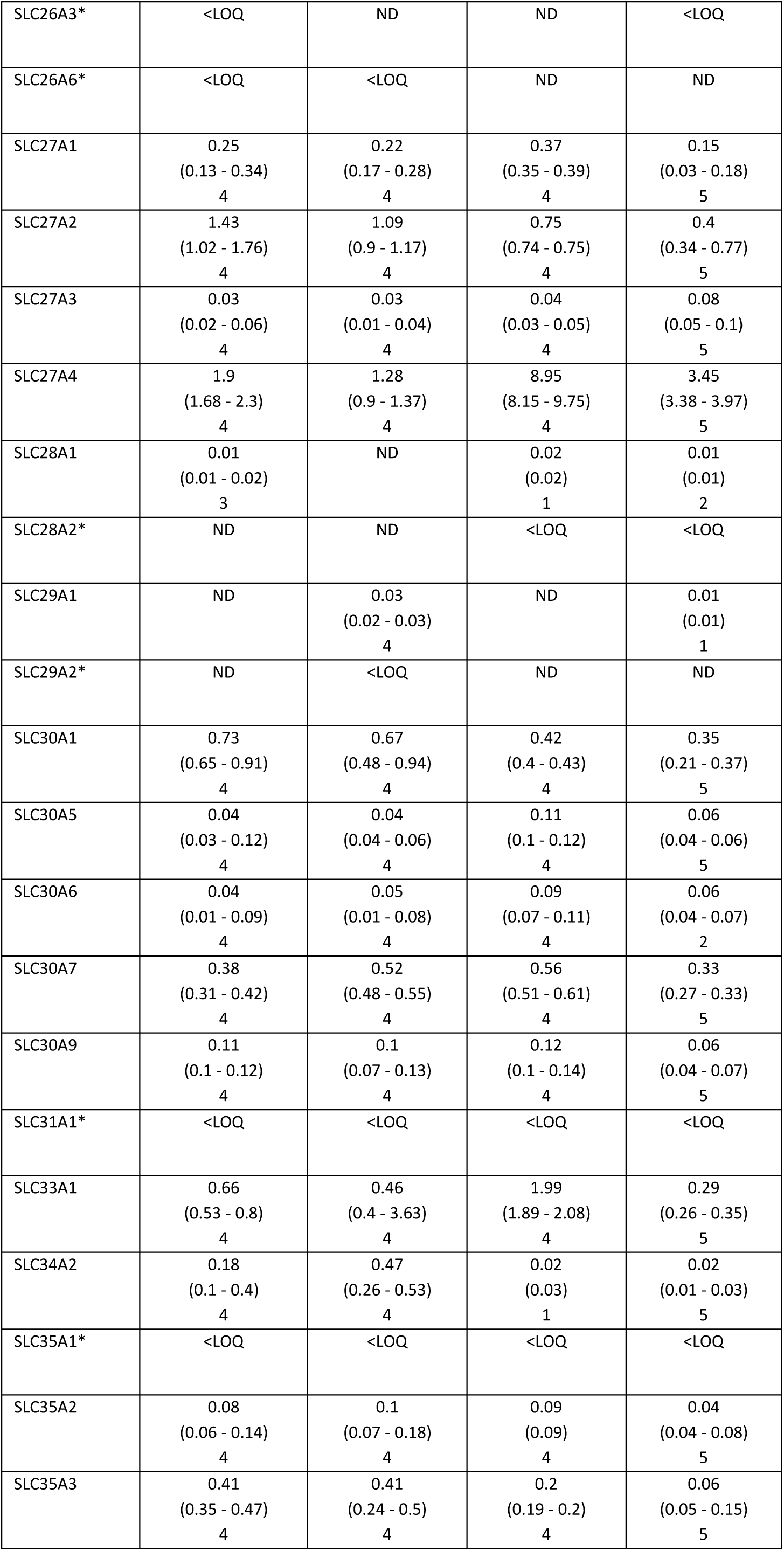
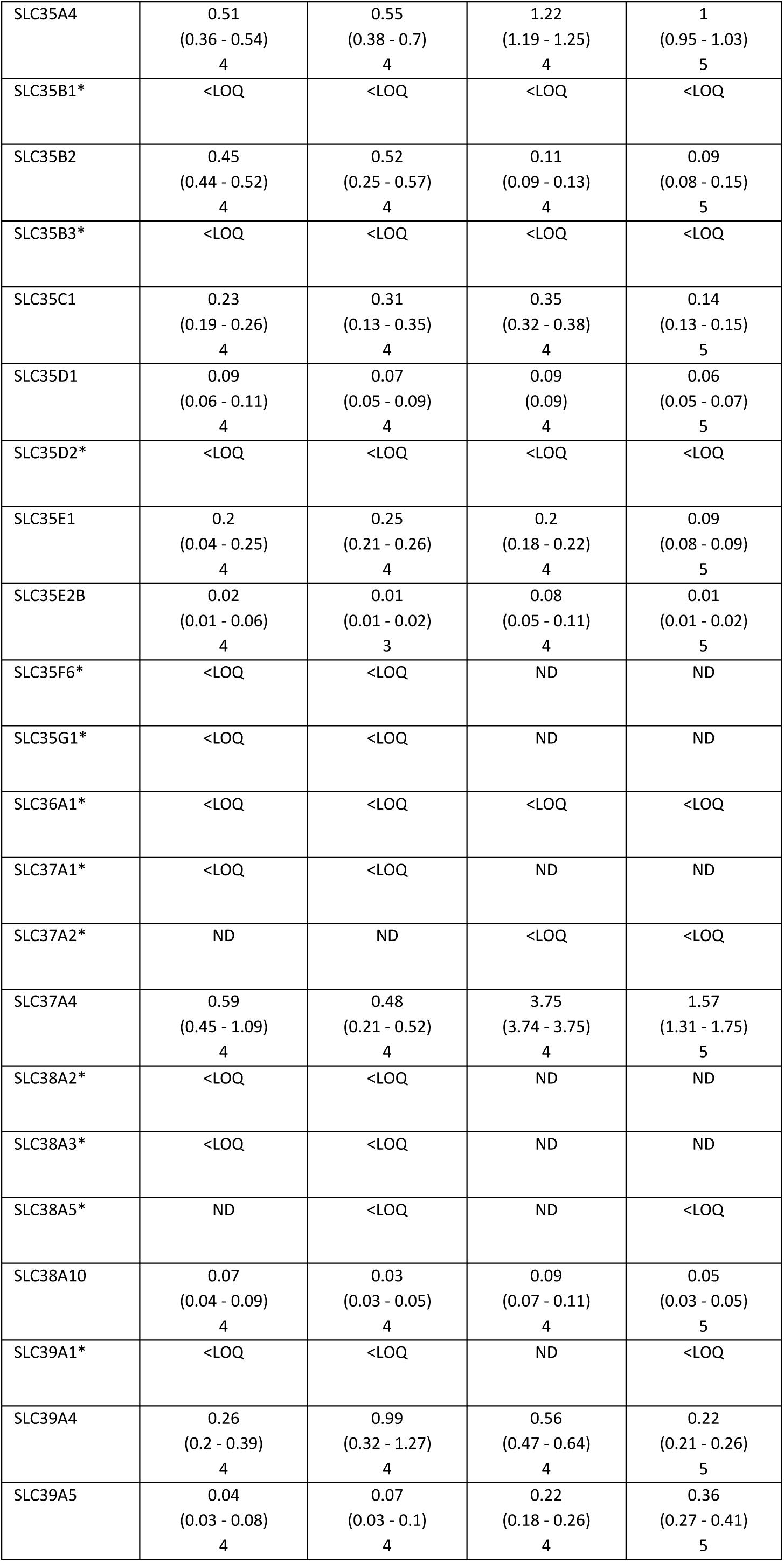
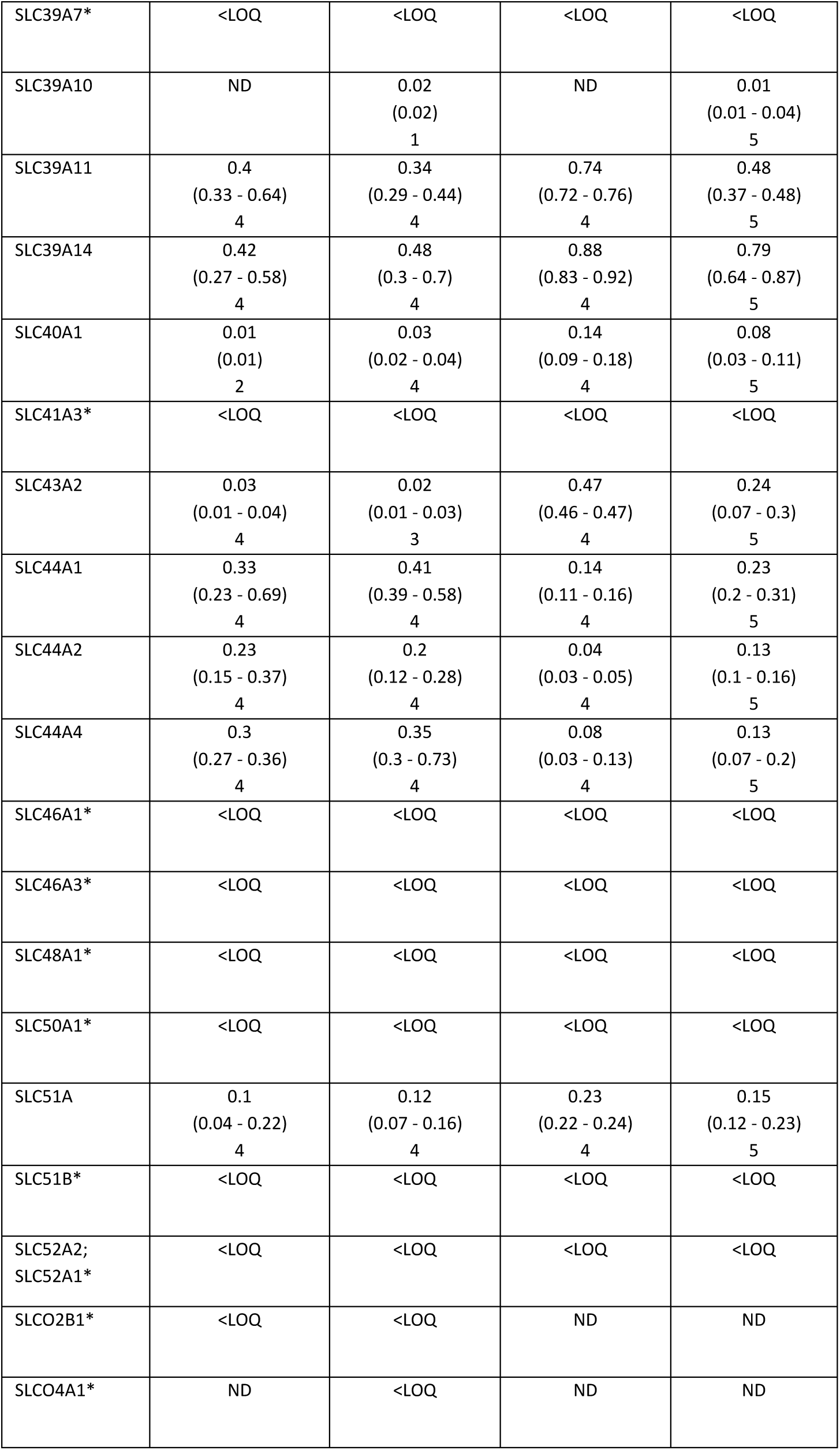
Abundance of SLC transporters in human 3D enteroids, primary enterocytes, and jejunal mucosa. Protein levels are expressed in fmol/µg protein and presented as mean (min – max). N indicates the number of donors for which the protein was quantified. An asterisk (*) denotes proteins that were identified but not quantified. ND indicates proteins that were not detected, and <LOQ refers to expression levels below the limit of quantification.

**Figure S6.**
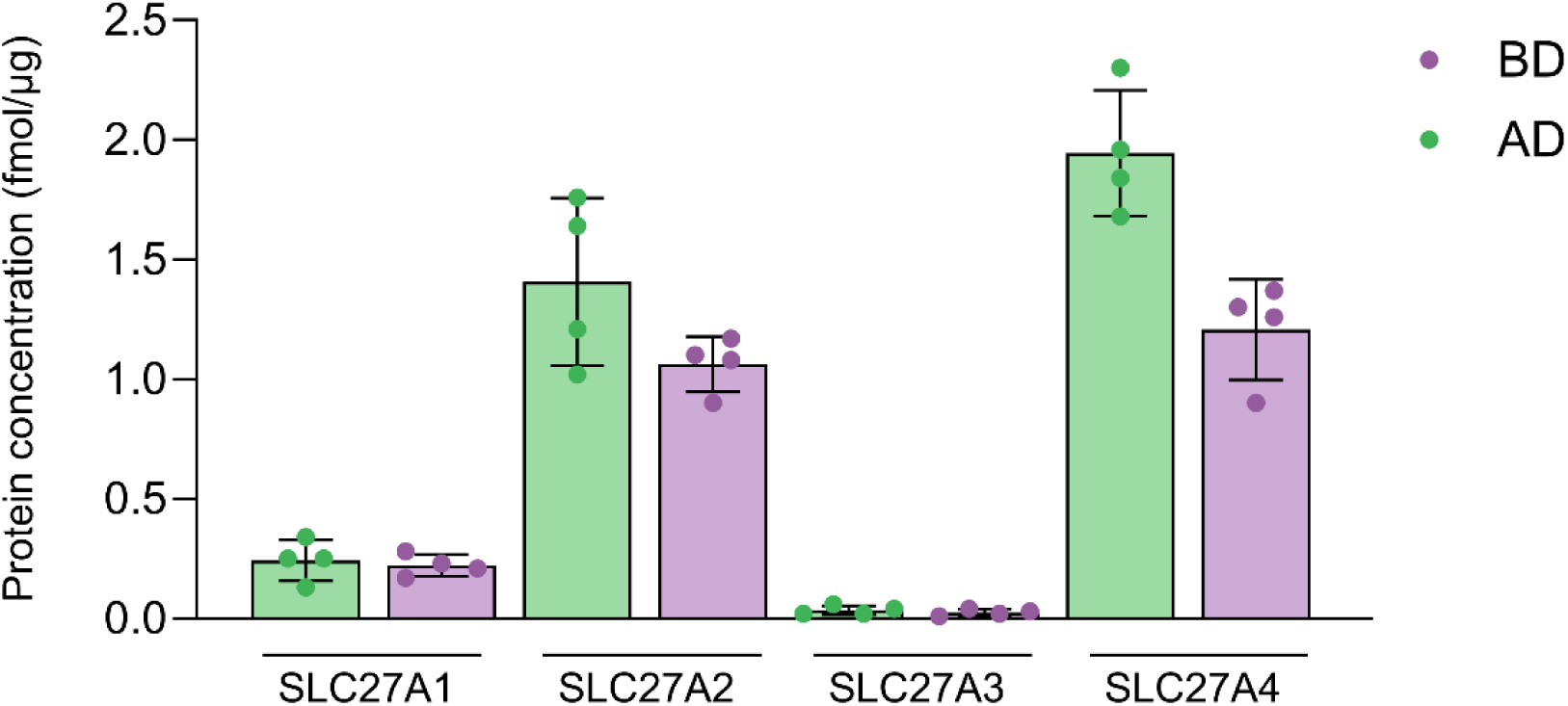
Expression of fatty acid transporting proteins (FATPs). Expression levels of FATPs as determined by global proteomics in AO (AD6) and BO (BD6) enteroids. Data from four donors are shown. Data are presented as mean ± SD.

**Figure S7.**
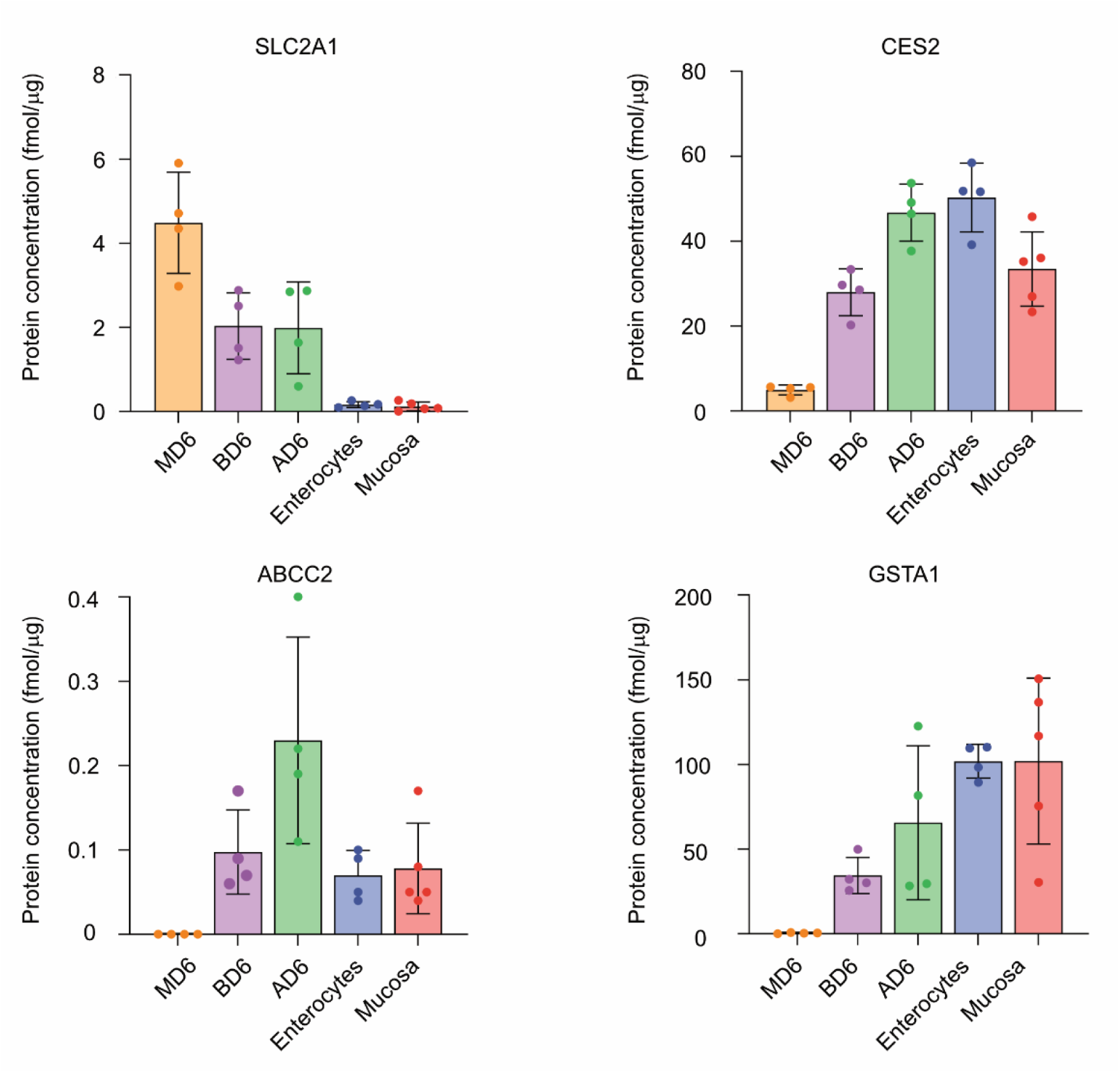
Expression of membrane transporters and metabolizing enzymes in enteroids grown in different 3D formats. Quantification of GLUT1, MRP2, CES2, and GSTA1 across 3D enteroid formats (M: Matrigel, B: basal-out, A: apical-out) cultured in ODM for 6 days, compared to primary enterocyte and jejunal mucosa *in vivo* levels. Data from four or five donors are shown. Data are presented as mean ± SD.

**Table S7.**
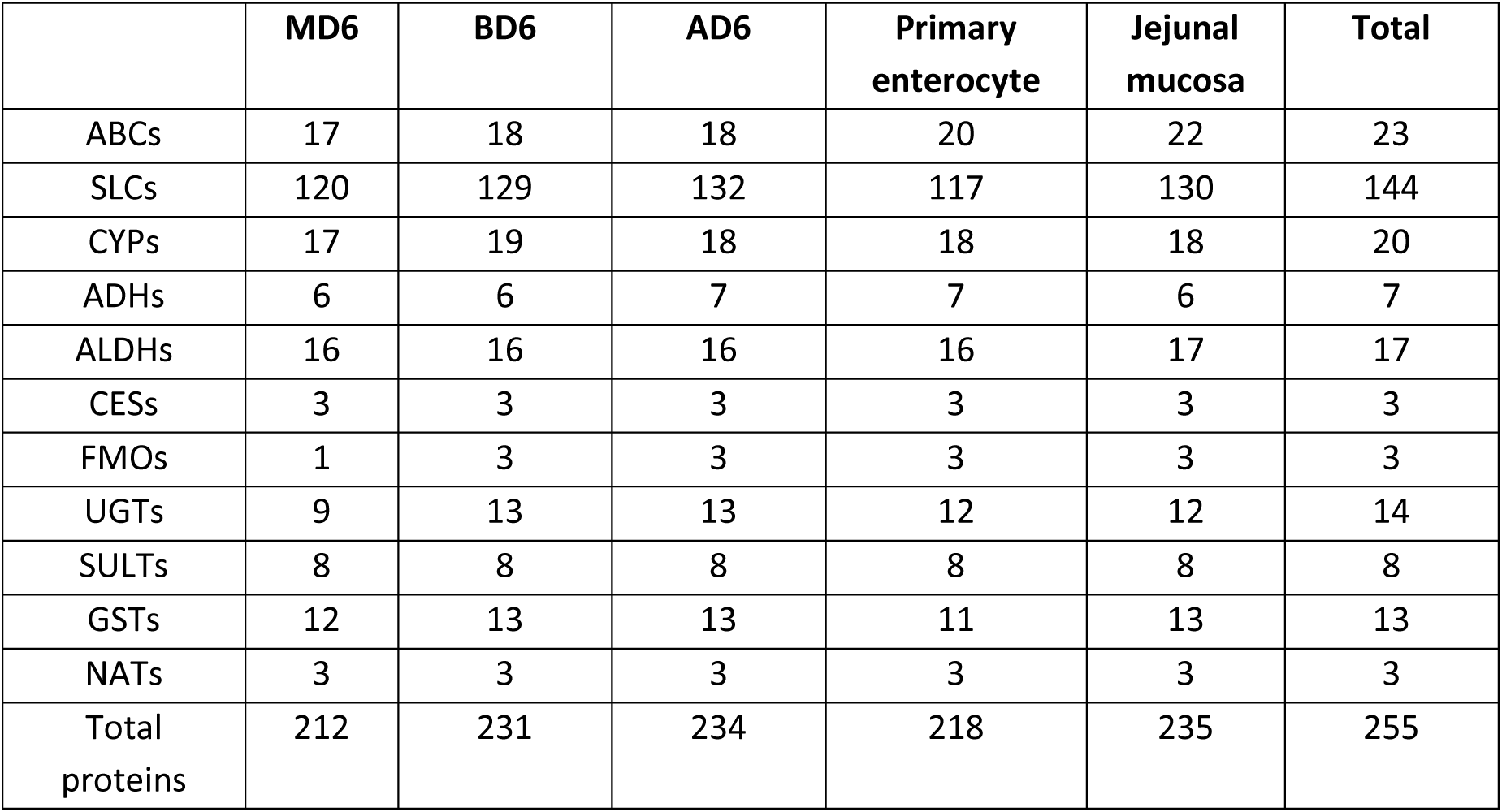
Number of quantified ADME transporters and enzymes in human enteroids, primary enterocytes, and jejunal mucosa as determined by global proteomics.

**Table S8.**
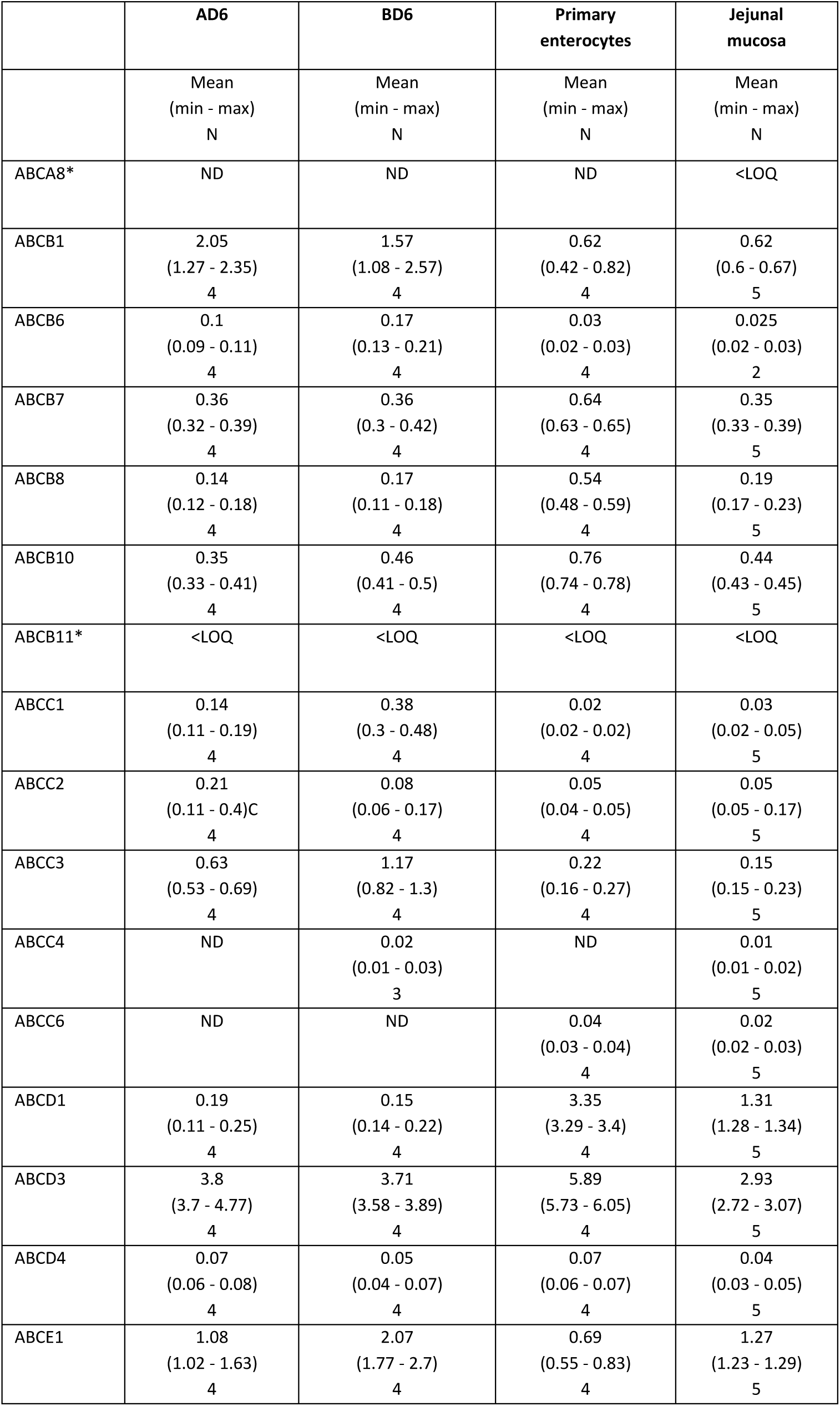
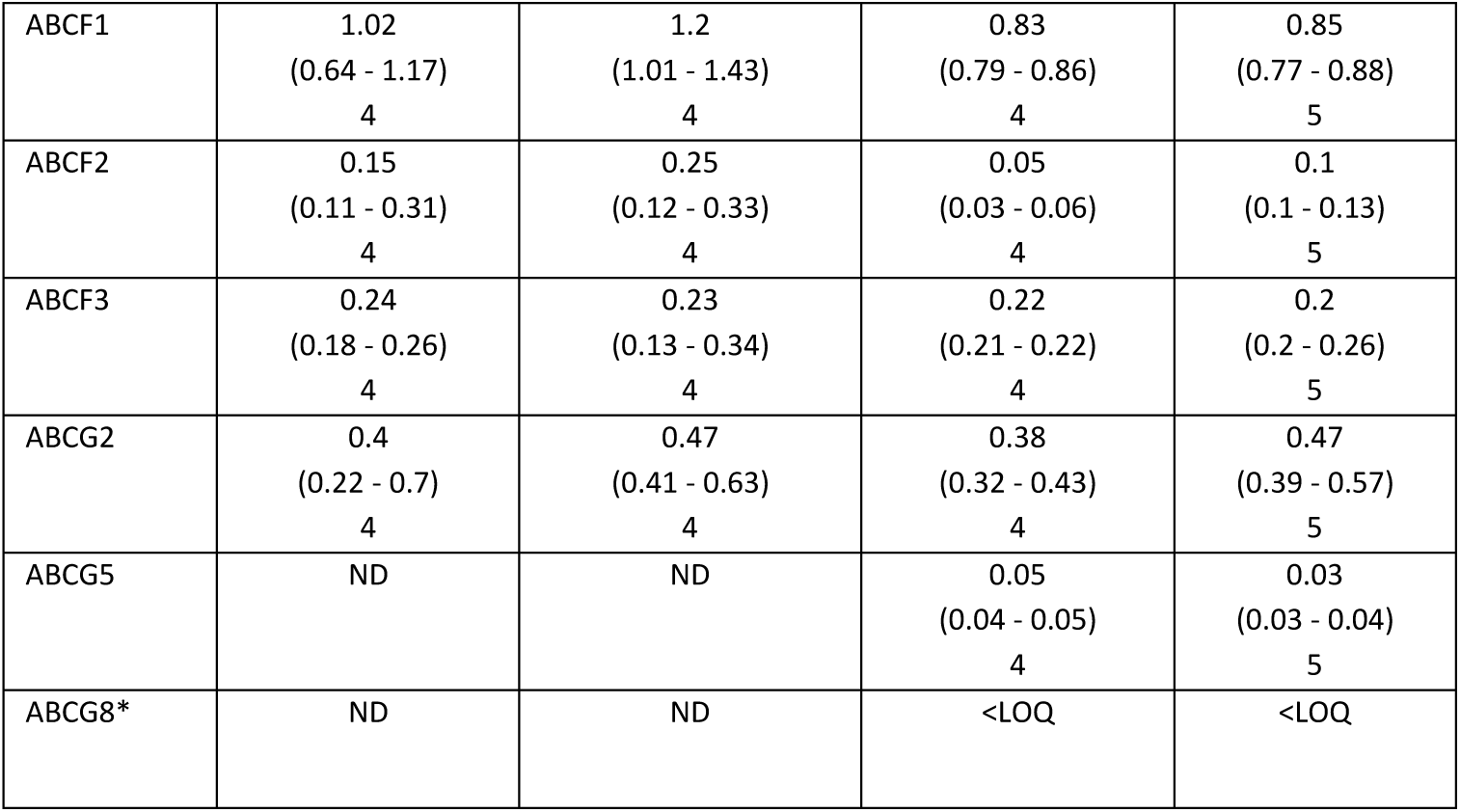
Abundance of ABC transporters in human 3D enteroids, primary enterocytes, and jejunal mucosa. Protein levels are expressed in fmol/µg protein and presented as mean (min – max). N indicates the number of donors for which the protein was quantified. An asterisk (*) denotes identified proteins. ND indicates proteins that were not detected, and <LOQ refers to expression levels below the limit of quantification.

**Figure S8.**
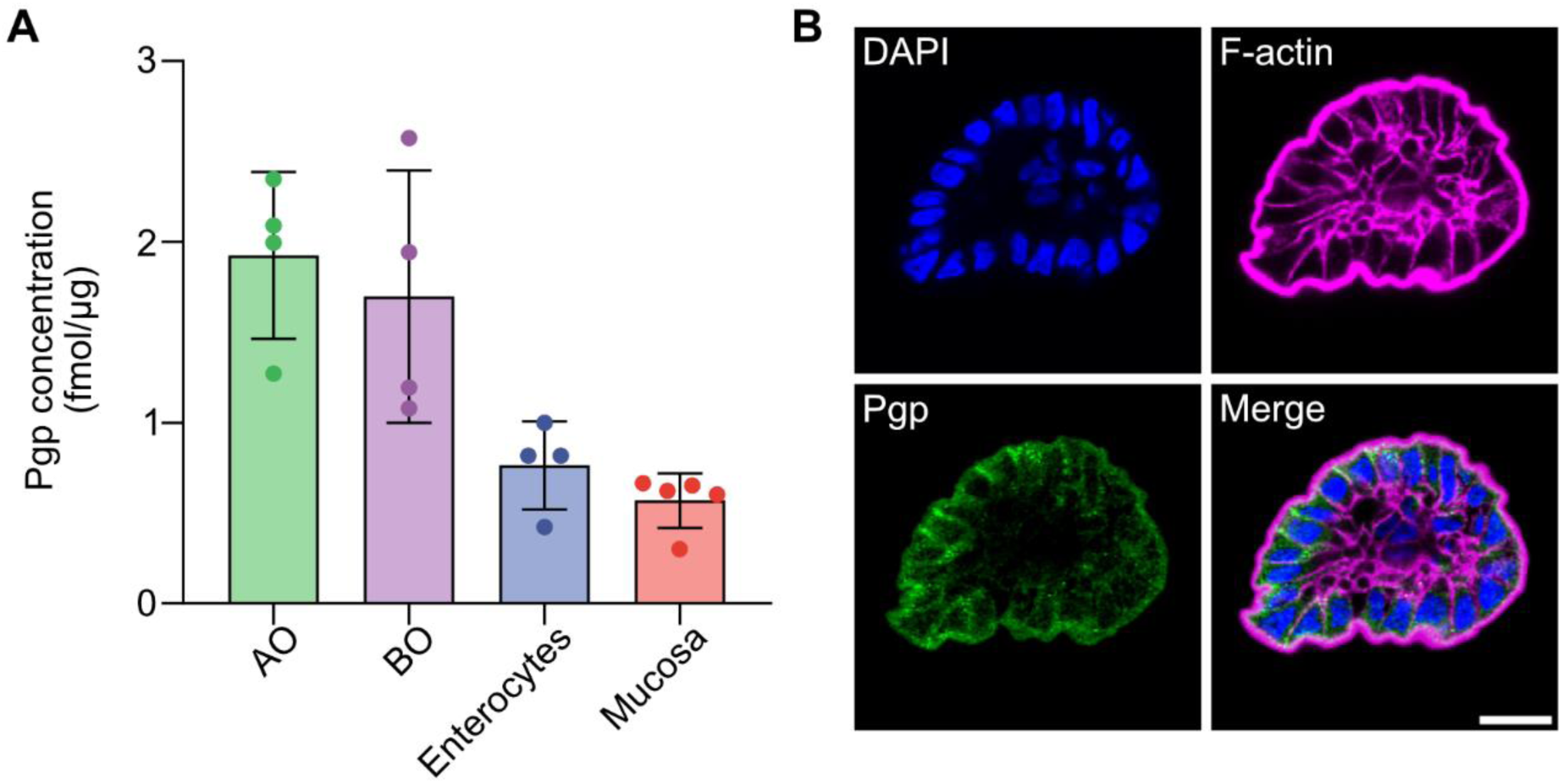
Expression of P-glycoprotein (Pgp). **(A)** Expression levels of Pgp protein as determined by global proteomics in AO and BO enteroids compared to primary enterocyte and jejunal mucosa *in vivo* levels. Data from four or five donors are shown. Data are presented as mean ± SD. **(B)** Representative confocal images following staining with DAPI (nuclei; blue), Pgp (green) and phalloidin (F-actin; magenta), highlighting expression pattern of the Pgp transporter in AO enteroids. Scale bars, 20 µm.

**Table S9.**
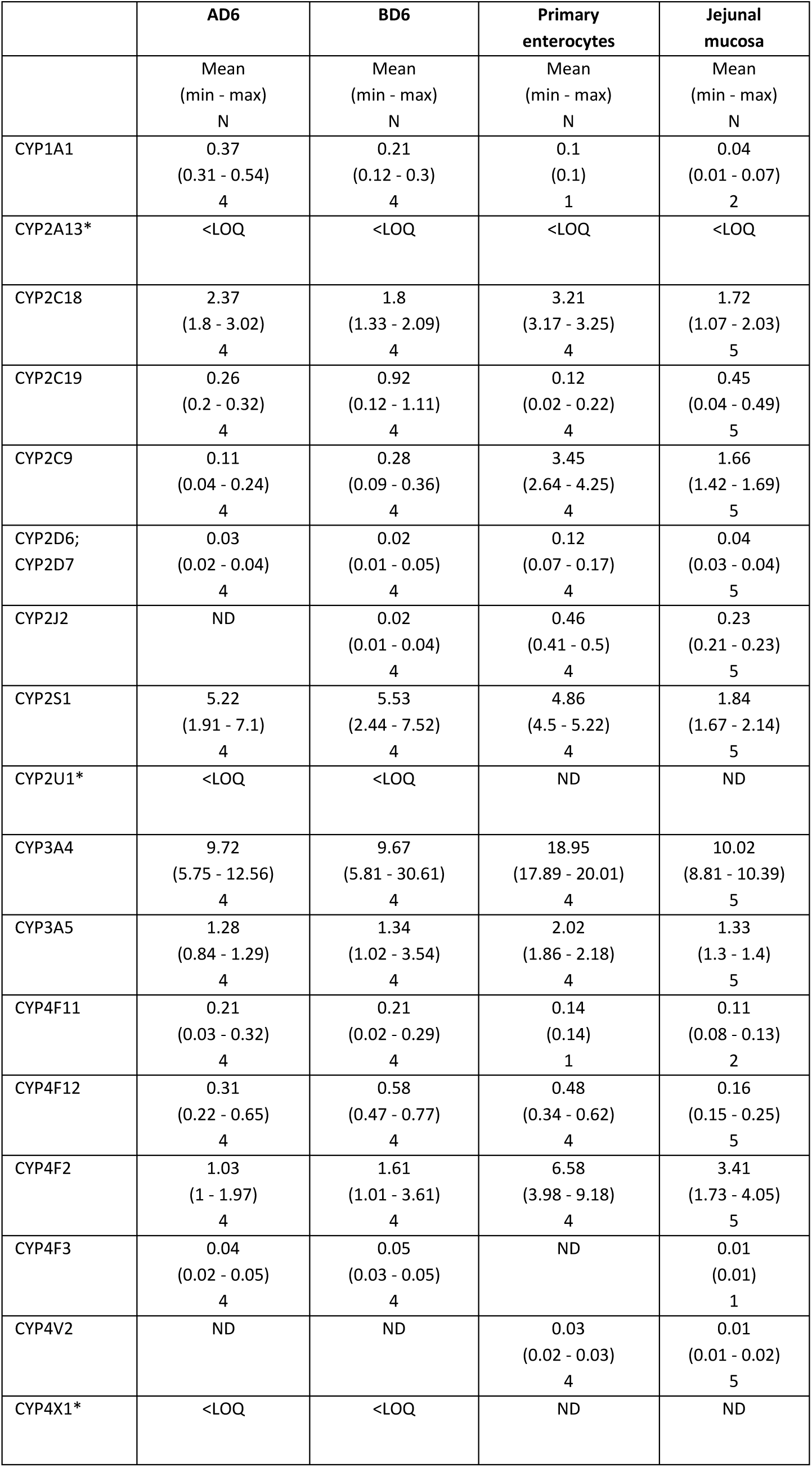
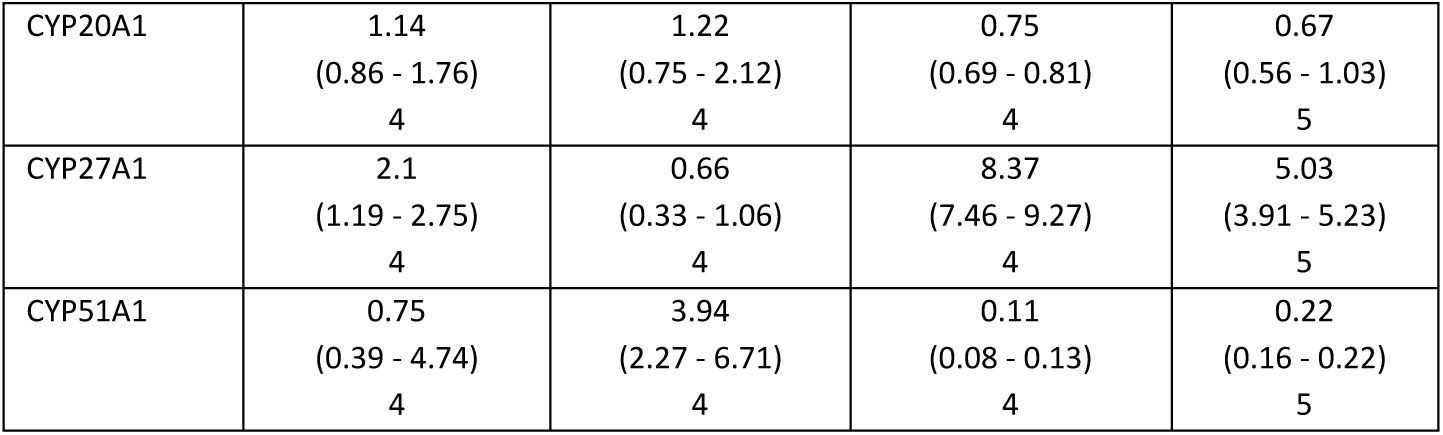
Abundance of CYP enzymes in human 3D enteroids, primary enterocytes, and jejunal mucosa. Protein levels are expressed in fmol/µg protein and presented as mean (min – max). N indicates the number of donors for which the protein was quantified. An asterisk (*) denotes identified proteins. ND indicates proteins that were not detected, and <LOQ refers to expression levels below the limit of quantification.

**Table S10.**
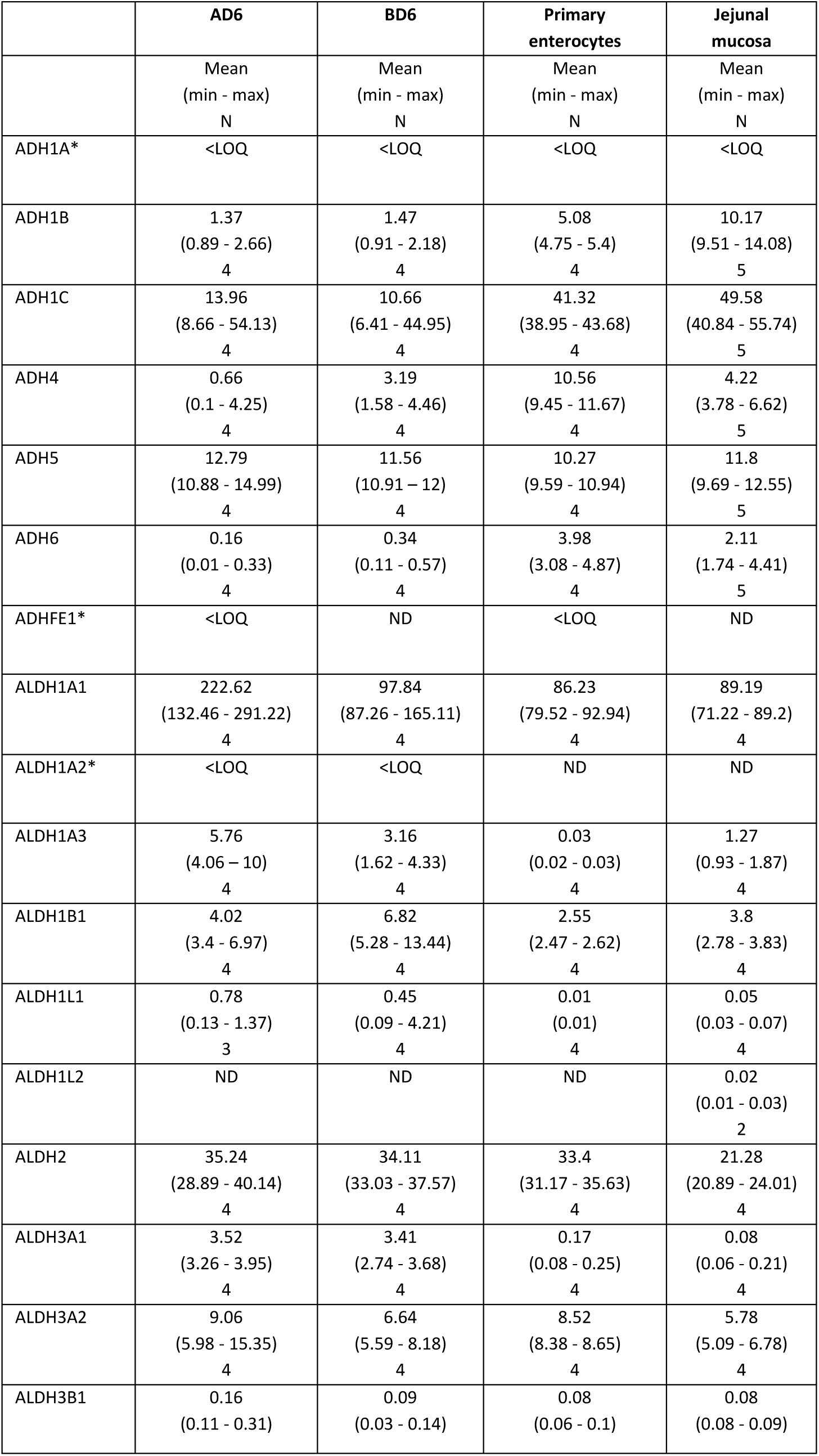
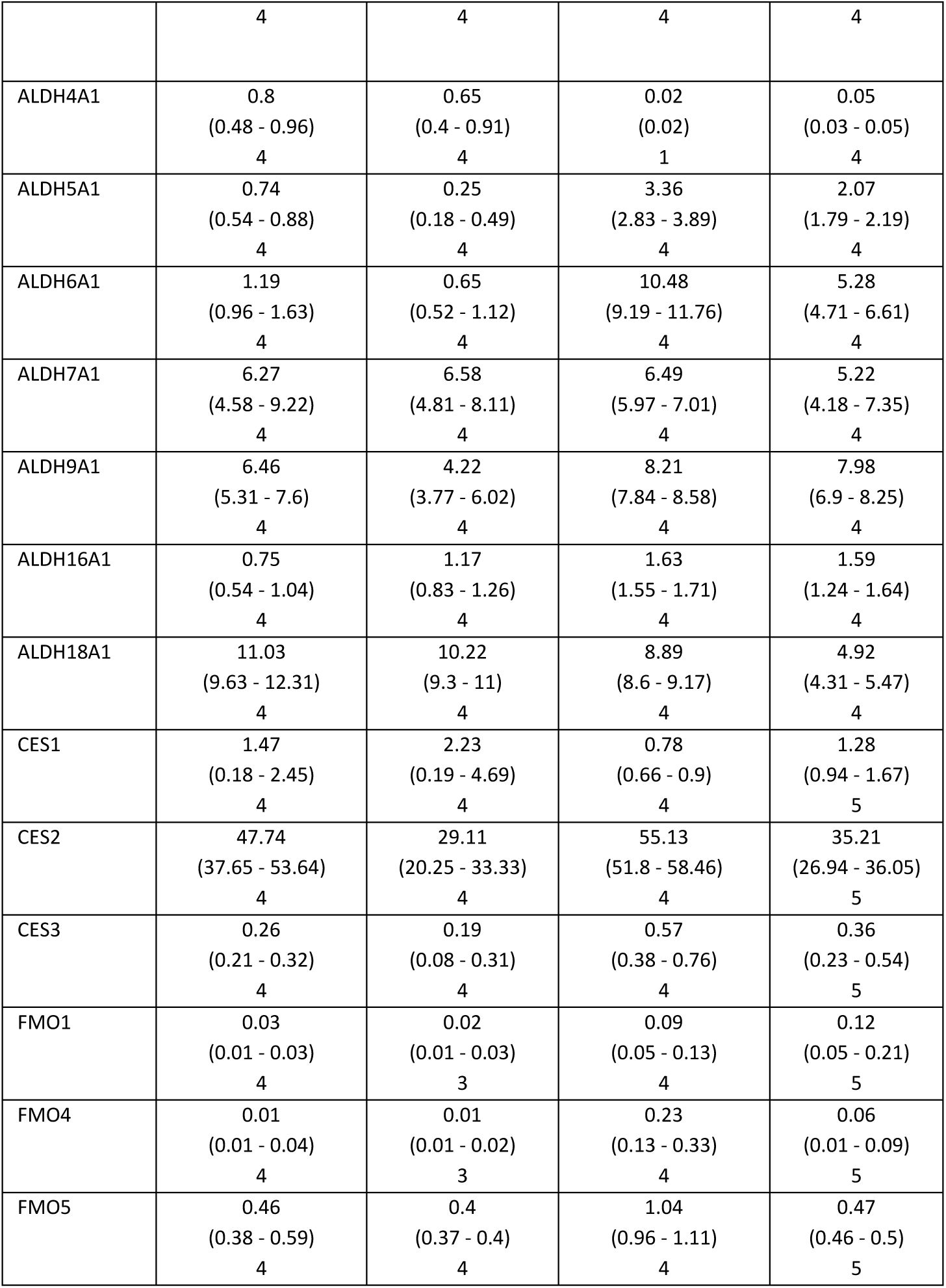
Abundance of other phase I enzymes (ADHs, ALDHs, CESs, FMOs) in human 3D enteroids, primary enterocytes, and jejunal mucosa. Protein levels are expressed in fmol/µg protein and presented as mean (min – max). N indicates the number of donors for which the protein was quantified. An asterisk (*) denotes identified proteins. ND indicates proteins that were not detected, and <LOQ refers to expression levels below the limit of quantification.

**Table S11.** Abundance of UDP-glucuronosyltransferases (UGTs) in human 3D enteroids, primary enterocytes, and jejunal mucosa. Protein levels are expressed in fmol/µg protein and presented as mean (min – max). N indicates the number of donors for which the protein was quantified. An asterisk (*) denotes identified proteins. ND indicates proteins that were not detected, and <LOQ refers to expression levels below the limit of quantification.

|  | AD6 | BD6 | Primary enterocytes | Jejunal mucosa |
| --- | --- | --- | --- | --- |
|  | Mean<br>(min – max)<br>N | Mean<br>(min – max)<br>N | Mean<br>(min – max)<br>N | Mean<br>(min – max)<br>N |
| UGT1A1 | 5.27<br>(2.53 - 7.72)<br>4 | 4.75<br>(4.15 - 5.6)<br>4 | 5.32<br>(4.97 - 5.67)<br>4 | 2.18<br>(1.39 - 2.82)<br>5 |
| UGT1A3 | 0.06<br>(0.02 - 0.1)<br>4 | 0.03<br>(0.01 - 0.11)<br>4 | 0.25<br>(0.21 - 0.28)<br>4 | 0.14<br>(0.1 - 0.14)<br>5 |
| UGT1A4 | 0.11<br>(0.03 - 0.35)<br>4 | 0.1<br>(0.01 - 1.15)<br>4 | 0.14<br>(0.14 - 0.14)<br>4 | 0.06<br>(0.06)<br>2 |
| UGT1A5* | <LOQ | <LOQ | ND | ND |
| UGT1A6 | 15.96<br>(11.65 - 20.23)<br>4 | 12.96<br>(10.24 - 16.6)<br>4 | 8.91<br>(8.23 - 9.58)<br>4 | 4.78<br>(3.5 - 5.45)<br>5 |
| UGT1A7* | <LOQ | <LOQ | ND | <LOQ |
| UGT1A8;<br>UGT1A9 | 0.18<br>(0.13 - 0.37)<br>4 | 0.22<br>(0.1 - 0.39)<br>4 | 0.08<br>(0.05 - 0.11)<br>4 | 0.13<br>(0.01 - 0.13)<br>5 |
| UGT1A10 | 5.58<br>(3.56 - 6.77)<br>4 | 4.71<br>(3.78 - 6.48)<br>4 | 2.63<br>(2.38 - 2.88)<br>4 | 0.95<br>(0.92 - 1.79)<br>5 |
| UGT2A1* | <LOQ | <LOQ | <LOQ | <LOQ |
| UGT2A3 | 0.89<br>(0.64 - 3.07)<br>4 | 0.3<br>(0.23 - 1.11)<br>4 | 9.51<br>(6.61 - 12.41)<br>4 | 6.29<br>(4.33 - 6.78)<br>5 |
| UGT2B7 | 0.32<br>(0.06 - 1.08)<br>4 | 0.05<br>(0.02 - 0.15)<br>4 | 6.08<br>(4.15 - 8)<br>4 | 3.26<br>(1.74 - 3.37)<br>5 |
| UGT2B17 | 17.75<br>(0.25 - 21.35)<br>4 | 5.7<br>(0.33 - 16.42)<br>4 | 23.71<br>(0.42 - 46.99)<br>4 | 11.31<br>(0.19 - 26.95)<br>5 |
| UGT8 | 0.23<br>(0.2 - 0.27)<br>4 | 0.36<br>(0.3 - 0.44)<br>4 | 0.12<br>(0.12 - 0.12)<br>4 | 0.11<br>(0.1 - 0.17)<br>5 |

**Table S12.** Abundance of sulfotransferases (SULTs) in human 3D enteroids, primary enterocytes, and jejunal mucosa. Protein levels are expressed in fmol/µg protein and presented as mean (min – max). N indicates the number of donors for which the protein was quantified.

|  | AD6 | BD6 | Primary Enterocytes | Jejunal mucosa |
| --- | --- | --- | --- | --- |
|  | Mean<br>(min - max)<br>N | Mean<br>(min - max)<br>N | Mean<br>(min - max)<br>N | Mean<br>(min - max)<br>N |
| SULT1A1 | 12.03<br>(11.17 - 12.61)<br>4 | 10.37<br>(6.91 - 16.03)<br>4 | 47.6<br>(44.28 - 50.91)<br>4 | 23.65<br>(22.82 - 34.1)<br>5 |
| SULT1A2 | 0.17<br>(0.1 - 0.49)<br>4 | 0.23<br>(0.17 - 0.61)<br>4 | 1.28<br>(0.6 - 1.96)<br>4 | 0.97<br>(0.59 - 1.01)<br>5 |
| SULT1A3;<br>SULT1A4 | 24.53<br>(19.05 - 39.11)<br>4 | 20.24<br>(14.8 - 23.23)<br>4 | 81.93<br>(80.87 - 82.99)<br>4 | 72.08<br>(54.98 - 77.53)<br>5 |
| SULT1B1 | 1.16<br>(0.2 - 2.76)<br>4 | 1.01<br>(0.6 - 1.84)<br>4 | 31.62<br>(25.87 - 37.36)<br>4 | 22.18<br>(20.28 - 33.25)<br>5 |
| SULT1C2 | 0.27<br>(0.18 - 0.36)<br>4 | 0.42<br>(0.06 - 1.44)<br>4 | 0.08<br>(0.08)<br>1 | 0.08<br>(0.01 - 0.13)<br>5 |
| SULT1E1 | 0.05<br>(0.04 - 0.06)<br>2 | 0.02<br>(0.02)<br>2 | 1.08<br>(0.72 - 1.43)<br>4 | 1.56<br>(0.85 - 1.7)<br>5 |
| SULT2A1 | 8.26<br>(6.43 - 10.25)<br>4 | 1.42<br>(0.56 - 9.82)<br>4 | 23.75<br>(15.52 - 31.98)<br>4 | 14.28<br>(9.28 - 18.4)<br>5 |
| SULT2B1 | 0.46<br>(0.28 - 0.64)<br>4 | 0.67<br>(0.52 - 0.89)<br>4 | 1.11<br>(0.67 - 1.54)<br>4 | 0.93<br>(0.49 - 1.03)<br>5 |

**Table S13.** Abundance of other phase II enzymes (GSTs and NATs) in human 3D enteroids, primary enterocytes, and jejunal mucosa. Protein levels are expressed in fmol/µg protein and presented as mean (min – max). N indicates the number of donors for which the protein was quantified. An asterisk (*) denotes identified proteins. ND indicates proteins that were not detected, and <LOQ refers to expression levels below the limit of quantification.

|  | AD6 | BD6 | Primary enterocytes | Jejunal mucosa |
| --- | --- | --- | --- | --- |
|  | Mean<br>(min - max)<br>N | Mean<br>(min - max)<br>N | Mean<br>(min - max)<br>N | Mean<br>(min - max)<br>N |
| GSTA1 | 55.6<br>(28.33 - 122.59)<br>4 | 31.18<br>(25.54 - 49.84)<br>4 | 99.77<br>(89.42 - 110.12)<br>4 | 116.72<br>(75.45 - 136.7)<br>5 |
| GSTA2* | <LOQ | <LOQ | <LOQ | <LOQ |
| GSTA4* | <LOQ | <LOQ | ND | <LOQ |
| GSTCD | 0.03<br>(0.02 - 0.05)<br>4 | 0.05<br>(0.03 - 0.06)<br>4 | ND | 0.01<br>(0.01)<br>2 |
| GSTK1 | 9.95<br>(9.2 - 11.75)<br>4 | 9.83<br>(9.1 - 11.23)<br>4 | 30.28<br>(29.01 - 31.54)<br>4 | 17.01<br>(16.66 - 17.39)<br>5 |
| GSTM1 | 2.73<br>(2.36 - 3.1)<br>2 | 1.3<br>(0.02 - 2.1)<br>3 | 1.91<br>(1.91)<br>1 | 2.72<br>(0.03 - 2.77)<br>5 |
| GSTM2 | 2.89<br>(1.18 - 5.91)<br>4 | 3.03<br>(1.01 - 5.02)<br>4 | 1.64<br>(0.48 - 2.8)<br>4 | 6.1<br>(3.7 - 7.16)<br>5 |
| GSTM3 | 5.84<br>(5.08 - 14.22)<br>4 | 6.4<br>(5.26 - 17.96)<br>4 | 2.9<br>(1.3 - 4.5)<br>4 | 3.3<br>(2.68 - 8.03)<br>5 |
| GSTM4 | 3.03<br>(0.58 - 7.03)<br>4 | 3.78<br>(1.3 - 6.1)<br>4 | 1.4<br>(0.08 - 2.71)<br>4 | 4.99<br>(0.49 - 5.88)<br>5 |
| GSTO1 | 19.66<br>(14.61 - 26.63)<br>4 | 17.2<br>(11.45 - 22.11)<br>4 | 25.8<br>(22.21 - 29.39)<br>4 | 25.83<br>(20.54 - 30.01)<br>5 |
| GSTP1 | 162.49<br>(126.38 - 207.09)<br>4 | 123.17<br>(99.26 - 131.25)<br>4 | 42.77<br>(40.94 - 44.59)<br>4 | 50.1<br>(47.74 - 59.18)<br>5 |
| GSTT1 | 2.16<br>(0.18 - 2.83)<br>4 | 1.65<br>(0.13 - 2.56)<br>4 | 3.48<br>(3.48)<br>1 | 2.85<br>(0.07 - 3.19)<br>5 |
| GSTT2B | 0.09<br>(0.02 - 0.2)<br>3 | 0.12<br>(0.05 - 0.18)<br>2 | ND | 0.12<br>(0.04 - 0.19)<br>2 |
| NAT1 | 1.17<br>(1.14 - 2.03)<br>4 | 0.97<br>(0.64 - 1.32)<br>4 | 1.23<br>(1.04 - 1.42)<br>4 | 0.8<br>(0.61 - 0.89)<br>5 |
| NAT2 | 0.34<br>(0.22 - 0.47)<br>4 | 0.53<br>(0.4 - 0.71)<br>4 | 0.16<br>(0.03 - 0.29)<br>4 | 0.31<br>(0.24 - 0.55)<br>5 |
| NAT10 | 0.26<br>(0.21 - 0.47)<br>4 | 0.74<br>(0.46 - 0.89)<br>4 | 0.07<br>(0.06 - 0.08)<br>4 | 0.2<br>(0.16 - 0.24)<br>5 |

**Figure S9.**
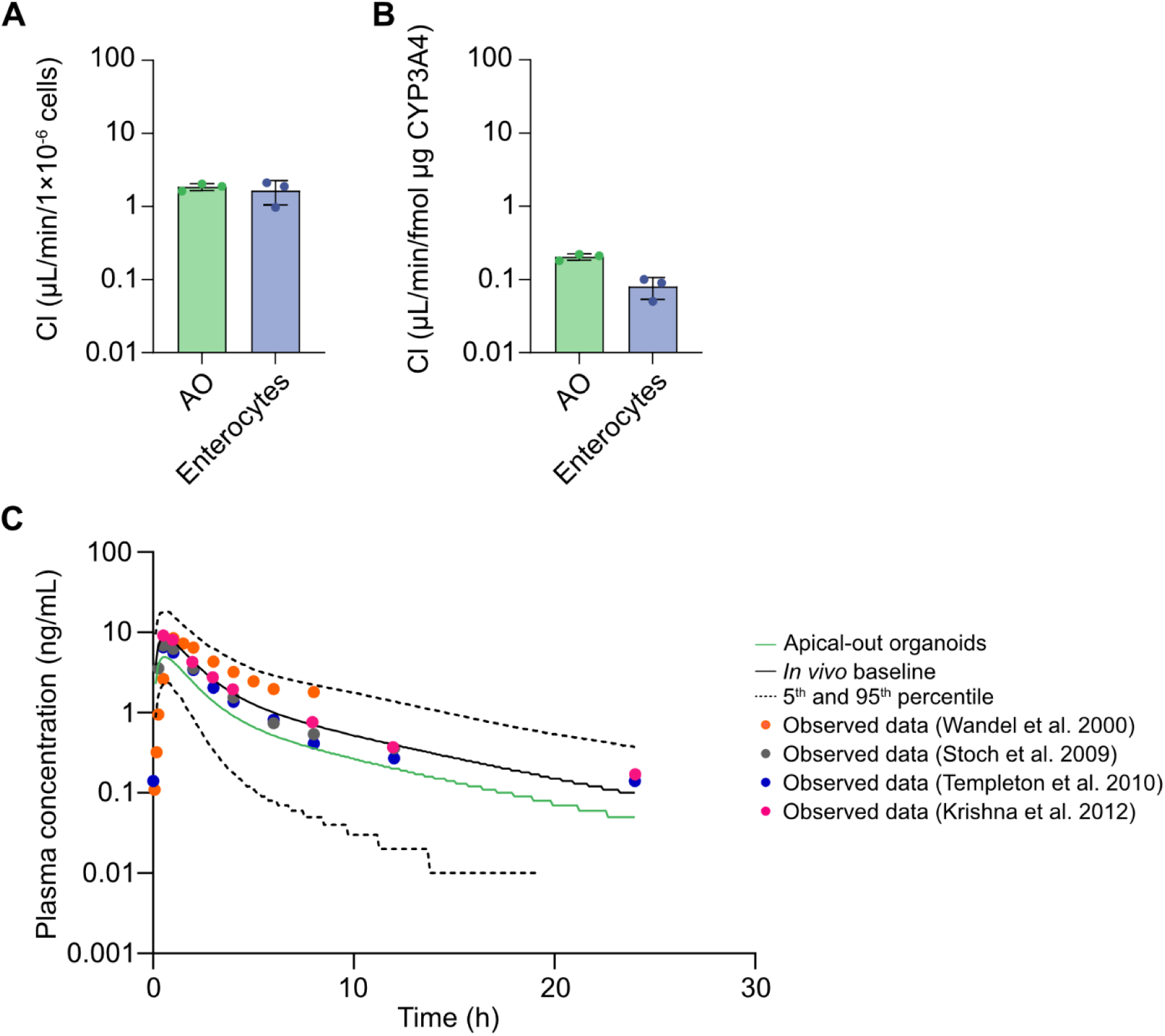
CYP3A4 metabolism of midazolam by AO enteroids and PBPK modelling of midazolam pharmacokinetics. (A) Intrinsic clearance of midazolam in apical-out enteroids and primary enterocytes. Data is shown as mean ± SD. (B) Corrected intrinsic clearance of midazolam considering the differences in CYP3A4 protein expression in apical-out enteroids and primary enterocytes. Data is shown as mean ± SD. (C) Simulated pharmacokinetics of a 5 mg dose of midazolam given to 100 virtual subjects. The green line represents the PBPK model mean prediction using data from apical-out organoids (log concentration). The black line represents model prediction *in vivo* with its 5th and 95th percentiles of the virtual population are shown as dashed lines. The colored dots depict mean observed data from the clinical studies of Wandel et al^5^, Stoch et al^6^, Templeton et al^7^ and Krishna et al.^8^

